# A Tripartite Co-culture System Reveals Defensive Mutualism Between a *Nanobdellati* Symbiont and Its Host

**DOI:** 10.64898/2026.06.04.730037

**Authors:** Hiroyuki D Sakai, Sabine Schwarzer, Satoshi Nakagawa, Milad Reyhani, Shigeru Shimamura, Matthew D Johnson, Yuya Tsukamoto, Michiru Shimizu, Ken Takai, Takuro Nunoura, Norio Kurosawa, Moriya Ohkuma, Thomas Hackl, Debnath Ghosal, Tessa Quax

## Abstract

Archaeal symbioses remain among the least understood cellular interactions. Ultra-small *Nanobdellati* (initially called DPANN) archaea rely on larger hosts for survival, yet their effects on host eco-physiology and their interactions with infecting viruses have not yet been examined experimentally. Here, we established the first stable tripartite co-culture comprising a *Nanobdellales* symbiont (YN4), its archaeal host (YN4HA), and a virus (MTIV4). We analyzed this system using physiological, genomic, transcriptomic, glycoproteomic, and cryo-electron tomography (cryoET) approaches. CryoET analysis revealed that the *Nanobdellales* archaeon formed cone-like structures to contact the host, as has been observed in other *Nanobdellati-*host systems. In this system, the host is infected by the virus. The *Nanobdellales* archaeon mitigated virus-induced growth inhibition of the host without any detectable fitness cost, indicating a defensive mutualism rather than a strictly parasitic relationship. This protective effect may involve *Nanobdellati-*driven remodeling of host cell surface glycans, suggesting a previously unrecognized glycan-mediated defense strategy. Overall, our tripartite system provides new insight into complex archaeal interactions in extreme environments that cannot be captured through conventional binary co-culture systems.

## Introduction

Archaeal symbioses, particularly those involving the diverse and evolutionarily distinct *Nanobdellati* archaea, remain among the least understood partnerships within microorganisms. The kingdom *Nanobdellati*, also referred to as the DPANN (acronym for Diapherotrites, Parvarchaeota, Aenigmarchaeota, Nanoarchaeota, Nanohaloarchaeota), forms a major evolutionary lineage that accounts for approximately half of all archaeal diversity ^1–3^. Members of this group of archaea exhibit surprising ecological diversity and can be found in various environments spanning thermophilic systems ^1,4–7^ to mesophilic habitats such as the human microbiome ^8–11^.

The DPANN superphylum, kingdom *Nanobdellati*, is characterized by nanosized cells with significantly reduced genomes, which encode limited biosynthetic and metabolic capabilities ^12,13^, suggesting a strong host dependence. Most knowledge of *Nanobdellati* originates from culture-independent methods, including genome reconstitution from metagenomes ^2,13–16^. Nearly all enriched or cultivated members known today exhibit an ectosymbiotic lifestyle that requires direct cell-to-cell contact to enable metabolic complementation and nutrient cross-feeding ^19,22^. However, the mechanisms supporting these relationships, the factors that determine host specificity, and the impact of these interactions on microbial community structure are still poorly understood.

Despite their ubiquity and presumed strong dependence on other organisms, the ecological roles of *Nanobdellati* in natural microbial ecosystems remain poorly understood, largely due to the challenges associated with their cultivation. The establishment of co-culture systems, beginning with the *Nanoarchaeum equitans*–*Ignicoccus hospitalis* model and more recently extended to additional *Nanobdellati*–host systems, has enabled direct experimental investigation of *Nanobdellati*-host interactions under defined conditions, showing parasitic or mutualistic relationships between *Nanobdellati* and host archaea ^23–32^. Recent advances in the development of controlled host-symbiont systems, combined with single-cell methods and high-resolution imaging, have provided initial insights into their cell biology ^17–21^.

The contact-dependent *Nanobdellati*-host interactions are often mediated by specialized cell surface structures that enable adhesion and the exchange of molecules ^33,34^. Direct insights into the cellular architecture of attachment are provided by cultured systems, such as *Nanoarchaeum equitans*, which forms a contact site with its host *Ignicoccus hospitalis* through fibre-like structures ^35,36^. The architecture and composition of these surface complexes can vary between systems across the kingdom. It has been shown that *Microcaldus variisymbioticus* ARM-1 interacts with its host, *Metallosphaera javensis* AS-7, via intercellular nanotubes that connect their cytoplasms ^37^. Meanwhile, the ectosymbiotic relationship between *Nanobdellales archaeon* YN1 and its host *Sulfurisphaera ohwakuensis* YN1HA is mediated through an attachment organelle that comprises a cone-like structure and an intracellular filament ^38^. The observed structural diversity of interaction types emphasizes the considerable variability of these symbiont-host associations.

Some members of the Huberarchaeota use other *Nanobdellati* lineages (e.g., Altiarchaeota) as their hosts and engage in interactions where both partners belong to the same superphylum ^39^. Others, such as *M. variisymbioticus*, can associate with multiple hosts within the *Sulfolobales*, indicating a broader host range rather than relying on a single host species or strain ^37^. Additionally, *Nanobdellati* archaea like *Nanobdella aerobiophila* can switch between free-living motile stages and contact-dependent ectobiotic phases ^40^. These dynamic life cycles suggest that the search for a host, attachment, and remodeling of its cell envelope are key elements of *Nanobdellati* physiology that facilitate metabolic exchange via specialized host-symbiont interfaces ^37,40–42^. Nevertheless, such binary systems capture only a fraction of the ecological complexity, as archaeal viruses are also widespread in natural environments. In some biological systems, the presence of a third interacting partner, including microbial, viral, insect, and plant partners, can fundamentally alter the nature of binary interactions, resulting in emergent ecological or evolutionary dynamics that cannot be predicted from pairwise studies alone ^43–48^. Indeed, it has been suggested that a member of the Candidate Phyla Radiation, an ultrasmall bacterial analogue of *Nanobdellati* archaea, protects its host bacterium from phage predation by downregulating the expression of genes involved in phage receptor biogenesis^49^. Emerging evidence suggests that viruses may act as modulators of *Nanobdellati*-host interactions, influencing host physiology, symbiont attachment, and the stability of their association ^50,51^.

Here we report the first *Nanobdellati*-host system, which is also infected with a virus. The *Nanobdellati* (YN4)-host (YN4HA)-virus (MTIV4) tripartite co-culture system was isolated from Kirishima geothermal area (Japan). The system is composed of *Nanobdellales archaeon* YN4 (Nanobdellati), *Metallosphaera hakonensis* YN4HA (host), and *Metallosphaera* Turreted Icosahedral Virus 4 (MTIV4). This work presents a comprehensive analysis of the complex interactions among these three novel players. We studied their physiology and dependence through methods such as genomics, glycoproteomics, transcriptomics, (viral) FISH, and both light and electron microscopy. Importantly, the *Nanobdellati* archaeon YN4 was found to enhance its host fitness during viral infection. This well-established system provides a detailed snapshot of the diversity and complexity of *Nanobdellati*-host-virus relationships, serving as a case study for future research into the role of viruses in interactions between archaea and their *Nanobdellati* symbionts.

## Results and Discussion

### Establishment of pure, bipartite, and tripartite co-culture systems

Novel *Nanobdellales*/DPANN host systems were isolated from samples collected in the Kirishima geothermal area, Japan. Enrichment cultures were grown under aerobic conditions at 65 °C, with the medium pH adjusted to 3.0. Microbial community analysis based on 16S rRNA gene amplicon sequencing revealed that the enrichment culture contained a previously uncultured *Nanobdellales* archaeon (4.1% abundance) alongside four other archaeal taxa: *Metallosphaera*, uncultured *Thermoplasmata* BSLdp215, uncultured *Sulfolobaceae*, and *Thermocladium* (Supplementary Information Fig. 1). After a threefold dilution-to-extinction series, a pure co-culture composed of a *Nanobdellales* archaeon (strain YN4), *Metallosphaera hakonensis* (strain YN4HA), and a virus was obtained. The presence of the virus was confirmed by genome sequencing (Supplementary Text). Negative-stain transmission electron microscopy (NS-TEM) revealed virus-like particles with a diameter of 37.6 ± 2.8 nm (n = 19) (Fig. 1), which are slightly smaller than previously reported Metallosphaera Turreted Icosahedral Viruses (MTIVs) (50-70 nm) ^53,54^. We therefore tentatively named this virus MTIV4. Starting from this tripartite culture, we subsequently established (i) a pure host culture (PURE-YN4HA), (ii) a host–virus co-culture (MTIV4–YN4HA), (iii) a *Nanobdellales*–host co-culture (YN4–YN4HA), and (iv) a reconstituted tripartite system containing the virus, DPANN, and host (MTIV4–YN4–YN4HA) (Extended Data Fig. 1; Methods; Supplementary Text).

**Figure 1.**
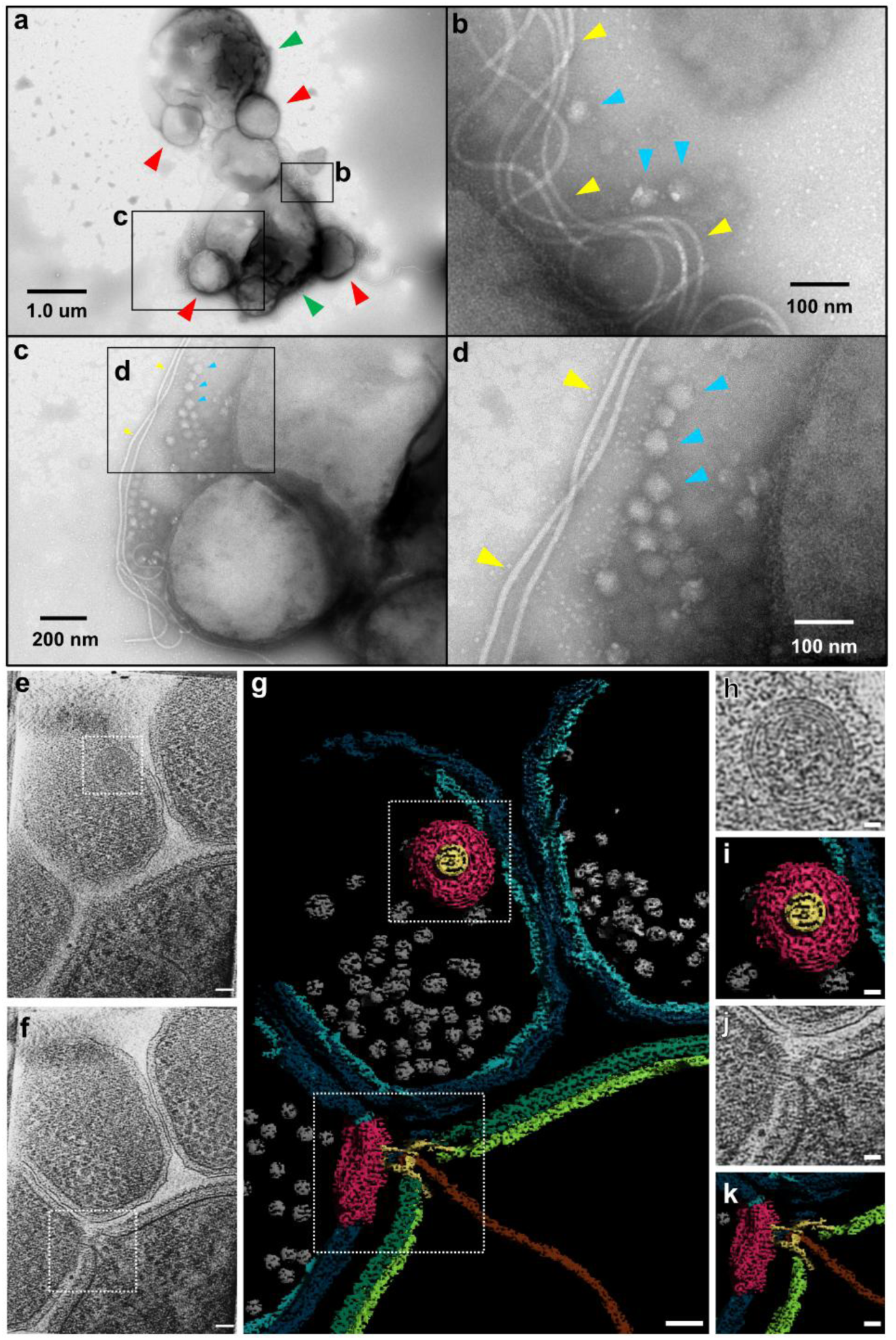
(a-d) Negative staining TEM of the YN4-YN4HA-MTIV4 co-culture. Green, red, and purple arrows indicate YN4HA cells, YN4 cells, and MTIV4 viruses, respectively. Yellow arrows indicate archaellum-like filaments. (e) Tomographic slice showing top view of an an attachment organelle. (f) Another slice of the same tomogram, showing an attachment organelle on the surface of another YN4, engaging with YN4HA. (g) 3D Segmentation of the tomogram showing YN4-YNHA interaction, attachment organelle layers and portal. (h) Zoomed in view of the attachment organelle. (i) 3D Segmentation view of the attachment organelle. (j-k) Zoomed in view of the YN4-YNHA interaction site. (j) tomographic slice, (k) 3D segmentation. Segmentation colors are host membrane green, host S-layer dark green, DPANN membrane teal, DPANN S-layer dark teal, DPANN ribosomes gray, attachment organelle layers red, attachment organelle portal yellow, and filament brown. Scale bars: (e-g) 50 nm; (h-k) 25 nm.

### Genome characteristics of YN4

The circular genome (758,205 bp) of the *Nanobdellales* archaeon YN4 encodes 837 protein-coding sequences (CDSs), 3 rRNAs, and 49 tRNAs, with a GC content of 23.4%. Of the 837 predicted CDSs, 542 were assigned to Clusters of Orthologous Groups (COG) functional categories, while the remaining CDSs lacked detectable homologs in the COG database. Of these genes, 372 (68.6%) were assigned to “Information Storage and Processing” and “Cellular Processes and Signalling” (Supplementary Information Fig. 2).

**Figure 2.**
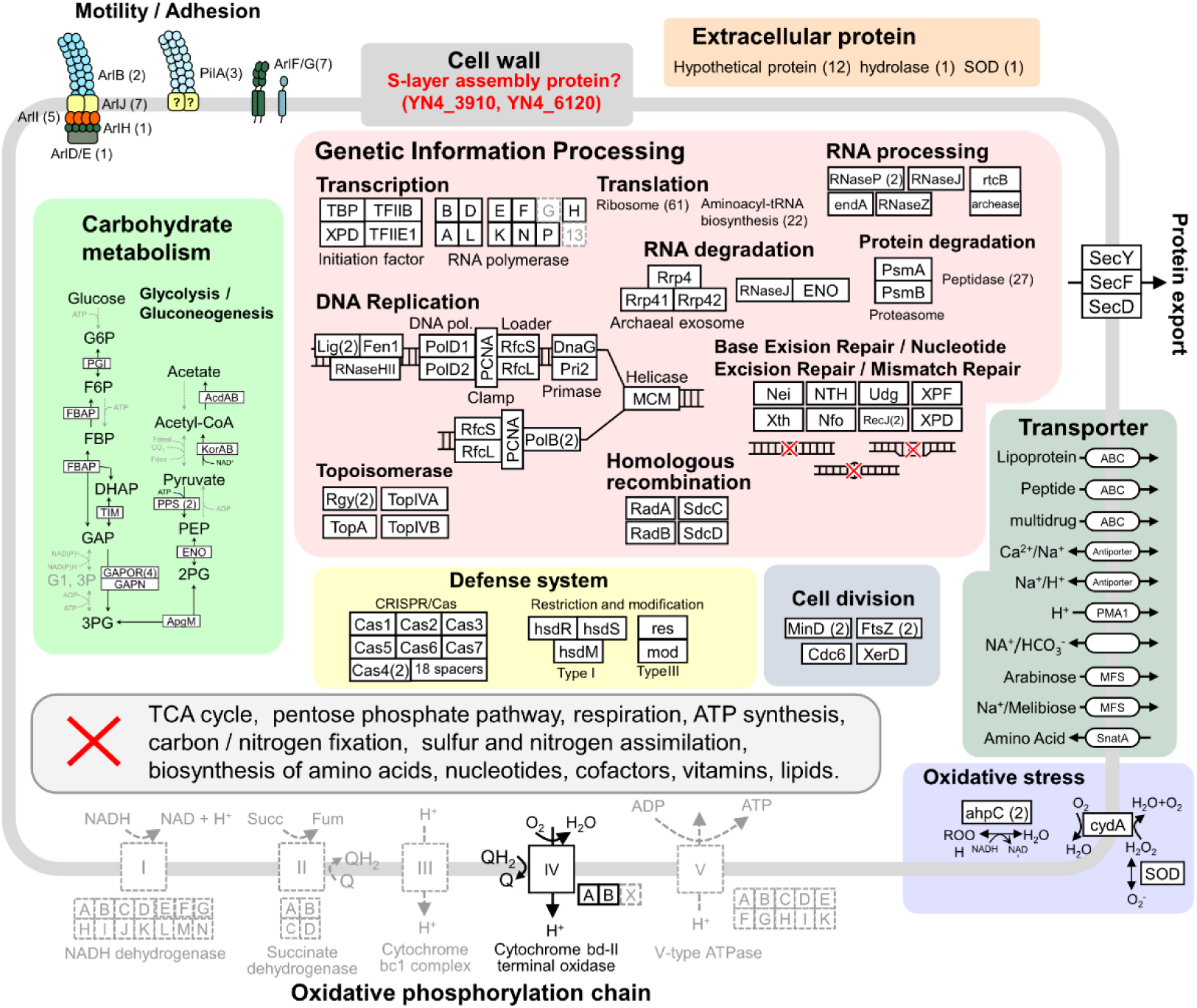
Metabolic potential of *Nanobdellales* YN4 predicted from genome annotation. Predicted metabolic and cellular functions encoded in the YN4 genome. Black and grey boxes/arrows indicate the presence and absence of genes or pathways, respectively. Numbers in parentheses represent gene copy numbers. Functional categories indicated by a red cross lack most genes required for these metabolic and biosynthetic pathways, including the TCA cycle, pentose phosphate pathway, respiration, ATP synthesis, carbon/nitrogen fixation, sulfur and nitrogen assimilation, and the biosynthesis of amino acids, nucleotides, cofactors, vitamins, and lipids.

The metabolic potential predicted from the genomic information of YN4 is shown in Fig. 2. YN4 possesses several genes involved in glycolysis and gluconeogenesis, although these pathways are incomplete. Additionally, most genes associated with central carbon metabolism pathways (i.e., the TCA cycle, pentose phosphate pathway, respiration, ATP synthesis, inorganic carbon fixation, and the biosynthesis of amino acids, nucleotides, cofactors, vitamins, and lipids) are absent from the genome (Fig. 2). Similar traits were observed among other *Nanobdellales* genomes related to YN4 (Extended Data Fig. 2). In contrast, many genes related to genetic information processing and cell division are present in the YN4 genome (Fig. 2), indicating that YN4 has an obligately symbiotic lifestyle that requires nutrients/energy from other community members, while it retains the genetic potential for independent cell proliferation.

### Genome characteristics of YN4HA

The circular genome (2,631,175 bp) of the *Metallosphaera hakonensis* host YN4HA encodes 2,817 coding sequences, 2 rRNAs, and 46 tRNAs, and has a GC content of 43.4%. The tricarboxylic acid cycle, gluconeogenesis, and semi- and non-phosphorylative Entner–Doudoroff pathways were all detected (Supplementary Table 1). Two archaeal carbon fixation pathways, the hydroxypropionate–hydroxybutyrate and dicarboxylate–hydroxybutyrate cycles ^55,56^, were also detected. Genes associated with sulfur reduction (CoADR) and tetrathionate/thiosulfate oxidation (doxAD) were present (Supplementary Table 1). These features indicate that YN4HA has the metabolic capacity for a free-living lifestyle. At present, no metabolic pathways appear to be complemented by the symbiont YN4.

### Genome characteristics of MTIV4

The genome of the virus detected in the tripartite *Metallosphaera hakonensis* host YN4HA, *Nanobdellales* YN4 culture, MTIV4, is 13,941 bp long, linear, with inverted terminal repeats and encodes 30 predicted protein-coding genes (Fig. 3a). The 109-bp, GA-rich terminal repeats are similar but shorter than those of the other MTIV viruses (160-353 bp) and comprise a distinct terminal motif ([GA]^5^GGAT[G]^11^). In terms of length and gene count, MTIV4 is the largest representative of this viral group, with the lowest GC content (45% compared with 51-53% for other members of the group), and, based on sequence similarity and core-protein-based phylogeny, is more closely related to MTIV3 than to MTIV1/2 (Fig. 3a). Out of the 30 protein-coding genes, only five have predicted functions (MTIV4_6: transcriptional repressor; MTIV4_[15-17, 21]: virion structure).

**Figure 3.**
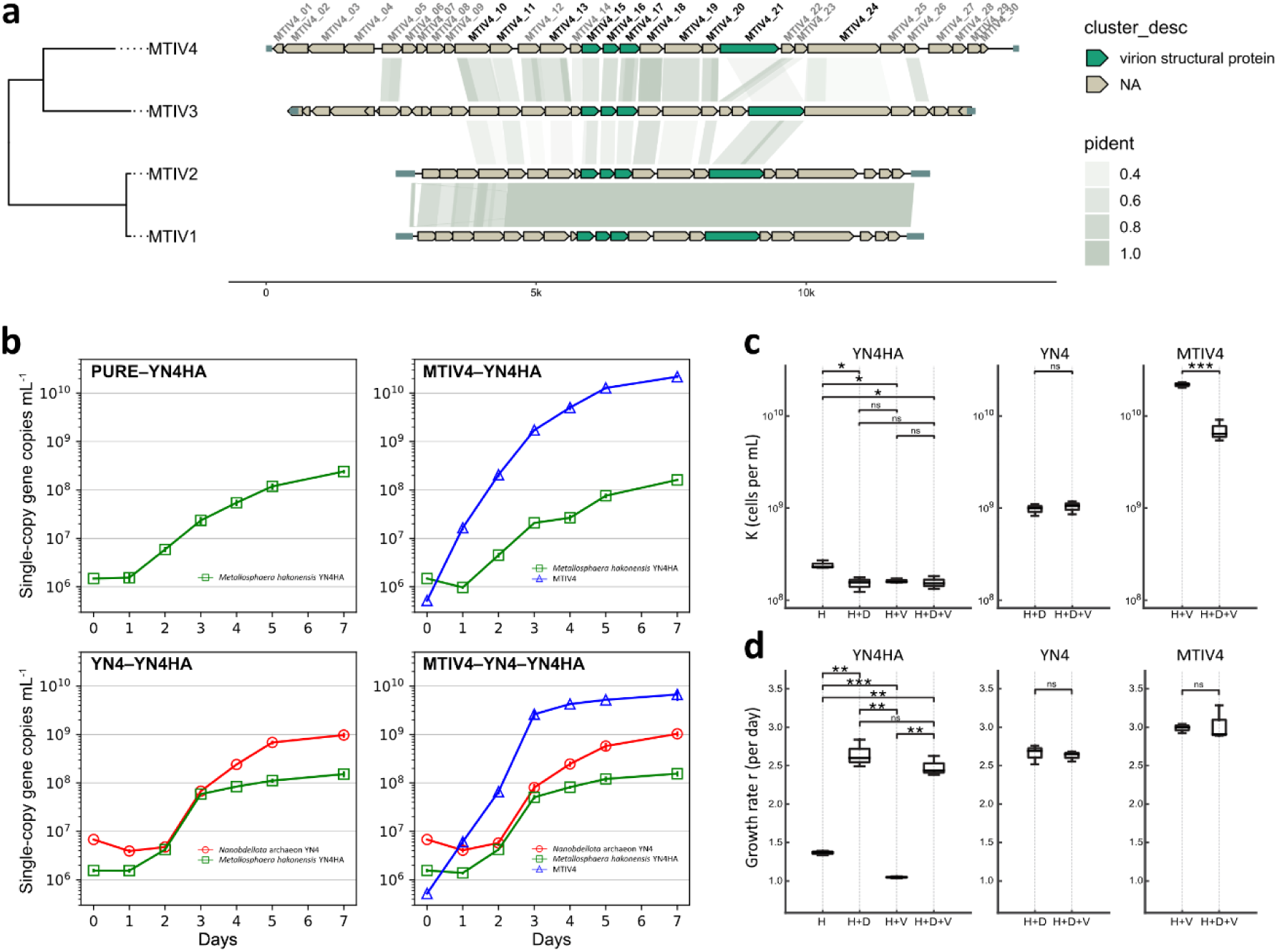
Genomic characteristics of MTIV4 and its biological interactions with the YN4–YN4HA system. (a) Comparison of MTIV4 genome architecture with previously described MTIV viruses. Evolutionary relationships among the four viruses are represented by the phylogenetic tree on the left reconstructed from eleven concatenated core protein alignments (dark black tags). Viral genes are indicated as arrows, with predicted virion structural genes highlighted in green. Shaded areas between genomes indicate regions of detectable amino-acid-level sequence similarity. (b) Growth curves of pure- and co-cultures of strain YN4 and strain YN4HA with or without MTIV4. Growth was measured by qPCR. (c, d) Carrying capacities and growth rates of YN4HA host (left), Nanobdellota YN4 (center), and MTIV4 virus (right) under different conditions. The four culture conditions are: host alone (YN4HA) (H), host + Nanobdellota (YN4HA + YN4) (H+D), host + virus (YN4HA + MTIV4) (H+V), and host + Nanobdellota + virus (YN4HA + YN4 + MTIV4) (H+D+V), Pairwise differences between conditions were evaluated using Welch’s t-tests; significance levels are indicated by asterisks above the brackets (*, P < 0.05; **, P < 0.01; ***, P < 0.001; ns, not significant).

### Growth analyses reveal YN4-driven defensive mutualism in the tripartite system

Growth curves (quantified by qPCR targeting the archaeal 16S rRNA gene and a viral single-copy conserved hypothetical protein gene shared among MTIV1–MTIV4, respectively) of the pure and co-culture systems are shown in Fig. 3b, and the estimated carrying capacities (K; maximum population size supported under the cultivation conditions) and specific growth rates (r) are shown in Fig. 3c and 3d.

When the YN4 symbiont was present with the host (H+D), the host’s K decreased by ∼37% (Fig. 3c), indicating that strain YN4 imposed a measurable competitive cost, consistent with resource consumption as suggested by previously characterized co-culture systems ^23,30,31^. The prolonged lag phase of YN4HA observed in the YN4–YN4HA growth curve (Fig. 3b) suggests that YN4 imposes early physiological stress on the host. While this early effect does not directly explain the reduction in carrying capacity, it indicates that YN4 influences host physiology across multiple growth phases, which may contribute to the overall decrease in K.

The presence of the MTIV4 virus also led to a similar ∼33% reduction in host K (H+V) (Fig. 3c). The early decline in host abundance (day 1) and the slight growth inhibition (day 3 to 4) observed in the MTIV4–YN4HA system (Fig. 3b) indicate that the virus negatively affects host growth during the initial and exponential phases, which could account for the reduced K. When both YN4 and MTIV4 were present with the host (H+D+V), the virus did not cause any further decrease in the host’s K beyond that imposed by YN4 alone, whereas the virus’s own K decreased by ∼68% (Fig. 3c), as estimated from logistic fits to the growth curves shown in Fig. 3b. Indeed, the host’s growth dynamics in the tripartite co-culture closely resembled those in the virus-free bipartite co-culture (Fig. 3b). These observations suggest that YN4 shields the host population from additional virus-induced reductions in carrying capacity while substantially suppressing viral abundance by ∼68%, thereby mitigating the impact of MTIV4 on host growth.

Specific growth rates (r) further clarified these interaction patterns (Fig. 3d). Although YN4HA showed a longer lag phase when YN4 was present, the symbiont enhanced the host’s growth rate in the co-culture (H+D) (Fig. 3b, d). To evaluate how effectively YN4 restores host growth suppressed by viral infection, a recovery index was calculated as follows: Recovery (%) = [(r_H+D+V_ − r_H+V_) / (r_H_ − r_H+V_)] × 100, where 100% is defined as the virus-induced reduction in growth rate between the host-only (H) and virus-infected (H+V) conditions. Using growth rates (r) derived from exponential fits to the growth curves, this calculation yielded (2.48 – 1.05) / (1.37 – 1.05) × 100 ≈ 447%. Thus, YN4 increased the host’s growth rate by approximately 4.5 times the magnitude of the viral suppression, demonstrating that YN4 not only compensated for the virus-induced inhibition but also boosted host growth beyond its virus-free baseline. Although host growth was suppressed in the presence of MTIV4, the presence of YN4 strongly counteracted the inhibitory effect of MTIV4 infection in the tripartite co-culture (H+D+V; Fig. 3d). This may also contribute to YN4’s efficient nutrient acquisition by enhancing the host’s growth, thereby improving the ability to acquire or utilize nutrients from the surrounding environment. Additionally, YN4 may provide metabolic products that could benefit the host’s growth. Another possibility is that the initial stress caused by YN4 might trigger adaptive responses in YN4HA, leading to more efficient growth once it is adapted. The molecular mechanisms underlying this phenomenon remain to be elucidated.

Despite these benefits to the host, YN4 itself was not significantly affected by the presence of MTIV4, as both its carrying capacity and growth rate remained unchanged between the H+D and H+D+V conditions (Fig. 3c,d). The absence of any measurable disadvantage for YN4 supports the interpretation that the interaction represents defensive mutualism rather than strict parasitism.

### Light microscopy and fluorescent microscopy (FISH) reveal enlargement of host cells in the presence of *Nanobdellales* archaeon

Co-cultures of YN4HA and YN4 were analyzed by light microscopy and FISH analysis (Fig. 4). Under phase-contrast and differential interference contrast microscopy, larger cells (>1 μm) and smaller cells (<500 nm) were clearly observed in the YN4-YN4HA co-culture at 1000× magnification (Fig. 4a, C; Supplementary Movie S1). The smaller cells were barely visible at magnifications of 400× or lower. FISH analysis confirmed that the smaller and larger cells correspond to strains YN4 and YN4HA, respectively (Fig. 4d). The smaller cells were not observed in the pure YN4HA culture, where cell sizes typically ranged from 1 to 2 μm (Fig. 4e; Supplementary Movie S2). These results indicate that the smaller cells observed under light microscopy are indeed strain YN4, rather than cell debris or membrane vesicles originating from YN4HA.

**Figure 4.**
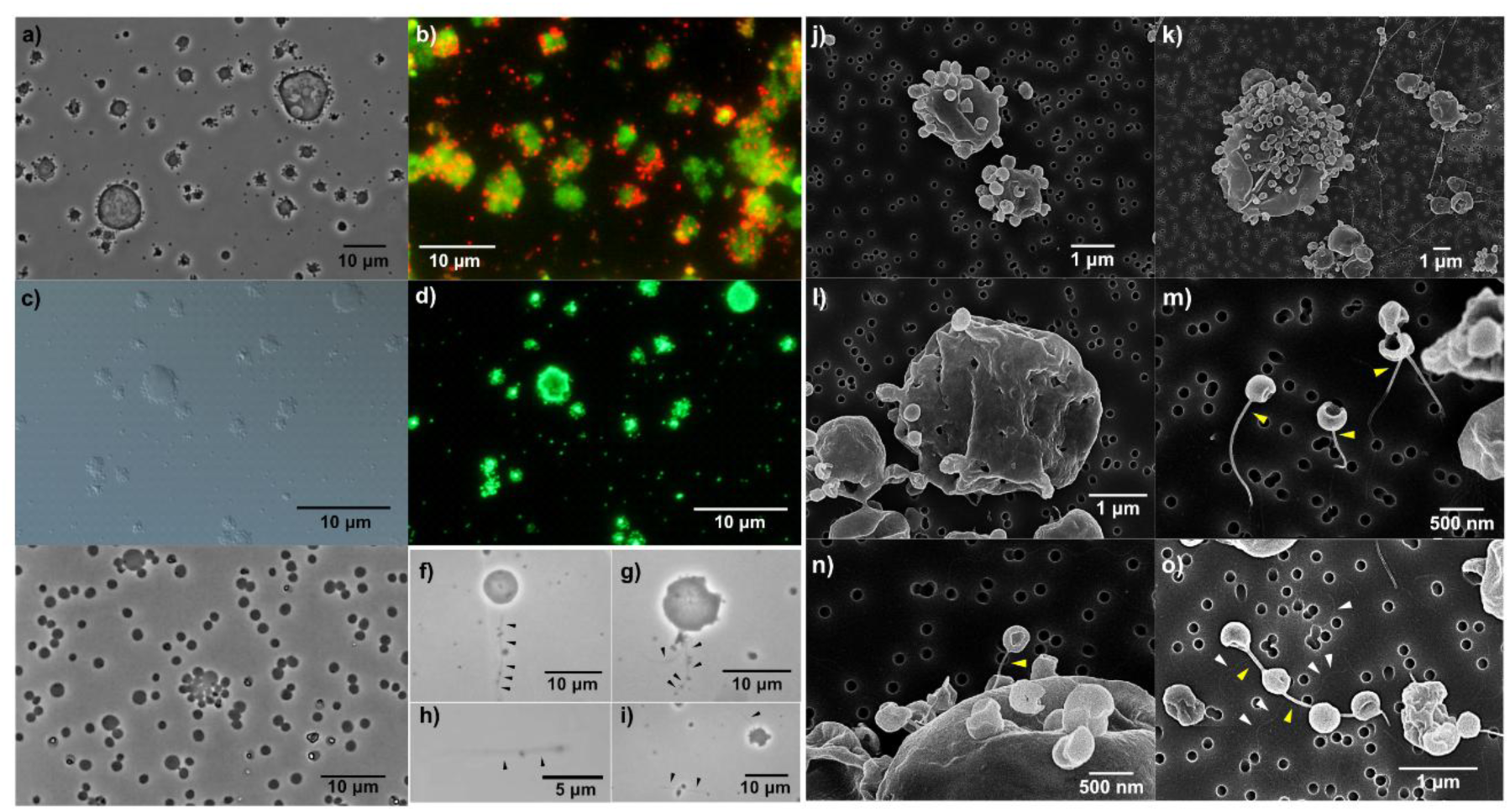
Scanning electron microscopy and light microscopy of YN4-YN4HA co-culture and YN4HA pure culture. (a) Phase-contrast microscopic image of YN4-YN4HA co-culture. (b) Differential interference contrast microscopic and (c) fluorescent microscopic images of YN4/YN4HA cells stained with SYBR green I in the same view area. (d) FISH image of YN4 (red) and YN4HA (green) cells. (e) Phase-contrast microscopic image of YN4HA pure culture. (f-i) Phase-contrast microscopic images of YN4-YN4HA co-culture showing nanotube-like structures (black arrows) similar to those observed under the scanning electron microscope (Extended Data Figs. 3 and 4). (j) Scanning electron microscopy images of YN4 cells attaching/surrounding to a YN4HA cell, (k) an enlarged YN4HA cell surrounded by YN4 cells, (l) an enlarged YN4HA cell having holes on its cell surface, (m) nanotubes emerged from YN4 cells, (n) a YN4 cell attaching to a YN1HA cell with nanotube, (o) a YN4 cell interconnected by a nanotube. White arrow heads: archaellum-like filaments, yellow arrow heads: nanotubes.

In the YN4-YN4HA co-culture, YN4HA cells were often enlarged compared to cells of this strain in the pure YN4HA culture (Fig. 4 a,e). In addition, they were surrounded by numerous YN4 cells (Fig. 4 a,d,k). Such enlargement was rarely observed in the pure YN4HA culture (Fig. 4e), indicating that this enlargement is induced by the attachment of YN4 cells to YN4HA cells. Moreover, the number of YN4 cells attached to the enlarged YN4HA cells was considerably higher compared to YN4 cells that were attached to normal-sized YN4HA cells. Given that enlarged host cells may increase the likelihood of YN4 recognizing and attaching to the host, this phenomenon could represent an important survival strategy for YN4. The enlarged YN4HA cells may serve as growth compartments for the YN4 symbiont.

Time lapse imaging under near-native growth conditions (65 °C) in phase contrast microscopy, revealed that the *Nanobdellales* YN4 is very motile. Numerous swimming cells with a diameter of ∼0.5 μm, corresponding to YN4, were observed in time-lapse movies (Supplementary Movie S3). The YN4HA host was not observed to display swimming motility under the current culturing conditions. Interestingly, the *Nanobdellales* YN4 was observed both in a free state during which it was mainly swimming, or non-motile attached to host YN4HA cells, which shows that YN4 displays two distinct lifestyles.

### Electron microscopy reveals tube-like structures and cones that connect host and *Nanobdellales* archaeon YN4

Scanning electron microscopy was performed on the tripartite cultures containing YN4HA, YN4 and MTIV4. Filamentous structures, approximately 20 nm in diameter, were observed (Fig. 4o). Similar structures are also present in the NS-TEM images shown in Fig 1, where these measure ∼10 nm. Given that a plasma osmium coating (5 s) could produce a surface layer of a few nanometres, the 20 nm-wide filamentous structures should correspond to the same filaments observed under NS-TEM. A typical archaellum is known to have a diameter of 10–14 nm; therefore, we assume the observed filaments to be archaella, which would contribute to the swimming capability of YN4 observed by live-cell imaging of the YN4-YN4HA co-culture, as described above (Supplementary Movie S3).

Nanotube-like structures, such as those reported in the *Thermococcus* archaeon (∼60–80 nm in diameter and often >1 µm in length) ^57^, were occasionally observed under SEM (Fig. 4m,n,o, Extended Data Fig. 3 and 4). The YN4 nanotubes, with diameters of approximately 30–50 nm and lengths ranging from 0.2 to 3 µm, were larger than the archaella, suggesting that the tubes have a different function from the archaellum-like filaments. Similar structures were also observed under phase-contrast microscopy (Fig. 4f,g,h,i). Three types of nanotube states were observed under SEM: (1) protruding from a YN4 cell without attaching to any host cell (Fig. 4m, Extended Data Fig. 3a,b); (2) connecting several YN4 cells (Fig. 4o, Extended Data Fig. 3c,d,e,f); and (3) attaching to a host cell (Fig. 4n, Extended Data Fig. 4). These observations suggest that the nanotubes have adhesive functions, such as gathering YN4 cells, recognizing host cells, or perhaps contributing to biofilm formation as well. The YN4 genome encodes multiple genes related to adhesive type IV pili (e.g., YN4_047, YN4_417 and YN4_710; Supplementary Table 2). Type IV pili are typically ∼4-6 nm in diameter ^58^, and thus these genes are likely not encoding the nanotubes, which might be encoded by still unknown genes.

As observed by light microscopy, enlarged host cells, >5-10 μm in diameter, surrounded by YN4 cells were also observed under SEM (Fig. 4k, Extended Data Fig. 5). A maximum of 93 YN4 cells were observed attaching to the visible side of a YN4HA cell (Fig. 4k). Considering the unseen side of the YN4HA cell, it is estimated that a total of ∼186 YN4 cells were attached to the enlarged YN4HA cell. The large size of the host cells may also increase their metabolic capacity, potentially supporting a greater number of attached YN4 cells through enhanced nutrient production. This interpretation is consistent with the upregulation of metabolic genes observed in the transcriptomic analysis described below, although further experiments will be needed to test this possibility. We also found that some of the enlarged cells had several holes on their cell surface (Fig. 4l, Extended Data Fig. 6a,b,c,d), which appear to be remnants of YN4 cells that had detached in the process of preparing the sample for imaging.

Another intriguing structure observed under SEM is a spider-web-like network (Extended Data Fig. 6e,f,g,h). This web-like structure comprises fibres of varying thickness, distinguishing it from the uniform diameters of archaellum-like filaments and nanotubes described above. Based on its morphology, we hypothesize that this structure consists of extracellular polymeric substance (EPS). Similar branched, web-like EPS strands have been reported in SEM analyses of hydrated biofilm matrices, where EPS collapses into filamentous, web-like structures during sample dehydration ^59^, suggesting that our spider-web-like structure may likewise represent a dehydration-induced artefact of collapsed EPS. Such EPS networks are commonly associated with biofilm formation in many microorganisms. Indeed, during cultivation of the YN4–YN4HA co-culture, visible precipitates consistently formed—a phenomenon not often observed in the pure YN4HA culture —resulting in significantly lower turbidity in the upper portion of the culture tube (Supplementary Information Fig. 3). Gentle mixing readily dispersed these precipitates into small flocs, thereby increasing the turbidity of the co-culture. These observations suggest that YN4 induces the formation of substantial cell aggregates, flocs or biofilm-like structures in co-culture. Although it remains unknown whether the EPS originates from YN4 or YN4HA, the EPS-like structures may contribute to biofilm formation and potentially facilitate interactions between YN4 and YN4HA cells.

Under NS-TEM, a cone-shaped structure, recently discovered in other *Nanobdellota* species ^32,60^, was also observed on the surface of YN4 cells (Extended Data Fig. 7). Three states of this cone were identified: (1) a cone on the surface of a YN4 cell, with its vertex likely intact and lacking an opening (Extended Data Fig. 7a,b); (2) a cone interacting with or attached to a YN4HA cell (Extended Data Fig. 7c,d); and (3) a cone with a short tube-like structure at its vertex opening—likely corresponding to the “portal” reported in other *Nanobdellota* archaea ^32,60^ (Extended Data Fig. 7e,f,g,h). Additionally, remnants of the S-layer-like structure from YN4HA were observed near the tip of this short tube (Extended Data Fig. 7f). These observations may reflect different stages in the lifecycle of YN4: (1) a preparatory phase in which the cone structure forms before attachment to a host; (2) an interaction phase in which the cone with the portal is used to obtain nutrients from the host cell; and (3) a post-interaction phase after acquiring the necessary nutrients from the host. The presence of S-layer-like remnants near the cone of YN4 suggests that the host YN4HA to which the *Nanobdellales* YN4 was attached had eventually decomposed. Since the portal vertex remained open, YN4 cells with an open short portal that are no longer attached to a host cell are likely dead. Without closing the portal, the cytoplasm of YN4 remains exposed to the external environment. Indeed, multiple membrane vesicle-like structures were observed on the outer cell wall of YN4 (Extended Data Fig. 7e,g), suggesting that the cell is undergoing degradation, although the structure might be an artifact of sample preparation.

Furthermore, cryo-electron tomography (cryoET) provided higher-resolution insights into the YN4–YN4HA attachment interface (Figure 1e-k, Supplementary Information Fig. 4, Supplementary Movie S4). In our tomograms, we observed both isolated YN4 cells and YN4 cells attached to their hosts. The diameter of YN4 cells were approximately 500–750 nm, whereas YN4HA cells were about 1–2 µm in diameter. On average, we observed two YN4 cells near YN4HA cells (n = 34, min = 0, max = 6).

Some YN4 cells established a connection with the host through a cone-like structure which we define as as an “attachment organelle” resembling that previously reported in other *Nanobdellota* ^32,60,61^. This organelle consists of multiple layers of a putatively repeating monomer stacked on top of one another, forming a cone-shaped structure with its wide opening facing the YN4 cytosol. At its apex, a portal-like structure protrudes toward the outside of the cell, from which a filament occasionally extends into the YN4HA cell by displacing the host S-layer and establishing contact with the host cell membrane. We observed a maximum of one attachment organelle per YN4 cell. Some cells lacked the organelle entirely, others possessed a single organelle that was not engaged in host attachment, while others used the organelle to establish contact with the host. This variation is consistent with the distinct cone states identified by NS-TEM analysis described above. The attachment organelle complex had an average base diameter of 221.1 nm (SD = 53.3 nm, n = 34), an average height of 59.9 nm (SD = 7.3 nm, n = 17), an average portal height of 20.8 nm (SD = 3.2 nm, n = 17), an average portal diameter of 24.6 nm (SD = 2.8 nm, n = 17), and the associated filament had an average diameter of 8.2 nm (SD = 2.3 nm, n = 11). These cryo-ET observations support the idea that the attachment organelle mediates direct physical interaction between YN4 and its host ^32,60,61^.

Thus, electron microscopy revealed the presence of several distinct cell surface structure that are involved in *Nanobdellota*-host interaction: (i) archaella used by *Nanobdellota* to swim to new hosts (ii) nanotubes used by *Nanobdellota* to extend from the cell and attach to host cells, (iii) cones formed by *Nanobdellota* to create direct contact with the host cell and (likely) continuous cytoplasmic bridge, (iv) spider web like structures, likely consisting of EPS that are part of a biofilm matrix made by the *Nanobdellota* and/or their host.

### Structural characterization of N-glycans and their remodeling in co-culture systems

Archaeal cell surfaces are commonly modified with N-glycans ^62^. In order to gain deeper insight into the impact of *Nanobdellota* YN4 presence on the YN4HA host, we performed structural characterization of N-glycans of both organisms. Given the efficiency of focused searches in detecting glycopeptides and glycoproteins ^63^, we conducted this analysis by targeting the predominant modification masses (Supplementary Information Fig. 5) for each strain. The glyco-PSM analysis revealed that many protein modifications in strain YN4HA were primarily associated with S-layer protein A, both in pure culture and in tripartite co-culture with strain YN4 and MTIV4, consistent with those previously reported in other *Thermoproteota* archaea (Supplementary Tables 3 and 4) ^64^. In strain YN4, hypothetical proteins (YN4_326 and YN4_730) and archaellin ArlB (YN4_015), the second most highly expressed gene in the YN4 transcriptome, were identified as glycoproteins (Supplementary Table 5). Archaellin glycosylation was previously reported in *N. aerobiophila* MJ1^T 64^, indicating a common and important function, given the streamlined genome and the energetic cost of glycosylation.

**Figure 5.**
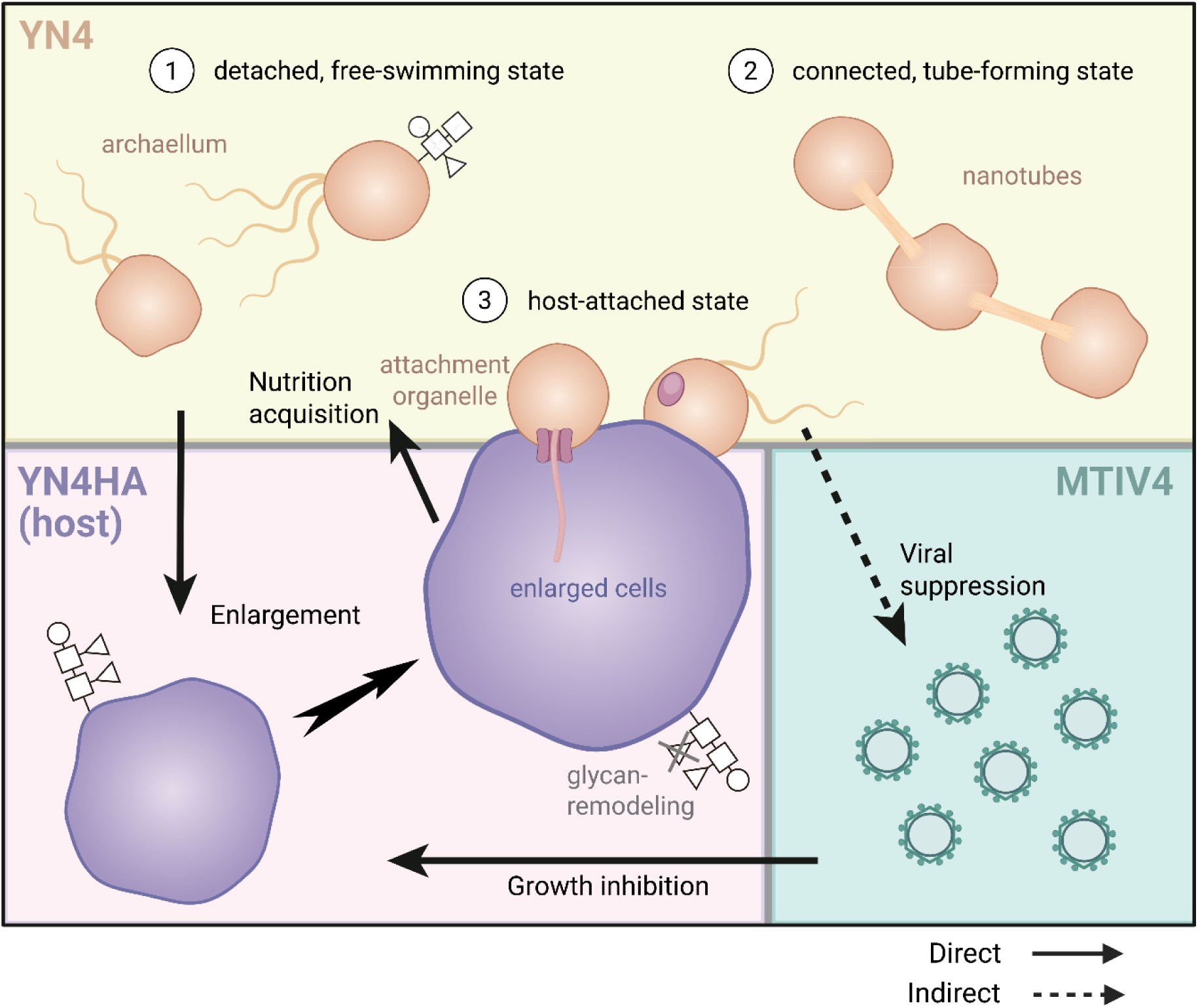
Proposed model of *Nanobdellales*-host-virus interactions. The *Nanobdellales* archaeon YN4 exhibits three distinct life cycle stages: a detached, free-living, motile state; a connected state that forms nanotubes or spider-web-like structures; and a host-attached state, during which it builds attachment organelles to connect to its host, *Metallosphaera hakonensis* YN4HA. The tripartite system comprises the *Nanobdellales* archaeon YN4, its host *M. hakonensis* YN4HA, and the virus MTIV4 that infects and inhibits YN4HA. In this relationship, MTIV4 and YN4 likely compete for host-derived resources. The presence of YN4 has a protective effect, potencially involving glycan remodeling of the host cell surface and reducing viral proliferation. In return, the *Nanobdellati* symbiont receives nutrients from its hosts. YN4HA host cells harboring YN4 are significantly enlarged.

The MS/MS spectra of glycopeptides from strain YN4 consistently identified potential sugar oxonium ions at 204.087 Da, indicative of HexNAc. The MS/MS analysis of peptides modified by 1200.43 Da revealed a glycan likely comprising HexNAc_2_Hex_4_dHex (theoretical mass: 1200.427947) (Extended Data Fig. 8a). This structure is likely one dHex larger than the glycan reported in *N. aerobiophila* MJ1^T 64^. Given these data, modifications of 1054.37, 1038.38, and 892.32 Da were likely HexNAc_2_Hex_4_ (theoretical mass: 1054.370039), HexNAc_2_Hex_3_dHex (theoretical mass: 1038.375124), and HexNAc_2_Hex_3_ (theoretical mass: 892.3172152), respectively. Archaeal protein *N*-glycosylation is known for its diverse array of linking sugars ^62^. HexNAc as a linking sugar is common among previously characterized glycoproteins in *N. aerobiophila* MJ1^T 64^.

MS/MS analysis of the glycopeptides with a 1305.42 Da modification from the pure culture of strain YN4HA revealed a glycan composed of HexNAc_3_Hex_2_SdHexdHex (theoretical mass: 1305.416397) (Extended Data Fig. 8b). Additionally, the 568.21 Da modification was associated with HexNAc_2_Hex (theoretical mass: 568.2115684). In the co-culture, the 1159.36 and 794.23 Da modifications were identified as HexNAc_3_Hex_2_SdHex (theoretical mass: 1159.358488) and HexNAc_2_HexSdHex (theoretical mass: 794.226292) (Extended Data Fig. 8c), indicating the loss of a dHex residue in the *N*-glycan of strain YN4HA during co-culture with strain YN4. Previously, we reported the loss of a pentose residue in the host archaeon co-cultured with *Nanobdella* ^64^. This glycan remodeling may represent a common phenomenon associated with host-DPANN interactions. Because cell-surface glycans are known to influence interactions between microorganisms and viruses in various systems, the observed loss of specific glycan residues may alter host cell surface properties and thereby influence host–virus interactions. If this remodeling represents a common feature of host–DPANN interactions, such modifications may have broader ecological implications in natural environments. By influencing host–virus interactions, DPANN symbionts could potentially modulate viral susceptibility of their hosts and thereby affect microbial population dynamics. However, further experimental studies will be required to determine whether glycan remodeling directly affects viral infection processes in this system.

### Differential gene expression analysis of YN4HA with or without YN4

In order to gain insight into the impact of *Nanobdellota* presence in YN4HA host cultures, RNA sequencing was used to compare the mRNA levels between pure YN4HA cultures and YN4HA –YN4 co-cultures. A total of 281 genes were significantly upregulated (padj < 0.01, log2 fold-change ≥ 1) in YN4HA during co-culture with YN4 (Supplementary Table 6, Supplementary Information Fig. 6). Of these, 154 were assigned to COG categories, while the remaining genes had no detectable homologs in the COG database (Supplementary Information Fig. 7c). The most abundant COG category among the upregulated genes was energy production and conversion (C; 34 genes), followed by mobilome (X; 18), general function prediction only (R; 17), transcription (K; 12), and defense mechanisms (V; 9). In contrast, 112 genes were significantly downregulated (padj < 0.01, log2 fold-change ≤ –1) (Supplementary Table 7, Supplementary Fig. 5c). Of these, 67 were assigned to COG categories. The most abundant COG category among the downregulated genes was amino acid transport and metabolism (E; 9 genes), followed by translation, ribosomal structure and biogenesis (J; 8), replication, recombination and repair (L; 8), mobilome (X; 7), and general function prediction only (R; 3).

### Differential gene expression analysis of YN4HA with or without MTIV4

A total of 11 genes were significantly upregulated in *Metallosphaera hakonensis* YN4HA upon infection with MTIV4 (padj < 0.01, log2 fold-change ≥ 1) (Supplementary Table 8, Supplementary Information Fig. 8). Of these, 7 were assigned to COG functional categories (Supplementary Information Fig. 7d). The identified categories included replication, recombination and repair (L; 1 gene), coenzyme transport and metabolism (H; 1), energy production and conversion (C; 1), cell cycle control (D; 1), mobilome (X; 1), general function prediction only (R; 1), and function unknown (S; 1). In contrast, 53 genes were significantly downregulated (padj < 0.01, log2 fold-change ≤ –1) (Supplementary Table 9, Supplementary Information Fig. 8), of which only 13 were assigned to COG categories (Supplementary Information Fig. 7d). The most represented categories among these were replication, recombination and repair (L; 3 genes) and mobilome (X; 3). Thus, the overall observed response of YN4HA to MTIV infection was relatively mild with a modest number of differentially regulated genes.

### Distinct transcriptional responses to *Nanobdellati* symbiosis and viral infection

Host transcriptomic responses differed markedly between *Nanobdellati* symbiosis and MTIV4 infection, with *Nanobdellati* inducing extensive transcriptional reprogramming, compared to the response to MTIV4 infection.

In the presence of YN4, the transcriptomic changes observed in YN4HA indicate a substantial physiological response to *Nanobdellati* symbiosis. In particular, the significant upregulation of genes related to energy production and conversion suggests that YN4HA increases its metabolic output, possibly to meet the energetic demands of supporting YN4 growth. This metabolic shift may contribute to the accelerated growth of YN4HA observed in the co-culture condition (Fig. 3b), potentially reflecting increased energy expenditure in response to nutrient or energy withdrawal by the *Nanobdellati* symbiont.

A similar but relatively limited host response has been reported for the *Ignicoccus hospitalis*–*Nanoarchaeum equitans* co-culture system ^65,66^. In that system, increased energy metabolism and transporter activity were observed, together with downregulation of genes associated with transcription, replication, and cell-cycle control, while stress- and defense-related pathways, including the CRISPR–Cas system, remained relatively unchanged ^66^. Thus, although both systems commonly exhibit enhanced energy metabolism during *Nanobdellati* symbiosis, YN4HA additionally shows strong upregulation of defense- and mobilome-related genes, indicating a more extensive host response in this *Nanobdellati*-host co-culture system.

In a metagenomic study, Esser et al. ^67^ reported that CRISPR spacers in the genomes of “*Candidatus* Altiarchaeum” species matched chromosomal regions of their *Nanobdellati* episymbionts, providing evidence that host CRISPR–Cas systems can target episymbiont chromosomal DNA. However, no CRISPR spacer sequences in the host genome matched the YN4 genome, indicating that direct CRISPR targeting of the symbiont is unlikely to occur in this system.

The upregulation of CRISPR-related genes observed in our experiments could be linked with a more general activation of host defense mechanisms that are possibly induced by *Nanobdellati* symbiosis. Comparable transcriptional responses have been reported in model archaeal systems during virus-host interactions. For example, infections of *Sulfolobus* species with archaeal viruses such as SIRV2 and STSV2 strongly activate host defense systems, including CRISPR–Cas and toxin-antitoxin modules, and increase expression of transposase and other mobilome-related genes ^68,69^. These parallels suggest that the concurrent upregulation of mobilome and defense genes in YN4HA may reflect a broadly conserved archaeal stress response to biological antagonists, regardless of whether they are viral or archaeal in origin.

Notably, genes strongly induced during *Nanobdellati* co-culture—such as those involved in energy production, mobilome activation, and defense mechanisms—remained largely unchanged during the MTIV4 infection (Supplementary Information Fig. 7c,d), indicating that the host response to MTIV4 is considerably weaker than that induced by *Nanobdellati* symbiosis. The limited effect on host gene expression contrasts with the lytic virus SIRV2, which strongly activates host defense genes, including CRISPR–Cas and toxin–antitoxin systems ^68^. By contrast, the archaeal chronic virus SMV1 establishes a long-term non-lytic infection without triggering significant activation of host defense systems ^70^. Although MTIV4 negatively affects host growth (Fig. 3b), its weak induction of defense- and mobilome-related genes more closely resembles the response to chronic viruses than to lytic ones. These parallels suggest that MTIV4 may adopt a chronic lifestyle or reside in the culture in a population-level carrier state while minimizing activation of host immune responses.

Direct comparison between the bipartite (YN4–YN4HA) and tripartite (MTIV4–YN4–YN4HA) cultures revealed only four marginally upregulated host genes, all close to the significance threshold, and principal component analysis showed tight clustering between these two conditions (see Supplementary Text and Supplementary Fig. 12 for details). Together, these results indicate that the transcriptional state observed in the tripartite culture largely reflects the host response to *Nanobdellati* symbiosis.

### Conclusions

This study describes a novel tripartite system, involving a novel *Nanobdellales* archaeon YN4, associated with the host *Metallosphaera hakonensis* YN4HA and its virus MTIV4. Electron microscopy revealed four distinct types of cell surface structures used by YN4 and its host during their life cycles, including archaella, nanotubes, cones, and spider-web-like (possibly EPS) structures (Fig. 5). Similar cone structures have been reported in other *Nanobdellati*–host systems ^32,60,61^, indicating that this feature is likely conserved among members of *Nanobdellales.* The *Nanobdellales* archaeon YN4 can exist in a free-swimming state and likely uses the observed archaella for swimming motility in liquid culture. In addition, it can make nanotubes with which it can make the first contact to its host and with other YN4 cells (Fig. 5). After this primary attachment, an extensive contact surface is formed by the cones that are present on the cell surface of YN4. Potentially also EPS is formed by either YN4 or its host when they are in co-culture, which could be the first steps towards a mixed biofilm (Fig. 5).

Transcriptomic analysis revealed an extensive host response in the presence of YN4, consistent with earlier RNA-seq studies on *Nanobdellati* systems ^65,66^. In contrast, only a minor response was observed in the presence of the virus MTIV4. Glycan remodeling of the host cell surface was observed in the presence of YN4.

In this system, the presence of the virus enabled the study of the interaction between all three partners. In the presence of YN4, the viral carrying capacity was reduced by ∼68%, indicating a negative impact of the symbiont on viral proliferation. This effect may be due to glycan alterations of the host cell surface, or alternatively due to transcriptional changes in host antiviral defense pathways, such as CRISPR-associated systems, affecting susceptibility to viral infection. Thus, the presence of the *Nanobdellales* archaeon YN4 has a protective effect on the host YN4HA and negatively impacts viral replication.

This tractable tripartite system offers new opportunities to experimentally investigate symbiont–host–virus interactions.

## Methods

### Strains used in this study

To investigate the host specificity of YN4, the following 16 thermoacidophilic strains were cocultivated with YN4*: Acidianus brierleyi* DSM1651^T^, *Metallosphaera cuprina* JCM15769^T^, *M. hakonensis* DSM7519^T^, *M. hakonensis* YN4HA (original host), *M. javensis* AS-7, *M. sedula* DSM5348^T^, *Saccharolobus caldissimus* HS-3^T^, *Scl. shibatae* DSM5389^T^, *Scl. solfataricus* JCM8930^T^, *Sulfodiicoccus acidiphilus* HS-1^T^, *Sulfolobus acidocaldarius* DSM639^T^, *Sulfuracidifex tepidarius* JCM16833^T^, *Saf. metallicus* DSM6482^T^, *Sulfurisphaera javensis* KD-1^T^, *Sfs. ohwakuensis* TA-1^T^, and *Thermoplasma acidophilum* JCM9062^T^. These strains were obtained from the German Collection of Microorganisms and Cell Cultures (DSMZ) or the Japan Collection of Microorganisms (JCM), except for *M. javensis* AS-7, *M. hakonensis* YN4HA, *Scl. caldissimus* HS-3^T^, *Sfd. acidiphilus* HS-1^T^, *Sfs. ohwakuensis* TA-1^T^, and *S. javensis* KD-1^T^, which were originally isolated and stored in our laboratory (Soka University).

### Sampling and enrichment culture

Hot spring water (68.7°C, pH 2.7) was collected at an acidic hot spring, “Yunoike”, in the Kirishima geothermal area, Kagoshima Prefecture, Japan ^71^. The sample was used as an inoculum for enrichment culture using the same procedures as previously described^72^, except that the incubation temperature was set at 65°C. Unless otherwise stated, modified Brock’s basal salt ^73^ supplemented with 1 g/L yeast extract (MBSY medium, pH 3.0) was used for all the cultivation experiments described below. The medium was dispensed into 20 mL screw-capped test tubes (16.5 × 160 mm, ST-16.5L, Nichiden-rika glass). When growth entered the stationary phase, 2 mL of enrichment culture was centrifuged (20,000 × g, 25°C, 30 min). The resulting pellet was collected and used for microbial DNA extraction as described below. The remaining enrichment culture was dispensed into 1 mL aliquots with 10% (v/v) glycerol as a cryoprotectant, and stored at −80°C. The cryostock was later used to establish a pure co-culture.

### Microbial community analysis of enrichment culture

Microbial DNA was extracted from the pellet using the Extrap Soil DNA Plus Kit version 2 (Nippon Steel and Sumikin Eco-Tech) and used for 16S rRNA gene amplicon sequencing, following the procedures described previously ^72^. After removing minor sequences (less than 0.1% of total reads), the microbial community structure was investigated using the QIIME2 pipeline ^74^.

### Establishment of a pure co-culture

To establish a pure co-culture comprising the *Nanobdellota* archaeon YN4 and its host YN4HA, we first performed the purification procedures described previously ^72^. Briefly, three rounds of dilution-to-extinction cultivation were conducted, while monitoring the presence of the *Nanobdellota* archaeon by qPCR using specific primers (Supplementary Table S10). The resulting co-culture was designated as “YN4-YN4HA-MTIV4”. The purity of the strains in YN4-YN4HA-MTIV4 was confirmed by genome sequencing, as described below.

### Exclusion of MTIV4 from the YN4-YN4HA-MTIV4 co-culture

Since the YN4-YN4HA-MTIV4 co-culture contained *Metallosphaera* Turreted Icosahedral Virus (MTIV4), we attempted to exclude this virus through repeated host-switching experiments as follows (Extended Data Fig. 1). (1) The YN4-YN4HA-MTIV4 co-culture was filtered through a sterilized syringe filter (pore size: 0.45 μm; material: polyethersulfone); (2) 1 mL of the filtrate, containing YN4 and MTIV4 but not YN4HA, was inoculated into 5 mL of fresh MBSY medium with 30 μL of *Metallosphaera cuprina* JCM15769^T^ and incubated at 65°C for 1 week, followed by three subcultivations (Extended Data Fig. 1c); (3) steps (1) and (2) were repeated with the host-switched co-culture (Extended Data Fig. 1d,e); (4) the resulting co-culture, composed of YN4 and *M. cuprina*, was designated YN4-Mcup and filtered again through a sterilized 0.45 μm syringe filter; (5) 1 mL of the filtrate, containing only YN4 cells (without MTIV4), was added to 5 mL of fresh MBSY medium with 100 μL of YN4HA and routinely subcultivated (Extended Data Fig. 1f); (6) the resulting co-culture system, consisting of YN4 and YN4HA with successful exclusion of MTIV4, was designated YN4-YN4HA.

### Isolation of original host YN4HA

The original host, *Metallosphaera hakonensis* YN4HA, was isolated by single-colony isolation from the YN4-YN4HA-MTIV4 co-culture on MBSY plate medium solidified with 0.7% (w/v) gellan gum (Wako), 10 mM MgSO_4_ and 2.5 mM CaCl_2_ (Extended Data Fig. 1g). After incubation at 65°C for 2 weeks, an individual colony was picked with a sterilized pipette tip and inoculated into 5 mL of MBSY medium, followed by subcultivation.

### Establishment of MTIV4-YN4HA co-culture

To obtain MTIV4 infecting YN4HA, the following procedures were performed. The YN4-YN4HA-MTIV4 co-culture was filtered through a sterilized syringe filter (pore size: 0.45 μm; material: polyethersulfone), and the filtrate was diluted 10^6^-fold in MBSY medium. 10 μL of the dilution and 120 μL of YN4HA culture were inoculated together into 6 mL of MBSY medium and incubated at 65°C. After 14 days of incubation, the presence of MTIV4 and the absence of YN4 in the system were confirmed by qPCR with specific primers (Supplementary Table 10). The resulting co-culture system, containing MTIV4 and YN4HA, was designated as “MTIV4-YN4HA”.

### Preparation of virus stock

Virus stock solution was prepared from the MTIV4-YN4HA co-culture. A total of 20 mL of MTIV4-YN4HA in the stationary phase was centrifuged (20,000 × g, 20°C, 15 min). The supernatant was filtered through a sterilized syringe filter (pore size: 0.2 μm; material: surfactant-free cellulose acetate). The resulting filtrate was concentrated to approximately 100 μL by ultrafiltration using a Spin-X UF concentrator (molecular weight cut-off: 100,000, Corning). The concentrated filtrate was used as the virus stock for the growth-monitoring test, as described below.

### Growth monitoring tests of each coculture system

The following pure- and co-culture systems, with or without MTIV4, were examined for growth monitoring: (1) pure culture of YN4HA (PURE-YN4HA); (2) MTIV4 infecting YN4HA (MTIV4-YN4HA); (3) co-culture composed of YN4 and YN4HA (YN4-YN4HA); and (4) MTIV4 infecting the YN4-YN4HA co-culture (MTIV4-YN4-YN4HA). Specifically, 50-100 μL of PURE-YN4HA or YN4-YN4HA at the late exponential phase was inoculated into 10 mL of MBSY medium, with or without MTIV4. For the MTIV4-infecting systems (MTIV4-YN4HA or MTIV4-YN4-YN4HA), 13.2 μL of a 10^2 diluted virus stock was added to each system. Incubation was performed at 65°C for 7 days in triplicate. 200-400 μL of the cultures were sampled regularly and stored in a freezer at −25°C until DNA extraction. The growth of each strain was monitored by qPCR as described below.

During the exponential phase, the growth data were fitted to an exponential growth model, N(t) = N_0_ e^rt^, and the specific growth rate (r, day^-1^) was estimated by linear regression of the log-transformed cell densities against time. Similarly, during the late exponential to stationary phase, the data were fitted to a logistic growth model, N(t) = K / [1 + ((K − N_0_)/N_0_) e^−r·t^], to obtain the carrying capacity (K, cells or particles ml^-1^). Curve fitting was performed independently for each biological replicate, and the resulting parameters were averaged to yield mean ± SD values.

### Determination of the growth temperature and pH ranges for YN4

The YN4-YN4HA co-culture was cultivated at temperatures of 45-80°C (pH 3.0) and pH values of 1.0-5.5 (65°C), in duplicate for up to 28 days. 200-400 μL of the cultures were sampled regularly and stored at −25°C until DNA extraction. The growth of each strain was monitored by qPCR, as described below.

### Quantitative PCR

The growth of YN4, YN4HA, and MTIV4 in each co-culture system was monitored by qPCR using specific primers for each strain (Supplementary Table 10). The same qPCR conditions described previously were used ^72^.

Total DNA was extracted from the samples using the Maxwell RSC Instrument and the Maxwell RSC Cultured Cells DNA Kit (Promega). Briefly, 100 μL of raw culture (containing both microbial cells and virus particles) or virus stock (prepared as described above) was mixed with 300 μL of 10 mM Tris-HCl (pH 8.0), and the mixture was subjected to DNA extraction using the Maxwell RSC Cultured Cells DNA Kit (Promega) according to the manufacturer’s instructions. A 5 μL aliquot of the extracted DNA was subjected to qPCR with specific primers designed using NCBI/Primer-BLAST ^75^, targeting a single-copy gene for each strain (Supplementary Table 10). The standard DNA for qPCR was prepared from the PCR product of each gene, which was purified using the FastGene Gel/PCR Extraction Kit (Nippon Genetics). Concentrations of the standard DNA were determined with the Qubit dsDNA HS Assay Kit (Invitrogen), and copy numbers were calculated using the online NEBio calculator (https://nebiocalculator.neb.com/#!/dsdnaamt)

### Host specificity test of *Nanobdellota* archaeon YN4

The 16 strains shown above were cultivated in MBSY medium (pH 3.0), except for *Saf. tepidarius* JCM16833^T^ and *Saf. metallicus* DSM6482^T^, which were cultivated in MBSY medium (pH 3.0) with 1 g/L elemental sulphur. Pure YN4 cells were collected from the YN4-YN4HA co-culture by filtration through a sterilized syringe filter (pore size: 0.45 μm; material: polyethersulfone). 1 mL of the filtrate (YN4 cells) and 30 μL of each pure strain were inoculated into 5 mL of MBSY medium. As a negative control, 1 mL of the filtrate, without inoculation of any of the strains, was inoculated into 5 mL of MBSY medium in triplicate. Incubation was carried out under aerobic conditions at 65°C except for *T. acidophilum* JCM9062^T^, which was cultivated at 55°C. After 7 days of cultivation, 30 μL of the cultures were subcultivated into 6 mL of MBSY medium and incubated under the same conditions for a further 7 days. The growth of YN4 in these cultures was assessed on day 7 of each cultivation by qPCR with specific primers (Supplementary table 10).

### Light and electron microscopy

General cell morphology was assessed by phase-contrast microscopy (Optiphot-2; Nikon) and differential interference contrast microscopy (BX51; Olympus) at room temperature. Cells stained with SYBR Green I using the same protocol described previously ^76^ were observed by fluorescence microscopy (BX51; Olympus). Scanning electron microscopy (JSM-7500F; JEOL) and negative-staining transmission electron microscopy were performed using the same method as described previously ^76^.

### Fluorescence in situ hybridization

Fluorescence in situ hybridization (FISH) was performed using the method previously described ^31^, with some modifications. The FISH probes used for YN4 (Nano-R4_TEX: GTATTCCCGTGGCGACTGC) and YN4HA (SFB-R_FAM: CGGTTACTAGGGATTCCTCG), both targeting the 16S rRNA gene, were designed using NCBI/Primer-BLAST ^75^. The probes were labelled with either Texas-red or 6-carboxyfluorescein at their 5’ end. Cells were fixed with 2% paraformaldehyde overnight, washed in phosphate-buffered saline (PBS), and resuspended in PBS before spotting 15 µL onto a silane-coated glass slide (S8111; MATSUNAMI). The spotted sample was dried, incubated in 100 µL of 0.25 N HCl solution for 30 minutes at room temperature, rinsed in distilled water, and dehydrated by serial treatment with 50%, 80%, 90%, and 100% ethanol solutions (3 minutes each, 2 times). Hybridization was carried out with probes (final concentration: 1 pmol/µL each) in buffer (0.9 M NaCl, 20 mM Tris-HCl, 0.1% SDS, pH 7.5) at 48°C overnight. After hybridization, the slide was washed in buffer (0.9 M NaCl, 20 mM Tris-HCl, 0.1% SDS, 5 mM EDTA, pH 7.5) at 48°C for 20 min, lightly rinsed in distilled water, and mounted with mounting medium (SlowFade Gold Antifade reagent; Life Technologies). The fluorescence signals were detected with an epifluorescence microscope (BX63; Olympus) fitted with filter sets specific for Texas-red and fluorescein.

### Genome sequencing and assembly

Genomic DNA from the YN4-YN4HA-MTIV4 co-culture was extracted as described previously ^72^. Nanopore long-read sequencing was performed on a MinION sequencer with a R10.4.1 flow cell (Oxford Nanopore Technologies: ONT), using the Native Barcoding kit 24 V14 (SQK-NBD114.24; ONT) according to the manufacturer’s protocol. Illumina short-read sequencing was conducted on the NextSeq platform (2 x 300 bp). The DNA library for the NextSeq sequencing was prepared with the Illumina DNA Prep kit (Illumina). The resulting short and long reads were quality-filtered using the procedures described previously ^72^. Given the excess data, 300 Mbp of the quality-filtered long reads were subsampled using SeqKit v.2.7.0 ^77^ and genome assembly was performed with Flye v.2.9.2 ^78^. The resulting assemblies were then polished three times with Medaka v.2.0.1 (https://github.com/nanoporetech/medaka), followed by three rounds of polishing with NextPolish v.1.4.1 ^79^ using 1 Gbp of the quality-filtered short reads. To confirm the purity of the co-culture, the quality-filtered short and long reads were mapped to the assemblies using CoverM v.0.7.0 (https://github.com/wwood/CoverM). The ratio of mapped reads and mean sequencing coverage were then calculated.

### Genome analyses

Genome annotation of the *Nanobdellota* archaeon YN4 and its host YN4HA was performed as previously described ^72^. Subcellular localization of each protein encoded in the genome sequences was predicted using PSORTb 3.0.314 ^80^. Genes related to antiviral defence systems and CRISPR arrays were identified using PADLOC-DB v2.0.0. ^81^. The 16S rRNA gene sequence similarity between YN4 and its related species was calculated by BLASTN against the NCBI-nt database, and average amino acid identity (AAI) values were calculated using EzAAI v1.2.4 ^82^.

### Phylogenomic analysis of *Nanobdellota* archaeon YN4

Amino acid sequences of 53 archaeal phylogenetically informative marker genes, as defined by the GTDB database (release 220), were collected and aligned using GTDB-Tk v.2.4.0 ^83^. The aligned sequences were trimmed using TrimAL ver.1.5.0. ^84^ with the automated1 option. A maximum-likelihood phylogenetic tree was reconstructed using IQ-TREE2 ver. 2.3.6 ^85^ with the LG+F+I+G4 model (automatically selected as the best model).

Additionally, maximum-likelihood trees were reconstructed from the 16S rRNA gene and universal single-copy marker genes. For the 16S rRNA gene phylogenetic tree, taxonomically related sequences to strain YN4 were retrieved from the NCBI database and analysed with IQ-TREE2 v2.3.6 ^85^ using the GTR+F+R3 model. For the phylogenetic tree based on universal single-copy marker genes, 17 genomes from the order *Nanobdellales* were retrieved from the NCBI database and used for downstream analyses with the YN4 genome. The universal single-copy marker genes were retrieved using fetchMGs 2.0 ^86,87^. A total of nine genes (COG0048, COG0086, COG0093, COG0094, COG0098, COG0124, COG0185, COG0186, and COG0522), present in all genomes, were selected for phylogenetic reconstruction. Sequences were aligned with MAFFT v7.525 with auto mode ^88^, and spurious sequences or poorly aligned regions were removed using TrimAL v1.5.0 ^84^. A phylogenetic tree was then reconstructed using IQ-TREE2 v2.2.7 ^85^ with the LG+F+R3 model. In all phylogenetic analyses, bootstrap support values were calculated using 1,000 replicates. The genomes from the order *Nanobdellales* were also analyzed using BlastKOALA (https://www.kegg.jp/blastkoala/) and visualized using KEGG-Decoder ^89,90^.

### Genome analysis of MTIV4

Although the initial polished genome assembly contained a contig with a near-complete version of the MTIV4 genome, its terminal inverted repeats were truncated. To obtain an optimal assembly of the MTIV4 genome, we performed a targeted reassembly of the viral reads from the nanopore data with Flye v.2.9.2 ^78^, followed by manual curation and extension of the terminal inverted repeats based on reads mapping to the edges (Minimap2 v2.28 ^91^, IGV v2.12.3 ^92^).

Viral genes were called and annotated with pharokka v1.7.2 ^93^, PHROG ^94^, and phold v0.1.4 [https://github.com/gbouras13/phold]. Initial comparison to known viruses was carried out via the NCBI web BLAST+ services ^95^. Detailed protein-level whole-genome comparisons with the three previously described MTIV viruses ^96,97^, as well as clustering of all viral proteins to determine single-copy core proteins (present once in each genome), were performed with MMseqs2 ^98^. Multiple sequence alignments of the core proteins were computed with MAFFT v7.490 ^88^ and the phylogenetic tree was reconstructed with FastTree v2.1.11 ^99^ from the concatenated core protein alignment. The MTIV core-protein phylogeny and whole-genome comparisons were visualized with ggtree v3.10.1 ^100^ and gggenomes v1.0.1 ^101^.

### RNA Sequencing

At the late exponential growth phase (day 4), microbial cells were harvested by centrifugation (20,000 × g, 4°C, 15 min) from approximately 2 mL of pure or co-culture samples, namely: PURE-YN4HA, MTIV4-YN4HA, YN4-YN4HA, and MTIV4-YN4-YN4HA. Each condition was prepared in biological triplicates. Total RNA was extracted from cell pellets using the Monarch Total RNA Miniprep Kit (New England BioLabs) according to the manufacturer’s protocol. The extracted RNAs were submitted to Novogene Co., Ltd. for RNA sequencing on an Illumina NovaSeq X Plus platform with 150 bp paired-end reads, generating ∼3 Gb (∼20 million reads) of sequencing data per sample. The resulting reads were used for downstream analyses.

### Transcriptome Analysis

Raw reads were quality-filtered with fastp ^102^ v.0.24.0 to remove adapter sequences and low-quality bases. Filtered reads were aligned to rRNA and tRNA genes from strains YN4HA and YN4 using Bowtie2 v.2.5.4 ^103^, and only unmapped reads were retained for further analysis. The retained reads were then mapped to the coding sequences (CDSs) of strains YN4HA, YN4, and MTIV4 using Bowtie2 v.2.5.4 with default parameters.

Read counts for each CDS were quantified with featureCounts v.2.1.1 ^104^. Transcript abundance was normalized to transcripts per million (TPM), and differential gene expression analysis was performed using DESeq2 v.1.48.1 ^105^; both analyses were based on biological triplicates. Significantly differentially expressed genes were identified using adjusted p-values (padj < 0.01) and log2 fold-change thresholds (>1). Volcano plots were generated to visualize expression differences across conditions.

### Sequence- and structure-based analysis of YNH_2540

The amino acid sequence of YNH_2540—a hypothetical protein from *Metallosphaera hakonensis* YN4HA that was the most abundantly expressed transcript in YN4HA during co-culture with YN4—was analyzed using TMHMM v2.0 ^106^, Phobius ^107^, and LipoP v1.0 ^108^ to predict transmembrane regions and signal peptides. Coiled-coil motifs were evaluated with DeepCoil2 ^109^. Structural modelling was performed using AlphaFold2 via the AlphaFold Server (https://alphafoldserver.com). Structural similarity was assessed using Foldseek (https://search.foldseek.com/) and the DALI server (http://ekhidna2.biocenter.helsinki.fi/dali/).

### Glycoproteome analysis

Peptide samples were prepared and analyzed using an Orbitrap Fusion^TM^ Tribrid^TM^ Mass Spectrometer (Thermo Fisher Scientific), as described previously ^110,64^. The LC-MS/MS data were then analyzed using Byonic software v3.11.3 (Protein Metrics Inc., Cupertino, CA) ^111^, following previously described methods ^110,64^.

### Live-cell imaging with precise temperature control

Co-cultures were imaged using a temperature-control unit (VAHEAT, Interference GmbH, Germany) with a Smart Substrate (SmS-R-4) chamber heated to 65 °C. Time-lapse images were captured in phase-contrast mode with a 100× Plan-Apochromat oil objective (NA = 1.46) at a rate of 178 frames per 300 seconds. Imaging was conducted on a Zeiss Axio Observer 7 inverted fluorescence microscope (Carl Zeiss Microscopy GmbH, Jena, Germany) equipped with a Colibri 7 LED illumination system and a Prime BSI Express sCMOS camera (Teledyne Photometrics, Tucson, AZ, USA).

### Probe design, synthesis, and chemical labelling

A set of 35 polynucleotide probes targeting the MTIV4 genome was designed using genePROBER ^112^. The 300-bp sequences are listed in Supplemental Table 11. Probes were synthesised as gBlocks (500 ng each; IDT, San Jose, CA, USA) and resuspended in 5 mM Tris, 1 mM EDTA (pH 8.0). A total of 2 µg of MTIV4-specific polynucleotides was labelled in a single reaction with the ULYSIS Alexa Fluor 594 Kit (Thermo Fisher Scientific) and purified using Micro Bio-Spin Columns (Bio-Rad). Probe specificity was confirmed by testing against uninfected cells to ensure no nonspecific binding.

### Virus targeting direct-geneFISH

Fluorescence in situ hybridisation (FISH) was performed following the direct-geneFISH protocol by Barrero-Canosa and Moraru (2021) ^113^. *M. hakonensis* cells infected with MTIV4 were fixed with 20% paraformaldehyde (Thermo Fisher GmbH) according to the core protocol. Fixed cells (10 µL) were placed on Superfrost™ Plus slides within Silicone Isolators™ (Grace Bio-labs), air-dried, and sequentially dehydrated in 50%, 70%, and 100% ethanol for 10 minutes each. Cells were permeabilised by briefly dipping slides in 0.1 M HCl at room temperature, followed by a quick rinse in ultrapure water and subsequent air-drying at room temperature. Samples were then covered with 80 µl of hybridisation buffer (35% formamide, 5× SSC, 20% dextran sulfate, 20 mM EDTA, 0.25 mg/mL salmon sperm DNA, 0.25 mg/mL yeast RNA, 1× blocking reagent, 0.1% SDS, and nuclease-free water) containing probes at 30 pg/µL. Probes and templates were denatured at 92 °C for 30 min, then hybridized at 46 °C for 2 h in a humidity chamber. Samples were washed sequentially in buffer I (2× SSC, 0.1% SDS, 5 min at 48 °C), buffer II (0.1× SSC, 0.1% SDS, 30 min at 48 °C), PBS for 25 minutes, a 1-minute wash in ultrapure water, and finally air-dried. Cells were counterstained for 10 minutes in the dark with 10 µL of SlowFade™ Diamond with DAPI (Thermo Fisher Scientific) and mounted with a #1.5 high-precision coverslip (Paul Marienfeld GmbH). Imaging was performed with a 100× Plan-Apochromat oil objective (NA 1.46) on a Zeiss Axio Observer 7 microscope equipped with a Colibri 7 LED system and a Prime BSI Express sCMOS camera (Teledyne Photometrics). Phase-contrast images were acquired at an exposure time of 9.8 ms. DAPI was excited at 385 nm and detected with a BP 460/50 filter (9.8 ms). Alexa Fluor 594 was excited at 555 nm and detected with a BP 630/75 filter (9.8 ms). Image acquisition and processing were performed using Zeiss ZEN Blue (v3.5).

### Sample vitrification and cryo-electron tomography data acquisition

*Nanobdellales*-host samples were concentrated to an optical density at 600 nm (OD600) of 8-10. They were then mixed in a 4:1 ratio with 10 nm colloidal gold beads precoated with 1% bovine serum albumin (BSA; Sigma-Aldrich, Australia). This mixture was applied to glow-discharged copper R2/2 Quantifoil holey carbon grids (Quantifoil Micro Tools GmbH, Jena, Germany). The grids were blotted for 4-5 seconds under 100% humidity and subsequently plunged into liquid ethane using a Vitrobot Mark IV (Thermo Fisher Scientific).

Initial screening of the grids was performed using a Thermo Scientific™ Talos™ Arctica transmission electron microscope. High-resolution tilt-series of the co-culture systems were acquired using an FEI Titan Krios G4 transmission electron microscope operating at 300 keV (Thermo Fisher Scientific), equipped with a BioQuantum K3 Imaging Filter (slit width 20 eV) and a K3 direct electron detector (Gatan). Tilt series were collected automatically using FEI Tomography 5.10, ranging from −51° to +51° at 3° intervals, with a defocus range of −2 to −5 μm, a total electron fluence of ∼120 e⁻/Å², and a pixel size of 1.603 Å.

### Tomogram reconstruction

Tilt-series movies were imported into Scipion v3.8.3 ^114^. for processing. Motion correction was performed using MotionCor3 v1.0.1 ^115^, followed by manual tilt-series alignment in IMOD v4.11.24 ^116^. The aligned tilt series were binned eightfold, and tomograms were reconstructed using Tomo3D v2.2 ^117^ with the simultaneous iterative reconstruction technique (SIRT) for 30 iterations, resulting in a final pixel size of 12.824 Å and a reconstruction thickness of 400 pixels. Selected tomograms were further denoised using CryoLithe ^118^ to enhance contrast for visualization and segmentation.

### Tomogram segmentation and visualization/ 3D-segmentation

Tomogram visualization and segmentation were performed using Dragonfly software v2025.1 ^119^. For automated segmentation, a neural network-based 5-class U-Net was employed, using 2.5D input consisting of five consecutive slices. Segmentation results were further refined manually to ensure accuracy.

## Acknowledgement

This work was supported by an NHMRC grant (APP1196924 to DG), an HFSP grant (RGEC33/2023 to DG, HDS and TQ), JSPS KAKENHI grants (21K15153 and 24K18201 to HDS; 23K13204 and 23KJ2174 to YT: JP20H03322 and JP23K20307 to SN), the Joint Research Program of the National Institutes of Natural Sciences (NINS), and the Exploratory Research Center on Life and Living Systems (ExCELLS program nos. 22EXC601 and 25EXC601-2 to HDS and SN). HDS was supported by the RIKEN Special Postdoctoral Researcher Program. Cryo-EM data were collected at the Ian Holmes Imaging Centre, Bio21 Institute, University of Melbourne. We thank NITTETSU MINING CO., LTD. for their support with field sampling.

## Data availability

Raw read data of genome/RNA sequencing have been deposited in DDBJ/ENA/GenBank under the accession numbers DRR641936-DRR641941 and DRR802084-DRR802087, respectively. Genomic sequences of YN4, Metallosphaera hakonensis YN4HA, and archaeal virus MTIV4 were deposited under the accession numbers AP046538, AP046537, and AP046536, respectively. Mass spectrometry proteomics data have been deposited to the ProteomeXchange Consortium via the jPOST repository ^120^ under accession codes PXD076122–PXD076126. The datasets are accessible via the following URLs and access keys: PXD076122, access key 9077: https://repository.jpostdb.org/preview/45248044869c390bf9eea4; PXD076123, access key 3303: https://repository.jpostdb.org/preview/96547101169c39105d5043; PXD076124, access key 5132: https://repository.jpostdb.org/preview/28987701969c3913cb5ff4; PXD076125, access key 1184: https://repository.jpostdb.org/preview/97806615269c39185d1986.

---

**Extended Data Fig. 1.**
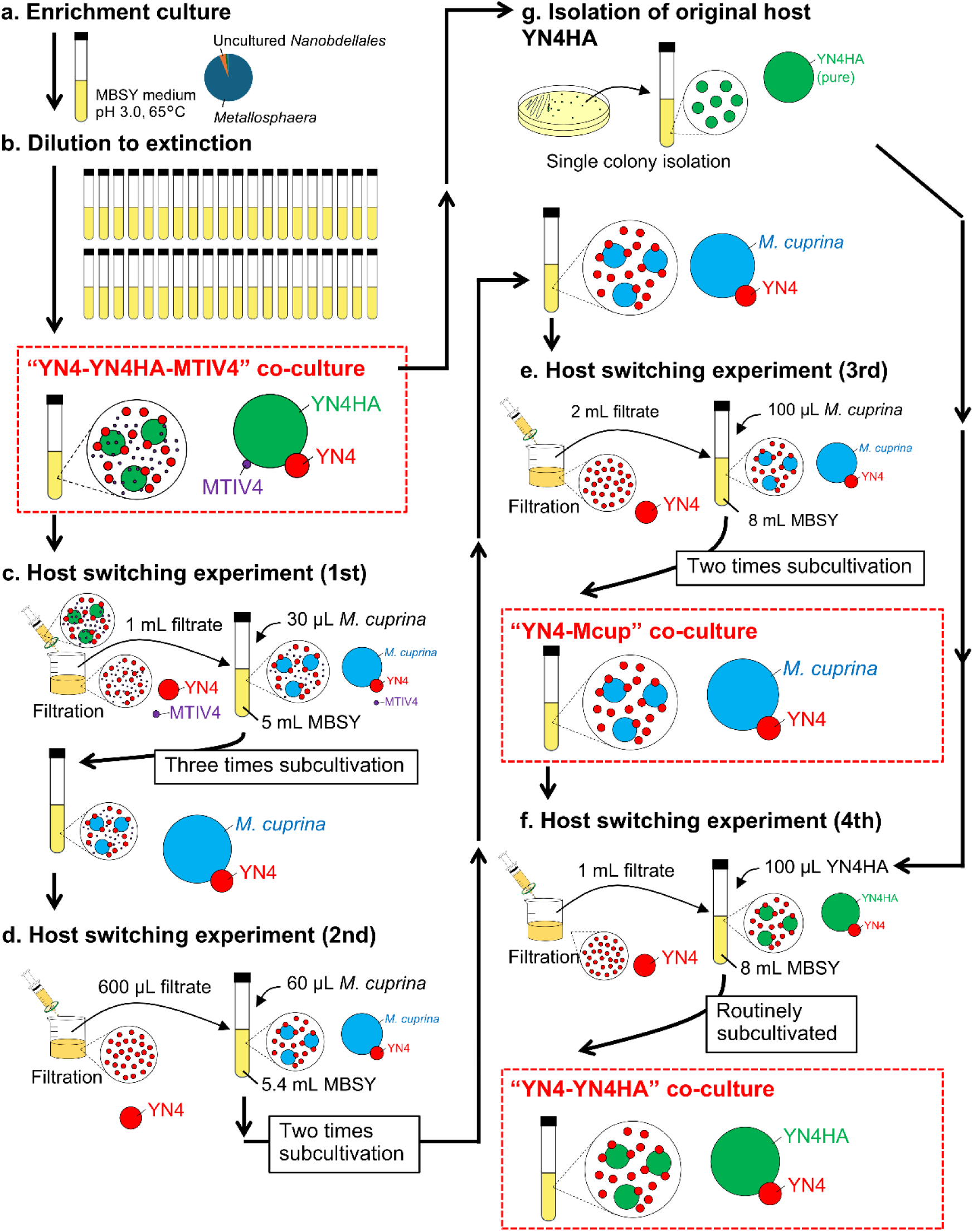
Scheme of a process for the establishment of each coculture system.

**Extended Data Fig. 2.**
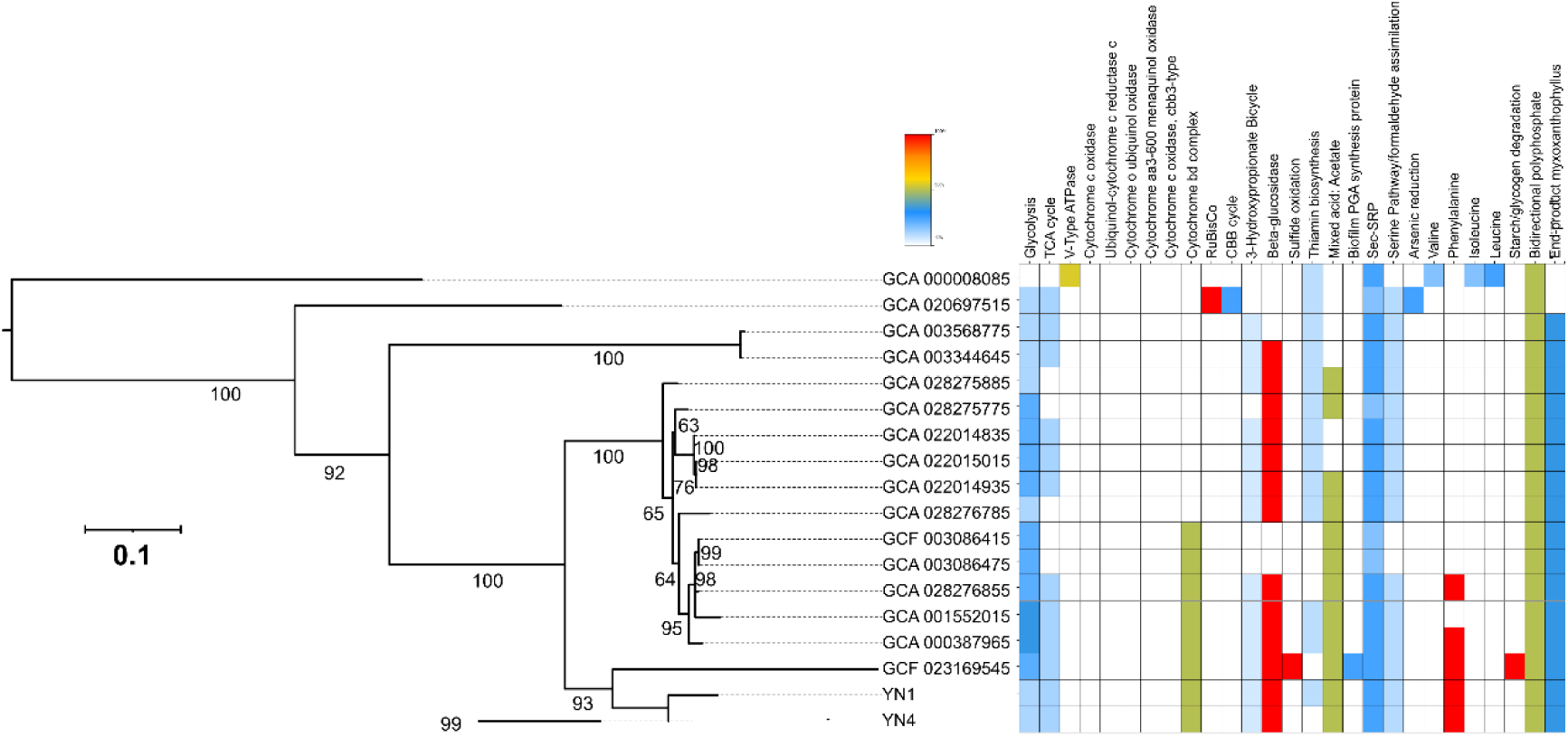
KEGG Decoder heatmap of YN4 with related *Nanobdellati* species based on KEGG annotation. The heatmap was constructed based on the completeness of metabolic pathway inferred from the KEGG Decoder (i.e., the presence or absence of genes). The colour scale is shown in the upper left, where red represents a complete or highly complete pathway, followed by yellow and blue, while white indicates the absence of the pathway.

**Extended Data Fig. 3.**
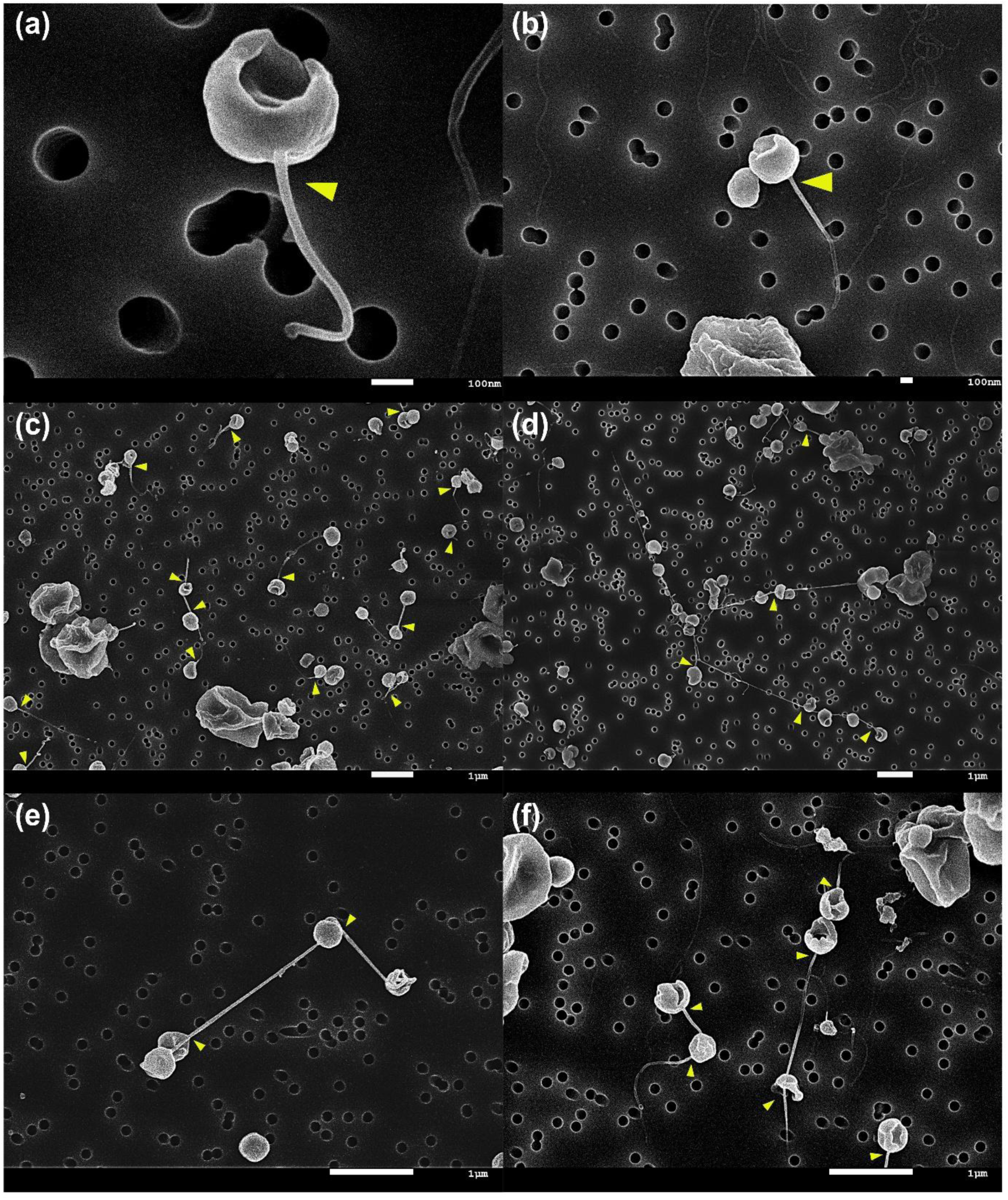
SEM images showing nanotubes emerged from YN4 cells (yellow arrows).

**Extended Data Fig. 4.**
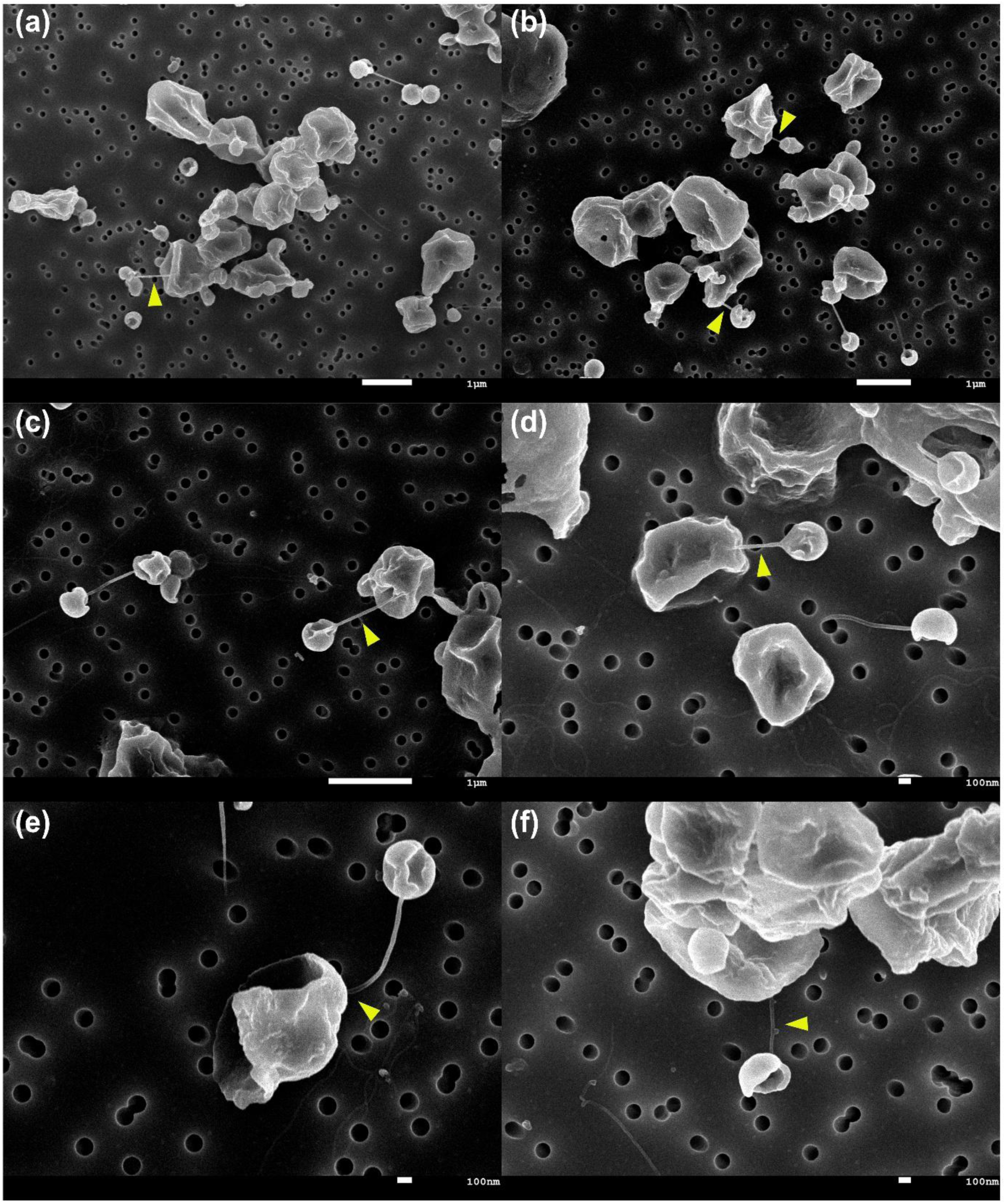
SEM images showing YN4 nanotubes attaching to a YN4HA cell (yellow arrows).

**Extended Data Fig. 5.**
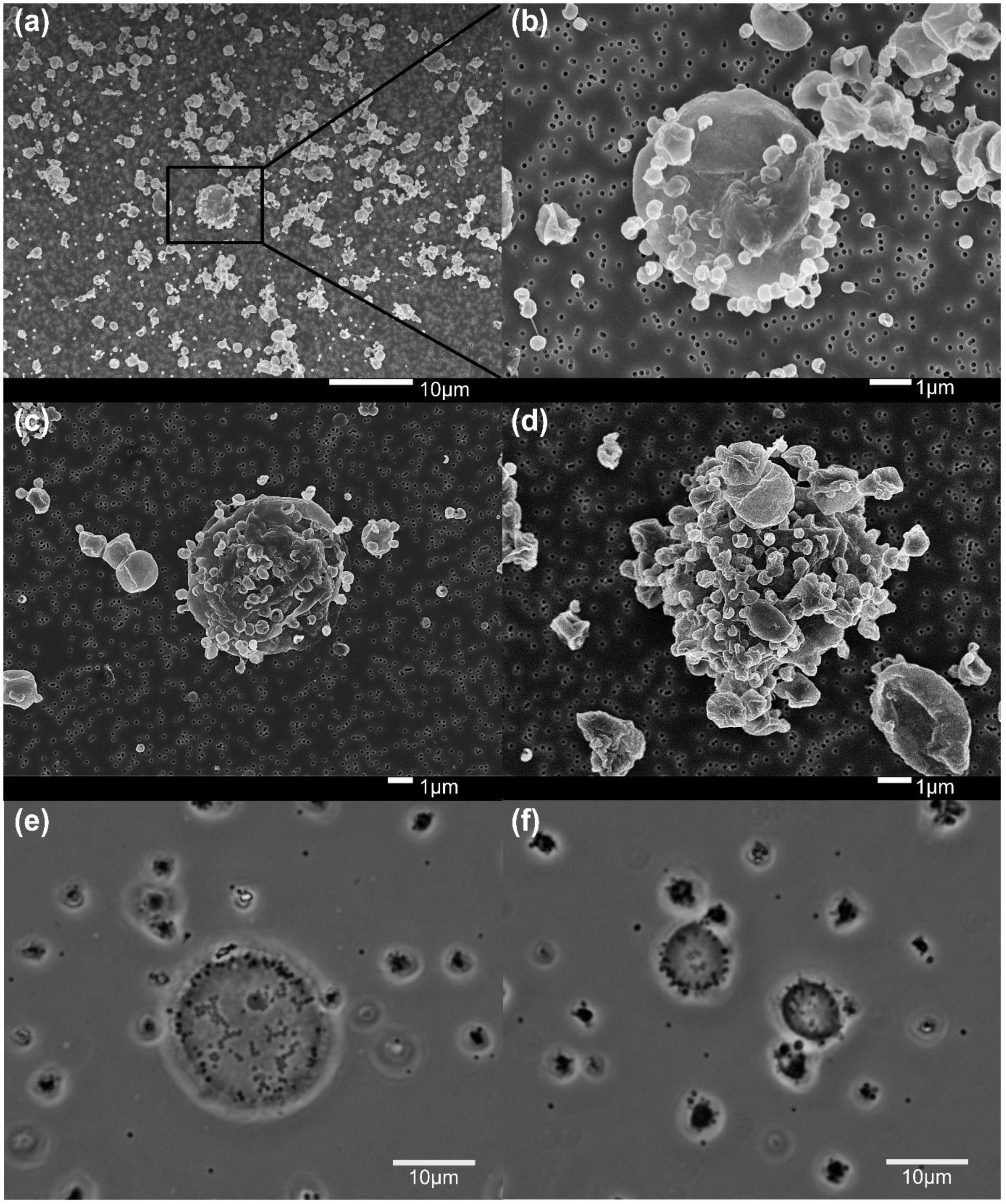
SEM (A-D) and phase-contrast microscopy (E-F) showing enlarged YN4HA cells.

**Extended Data Fig. 6.**
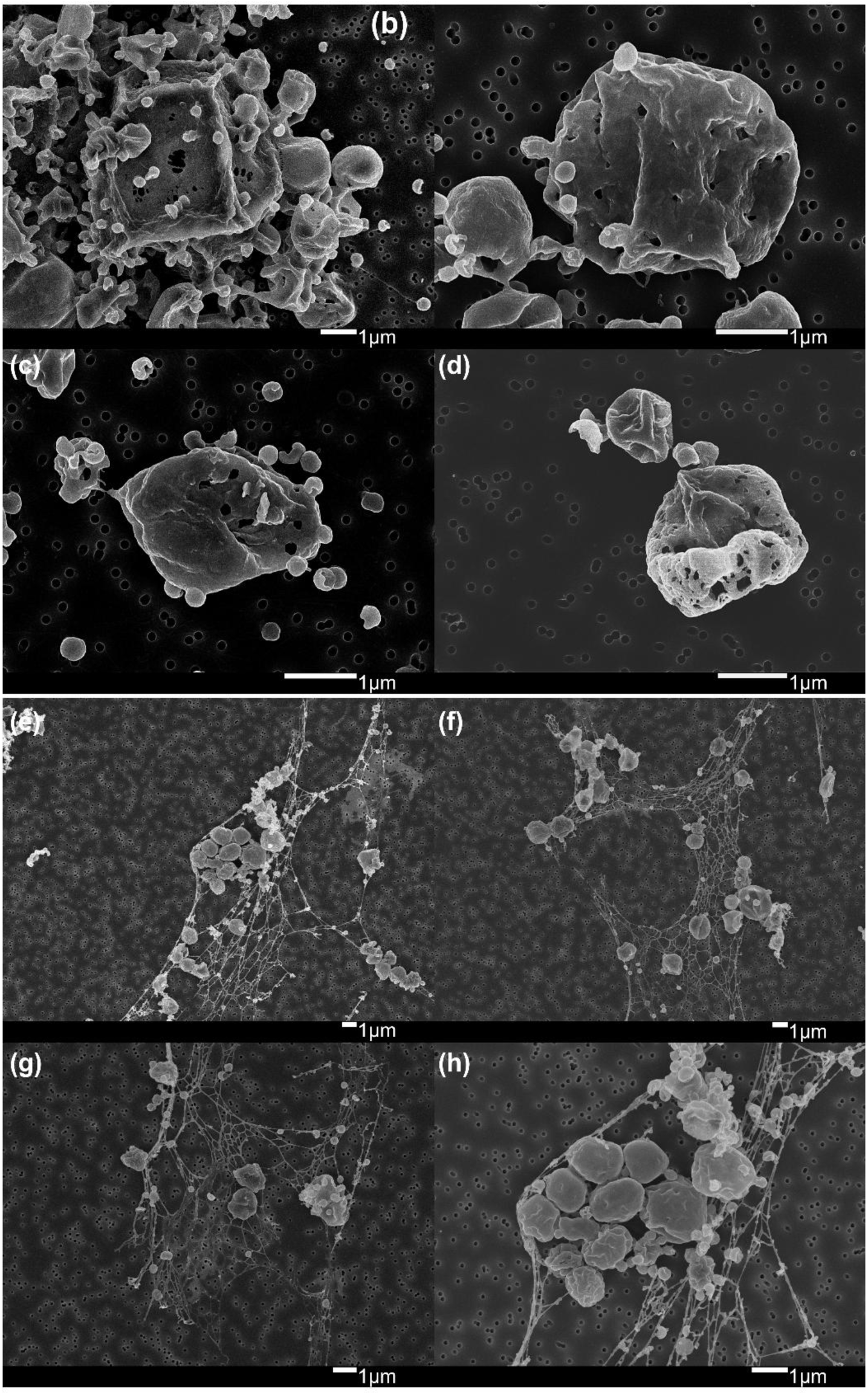
SEM images showing holes on the cell surface of YN4HA (a-d) and spider web-like structures (e-h).

**Extended Data Fig. 7.**
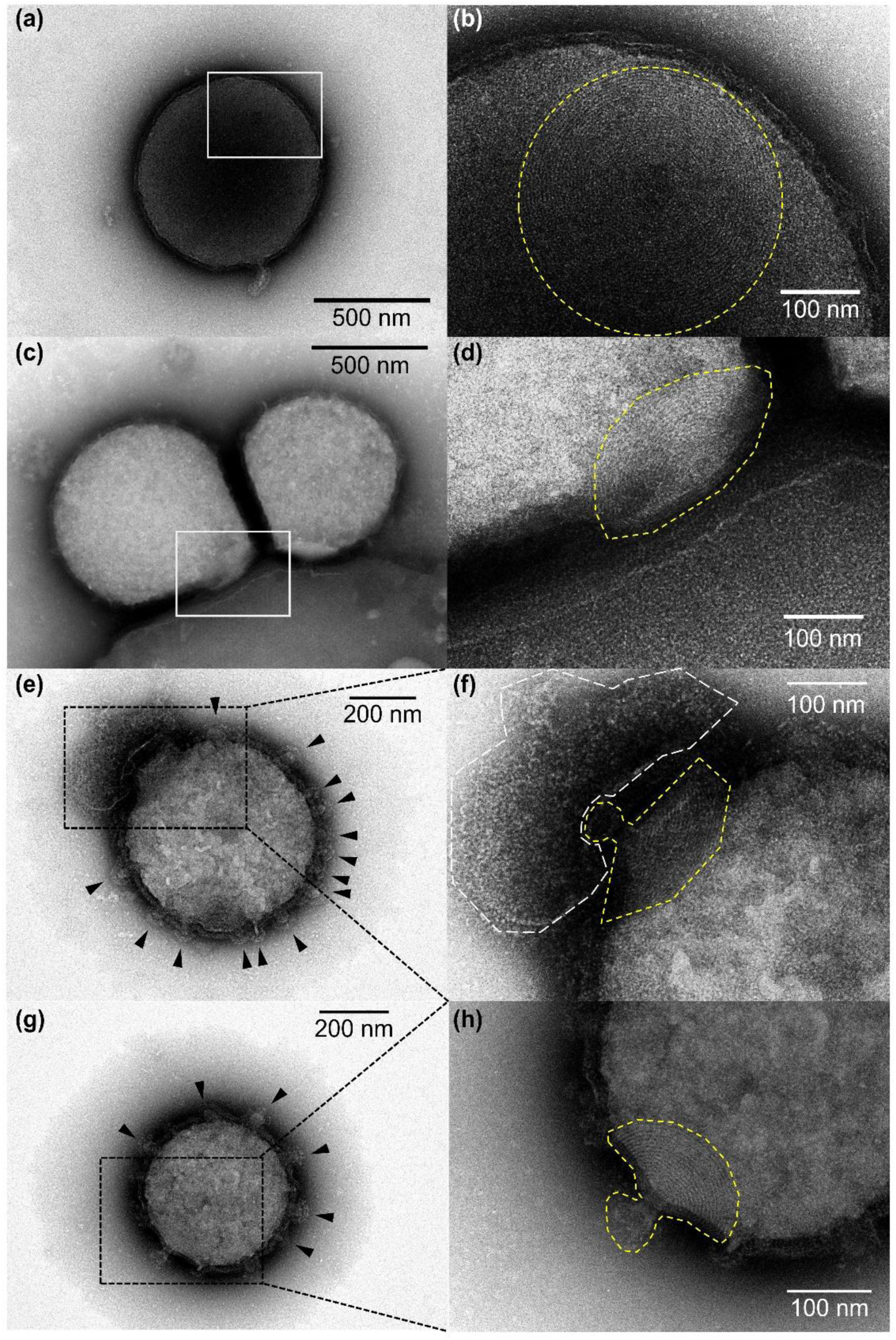
Negative staining TEM showing a cone structure on the surface of YN4 cells. Black arrows indicate membrane vesicle-like structures. White line indicates S-layer-like structure of YN4HA. Yellow lines indicate a cone structure with a short tube-like structure at its vertex opening. White line indicates S-layer like structure of YN4HA.

**Extended Data Fig. 8.**
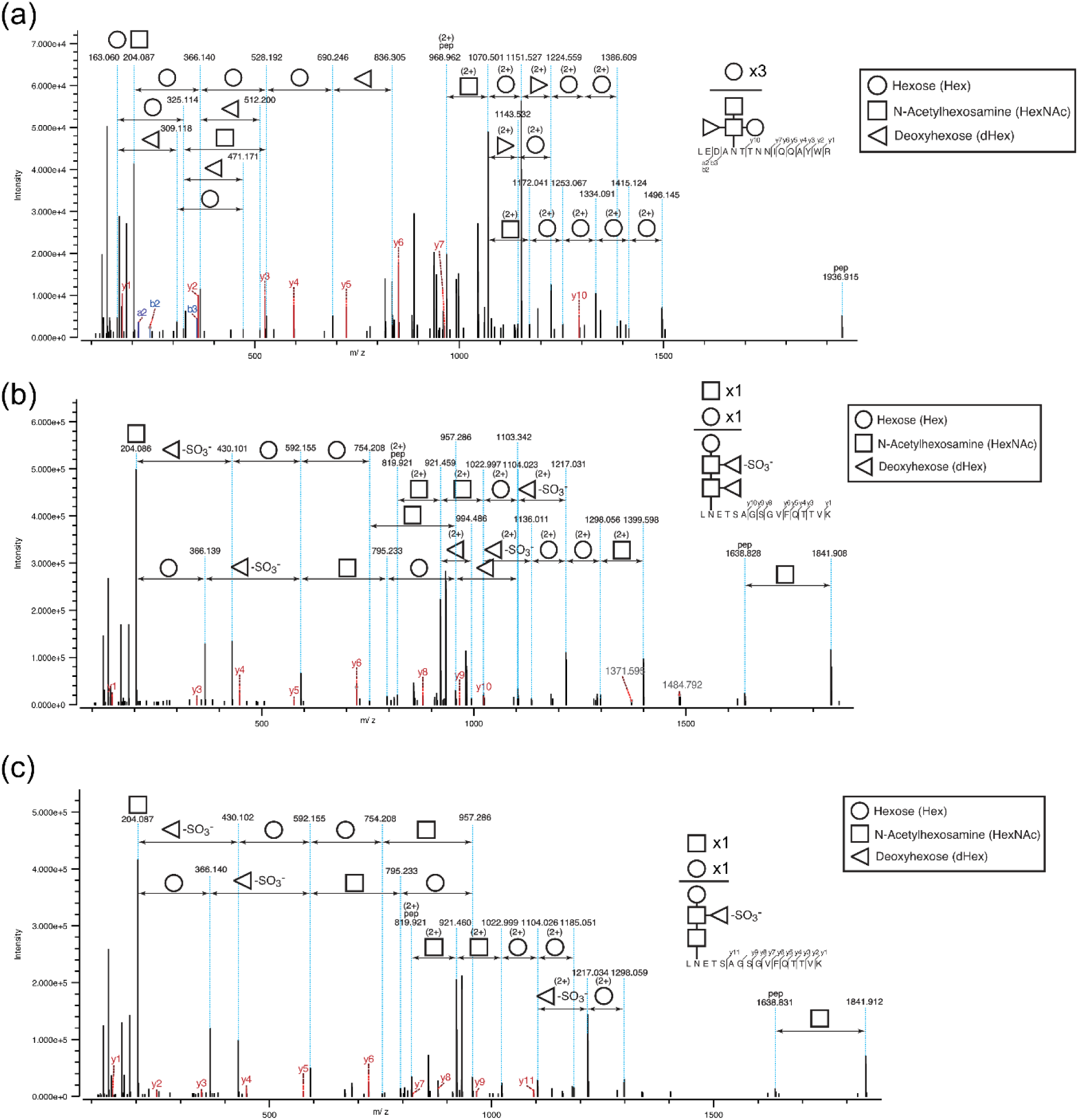
MS/MS spectra showing glycan assignments on proteins. (a) Higher-energy collision dissociation (HCD) spectrum showing the N-glycan structure of 1200.43 Da on a hypothetical protein (YN4_730) of strain YN4. (b) Higher-energy collision dissociation (HCD) spectrum showing the N-glycan on S-layer protein A of strain YN4HA. (c) Higher-energy collision dissociation (HCD) spectrum showing the N-glycan on S-layer protein A of strain YN4HA co-cultured with strain YN4.

**Extended Data Fig. 9.**
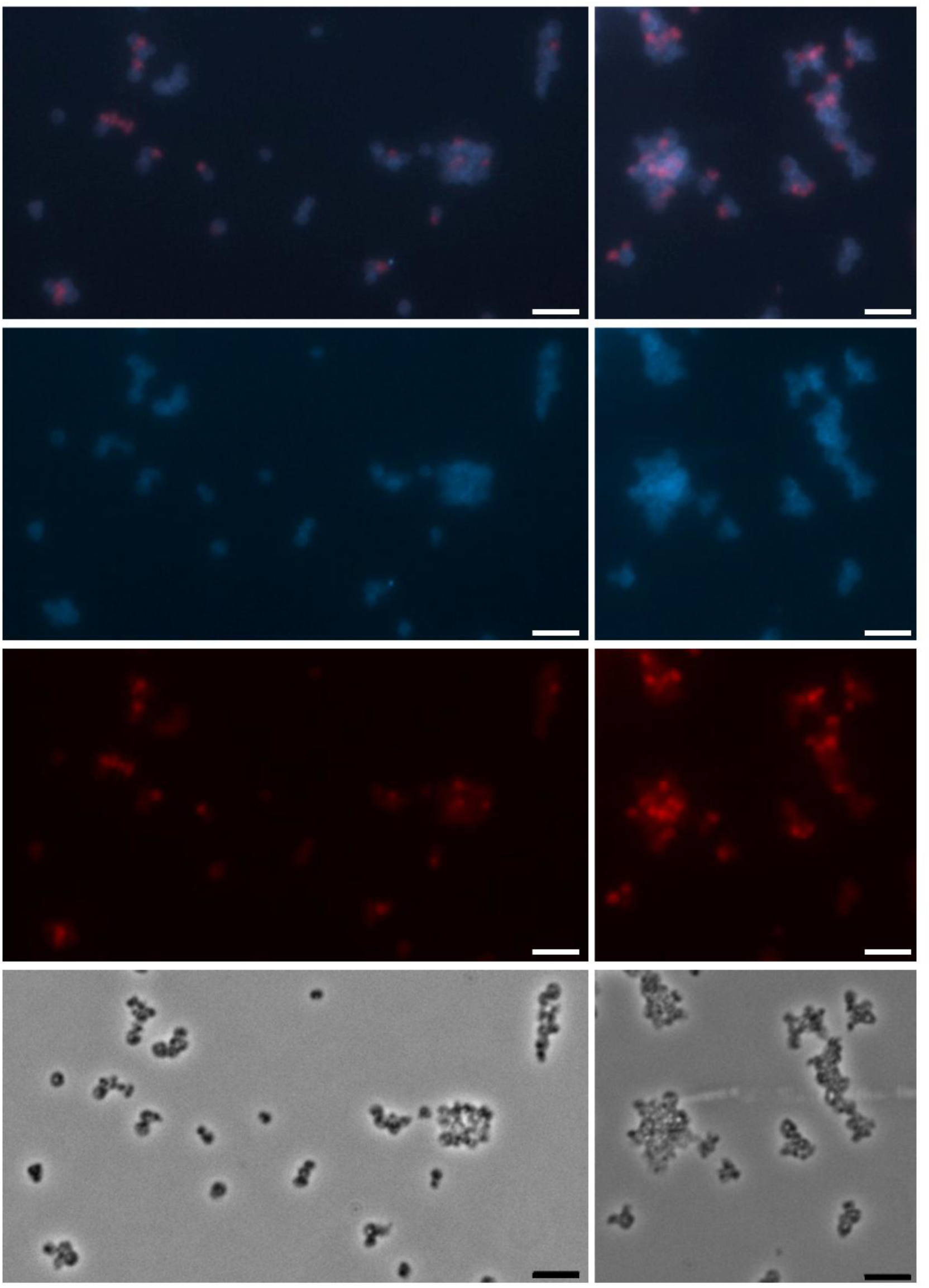
Detection of MTIV4 in *M. hakonensis* using virus-targeted direct-gene FISH. Intracellular and attached MTIV4 (red) were visualised using Alexa Fluor 594–labelled polynucleotide probes. Panels show (A) merged DAPI and MTIV4 signal, (B) DAPI-stained total DNA, (C) MTIV4 probe signal, and (D) phase contrast. Scale bars: 5 µm.

