## Supplemental Data 1 for "A Tripartite Co-culture System Reveals Defensive Mutualism Between a *Nanobdellati* Symbiont and Its Host"

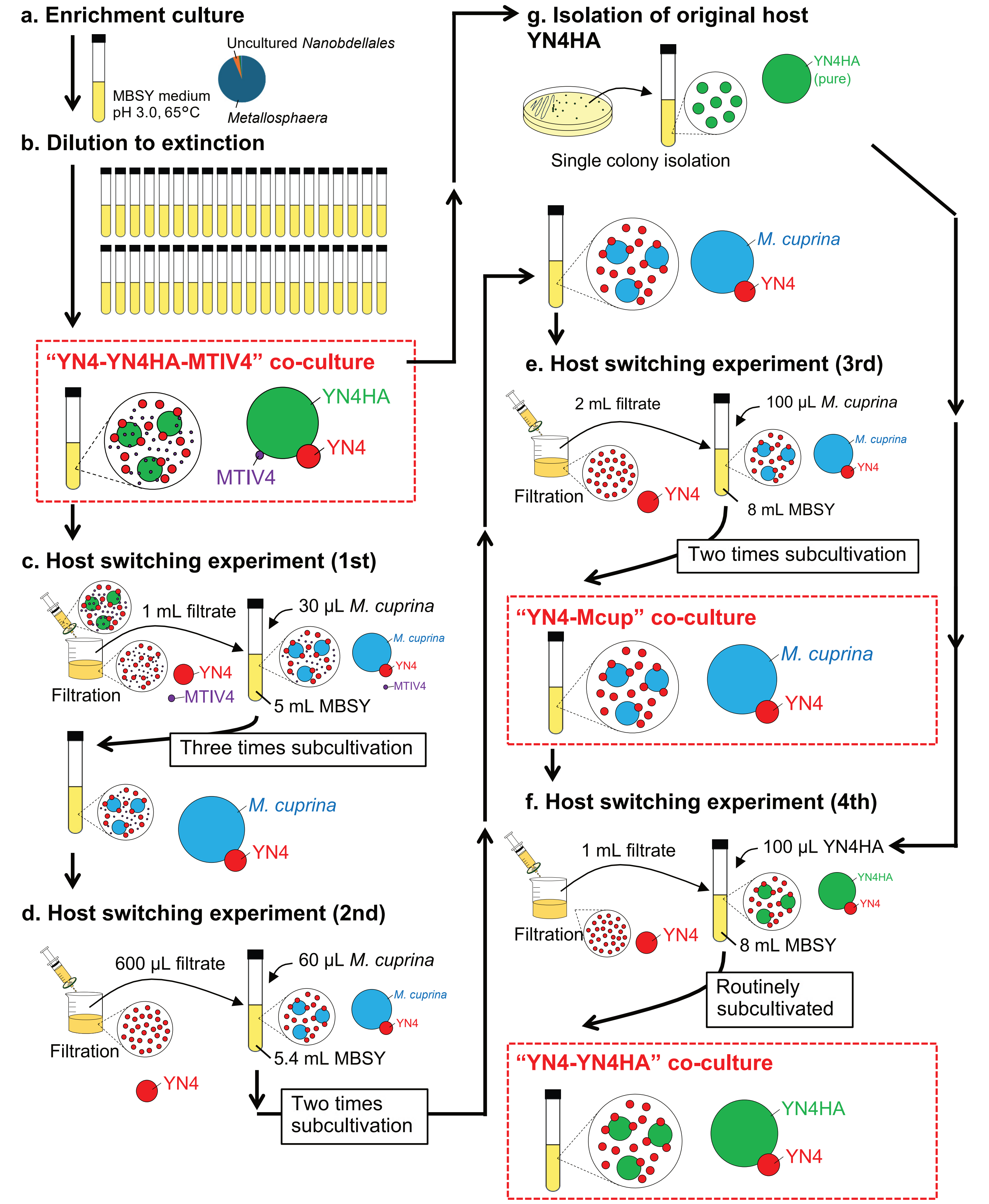


**Extended Data Fig. 1 |** Scheme of a process for the establishment of each coculture system.

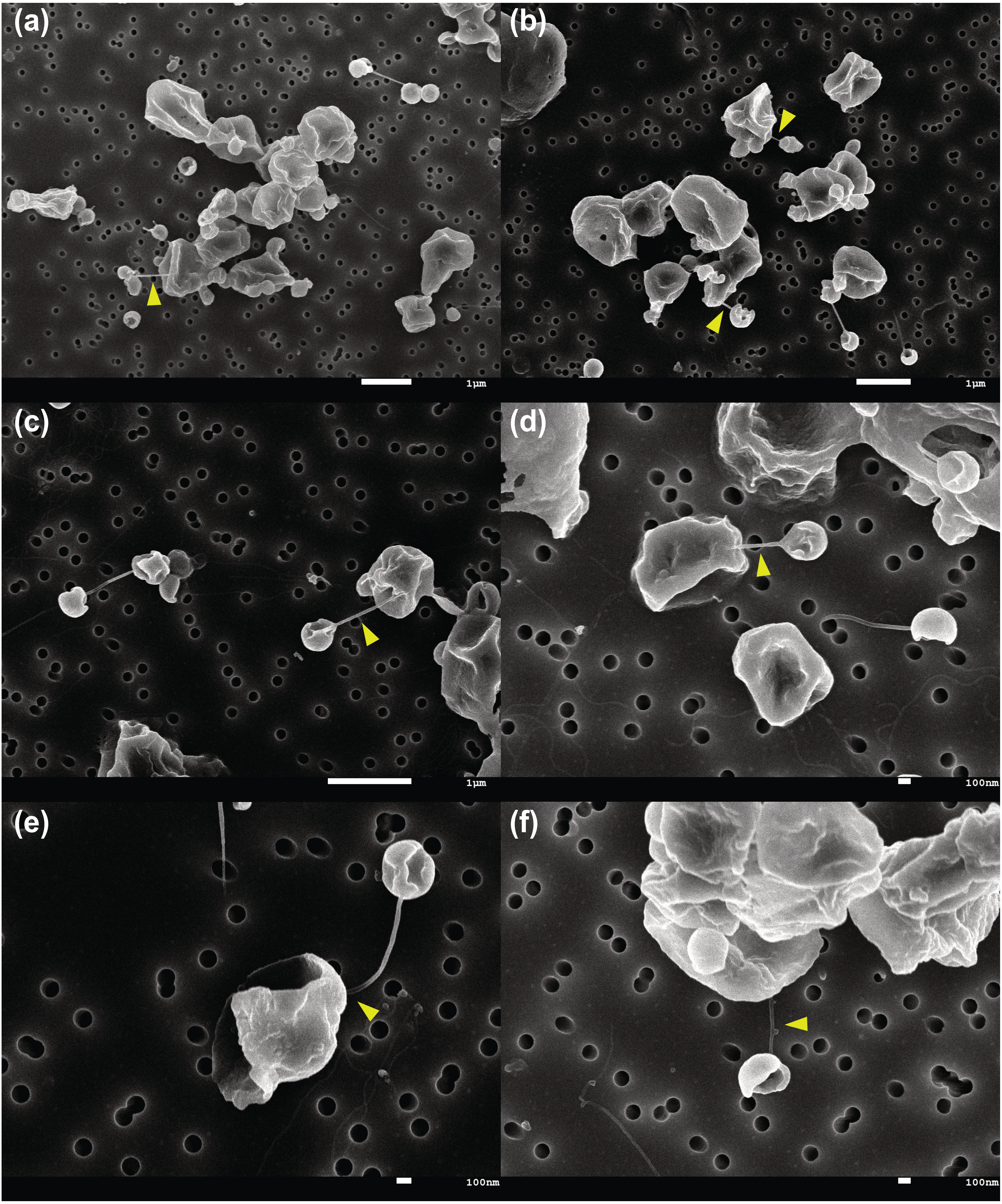


**Extended Data Fig. 4 |** SEM images showing YN4 nanotubes attaching to a YN4HA cell (yellow arrows).

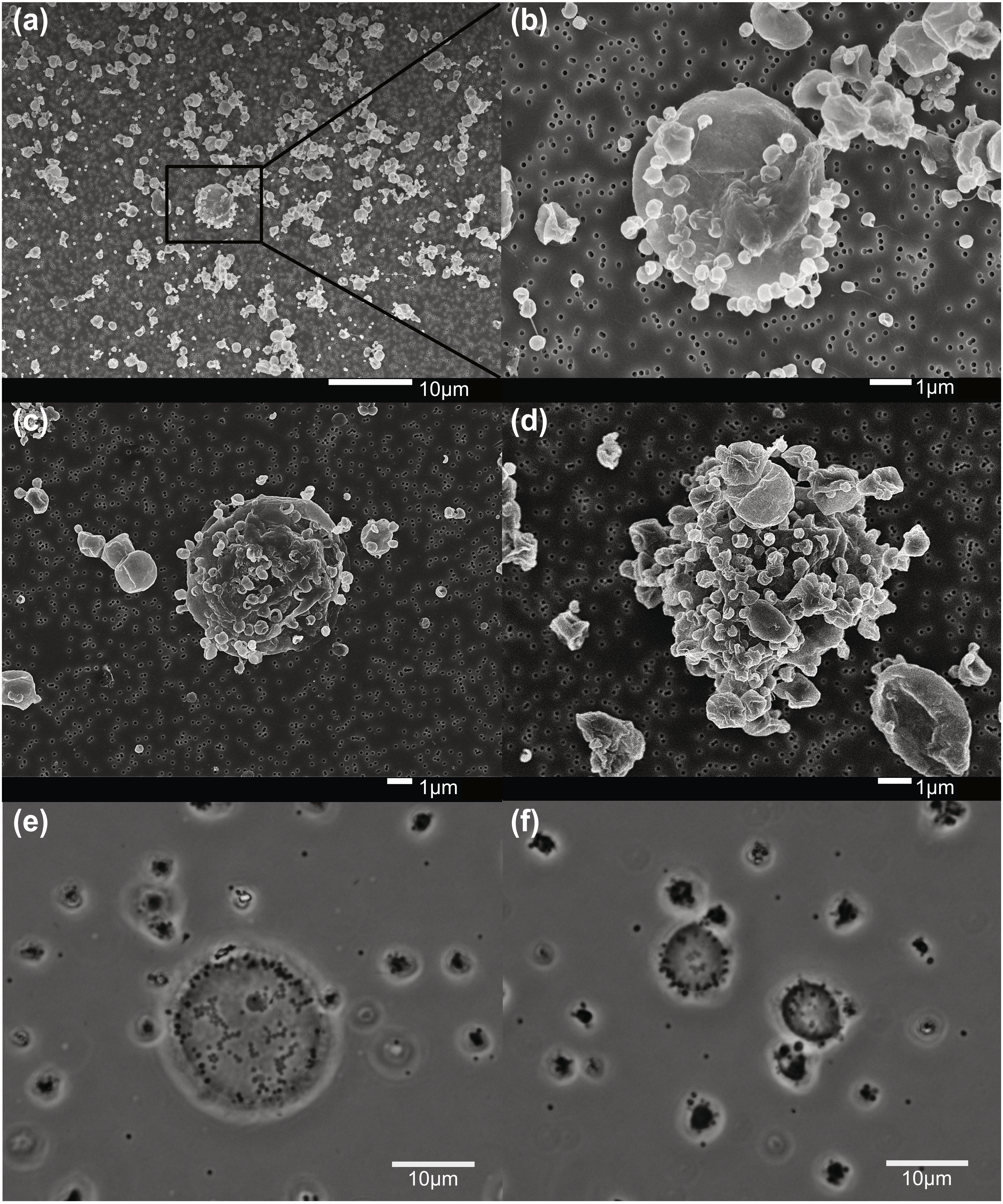


**Extended Data Fig. 5 |** SEM (A-D) and phase-contrast microscopy (E-F) showing enlarged YN4HA cells.

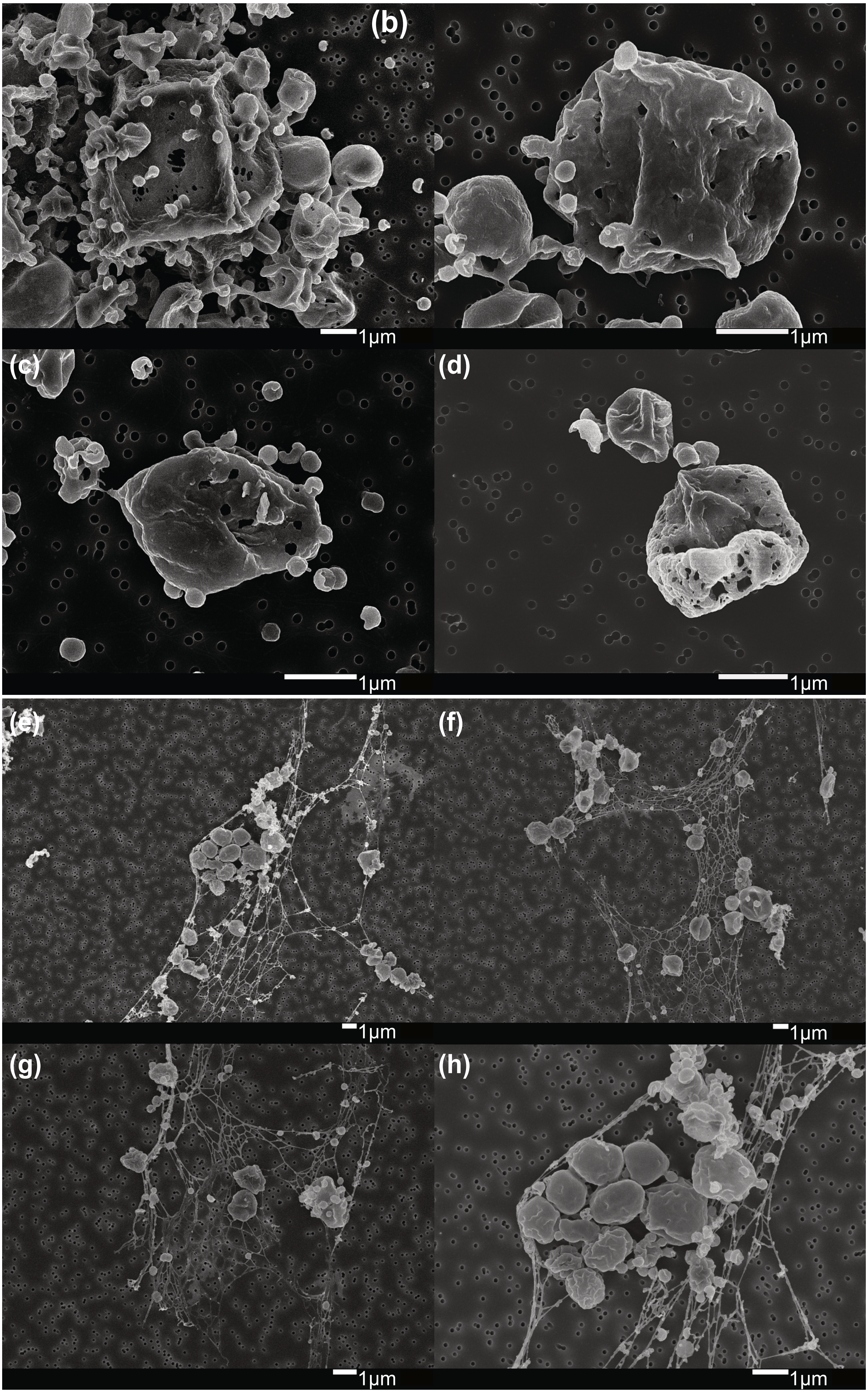


**Extended Data Fig. 6 |** SEM images showing holes on the cell surface of YN4HA (a-d) and spider web-like structures (e-h).

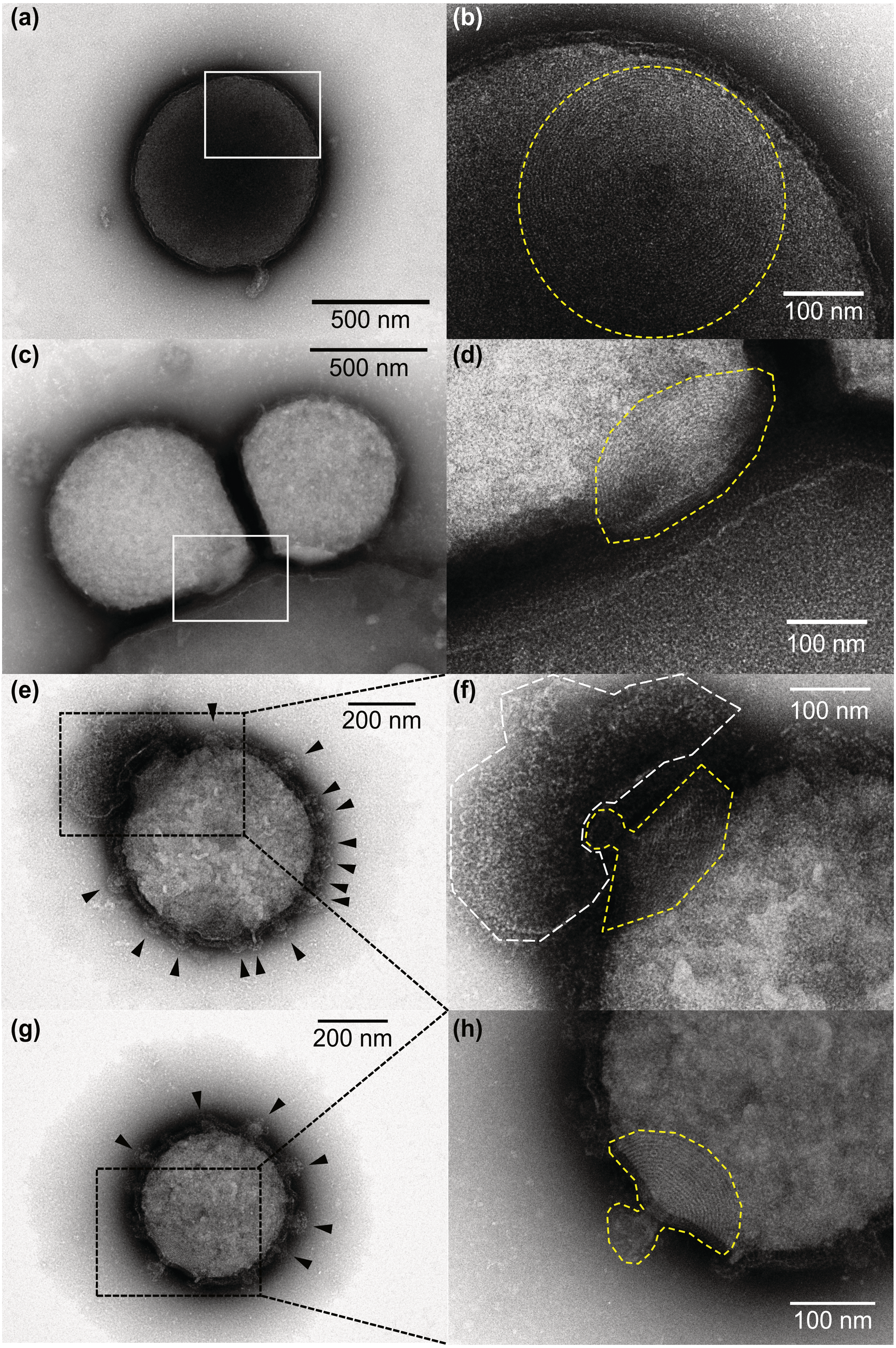


**Extended Data Fig. 7 |** Negative staining TEM showing a cone structure on the surface of YN4 cells. Black arrows indicate membrane vesicle-like structures. White line indicates S-layer-like structure of YN4HA. Yellow lines indicate a cone structure with a short tube-like structure at its vertex opening. White line indicates S-layer like structure of YN4HA.

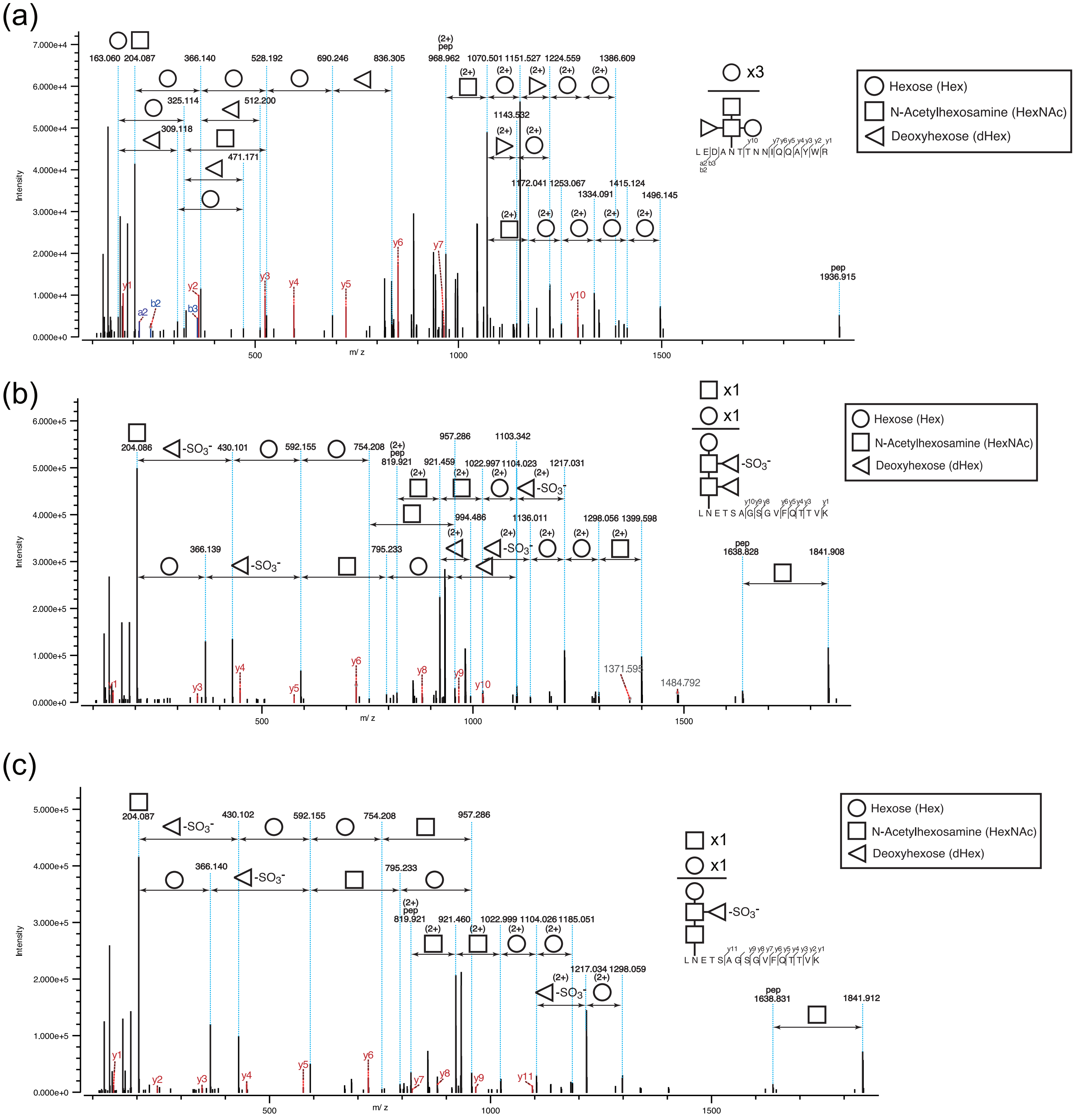


**Extended Data Fig. 8 | MS/MS spectra showing glycan assignments on proteins.** (a) Higher-energy collision dissociation (HCD) spectrum showing the N-glycan structure of 1200.43 Da on a hypothetical protein (YN4_730) of strain YN4. (b) Higher-energy collision dissociation (HCD) spectrum showing the N-glycan on S-layer protein A of strain YN4HA. (c) Higher-energy collision dissociation (HCD) spectrum showing the N-glycan on S-layer protein A of strain YN4HA co-cultured with strain YN4.

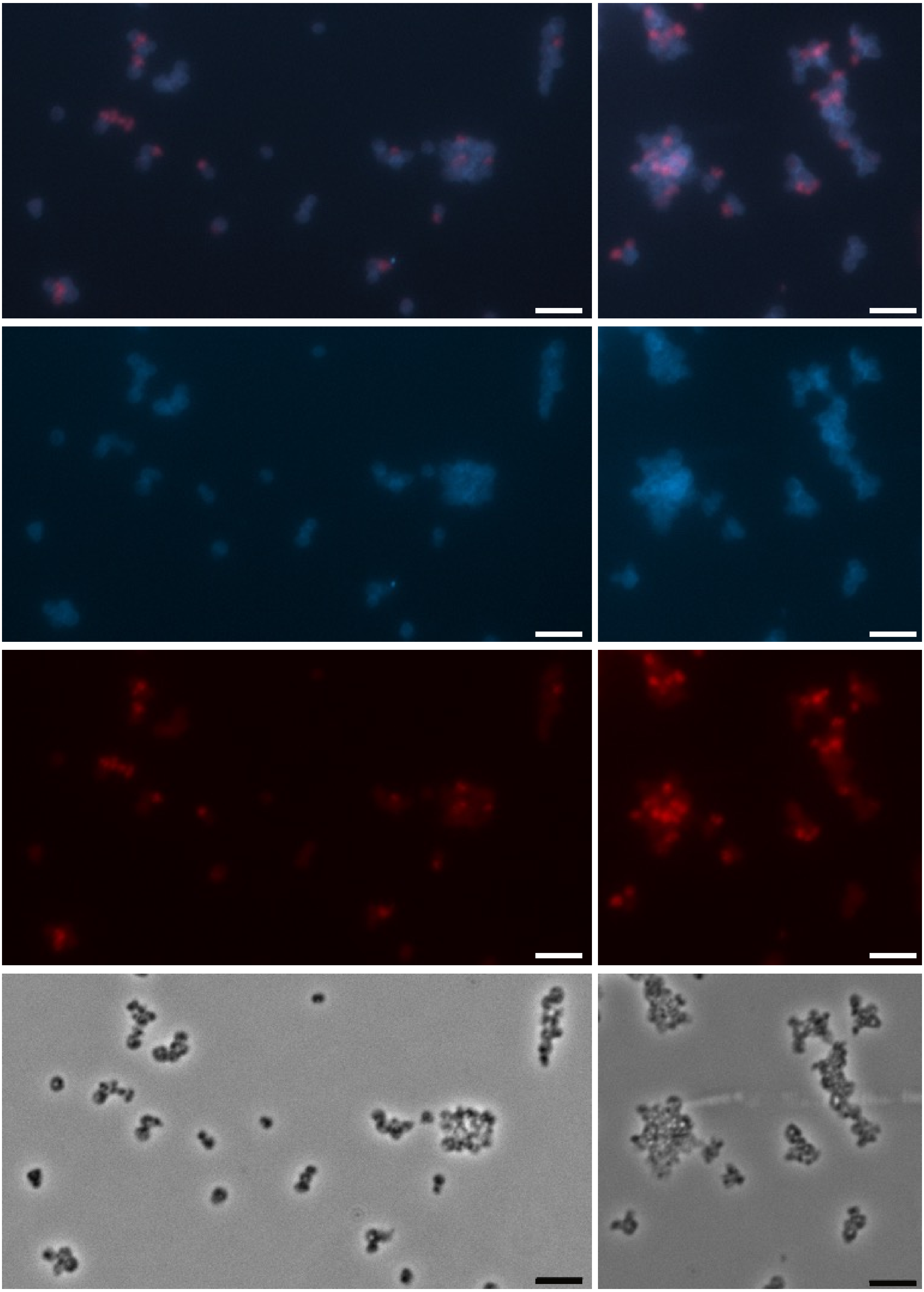

**Extended Data Fig. 9 | Detection of MTIV4 in *M. hakonensis* using virus-targeted direct-gene FISH.** Intracellular and attached MTIV4 (red) were visualised using Alexa Fluor 594–labelled polynucleotide probes. Panels show (A) merged DAPI and MTIV4 signal, (B) DAPI-stained total DNA, (C) MTIV4 probe signal, and (D) phase contrast. Scale bars: 5 μm.

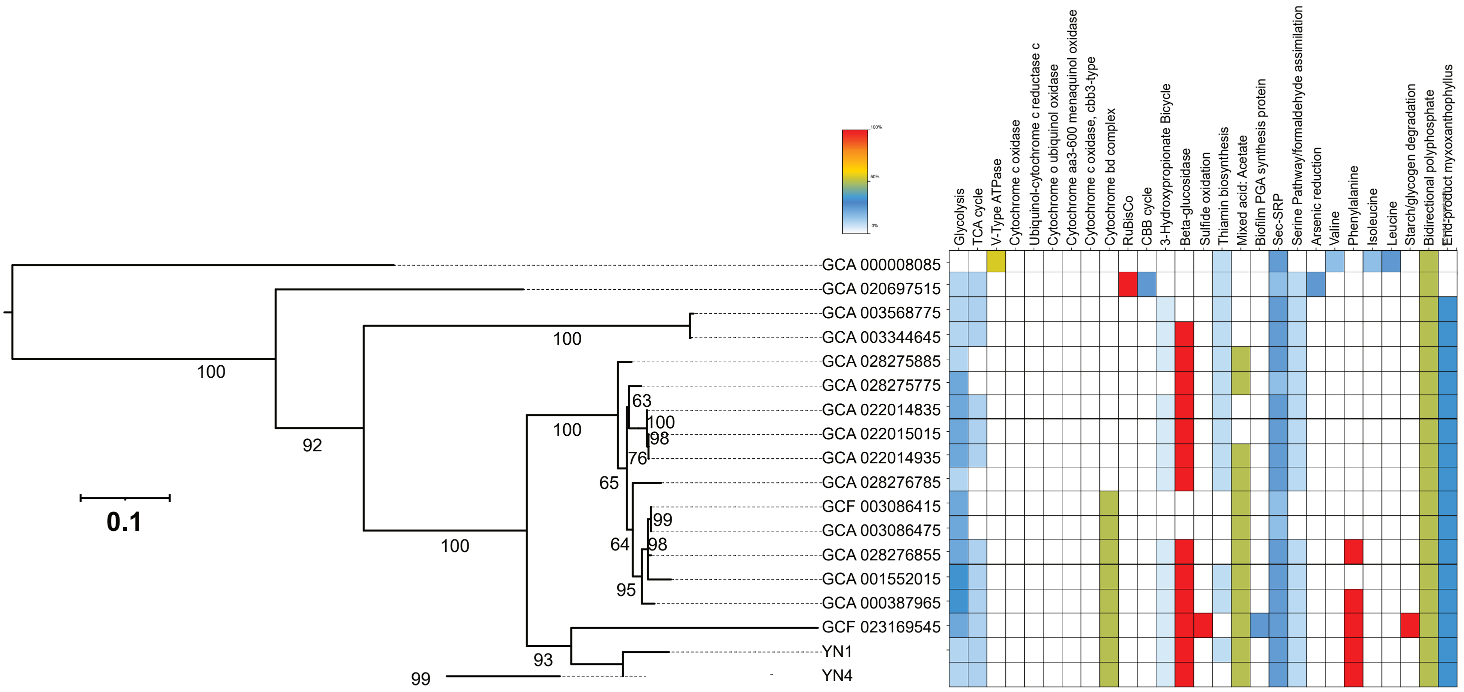


**Extended Data Fig. 2 |** KEGG Decoder heatmap of YN4 with related *Nanobdellati* species based on KEGG annotation. The heatmap was constructed based on the completeness of metabolic pathway inferred from the KEGG Decoder (i.e., the presence or absence of genes). The colour scale is shown in the upper left, where red represents a complete or highly complete pathway, followed by yellow and blue, while white indicates the absence of the pathway.

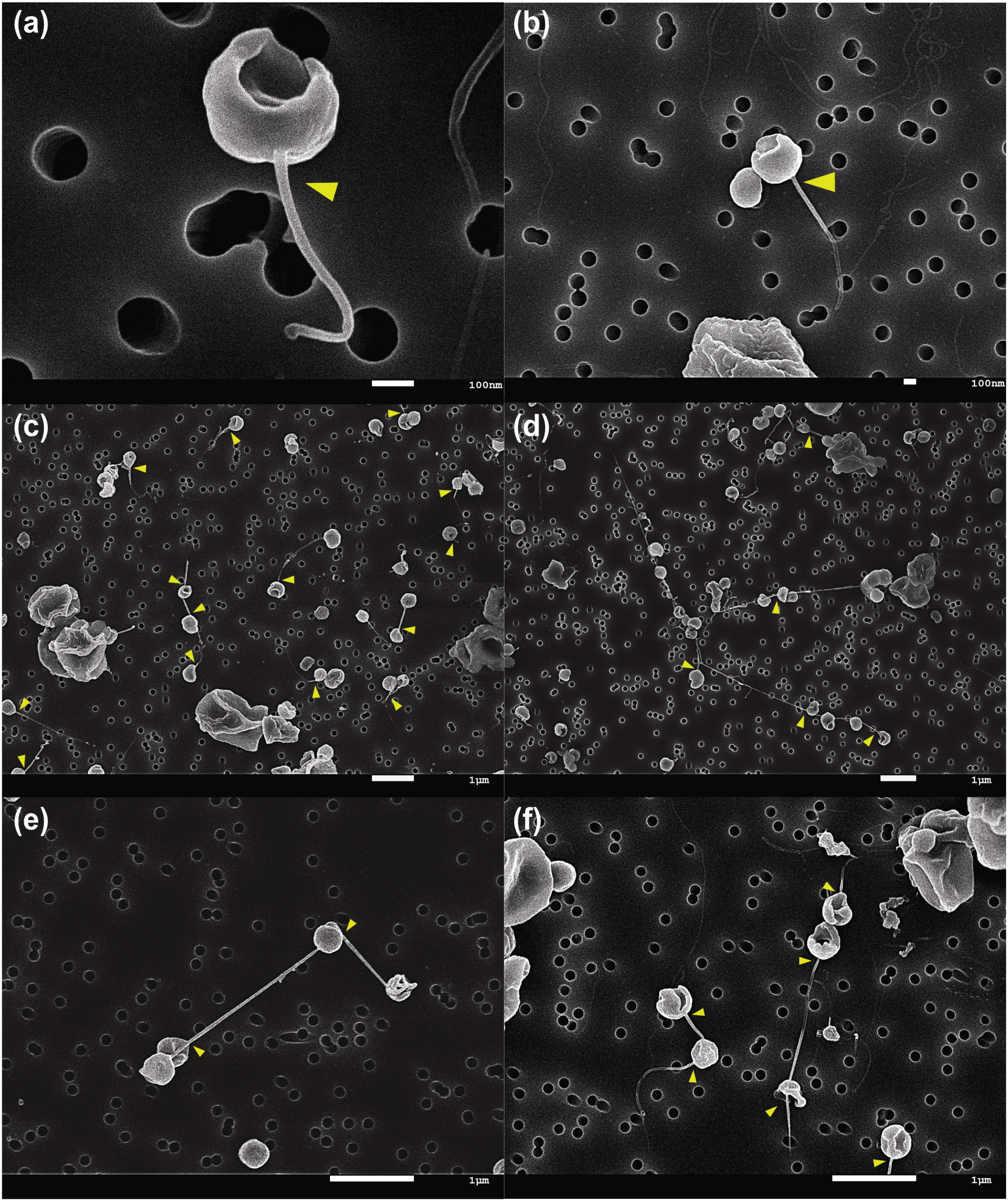


**Extended Data Fig. 3 |** SEM images showing nanotubes emerged from YN4 cells (yellow arrows).
