## Supplemental Data 2 for "A Tripartite Co-culture System Reveals Defensive Mutualism Between a *Nanobdellati* Symbiont and Its Host"

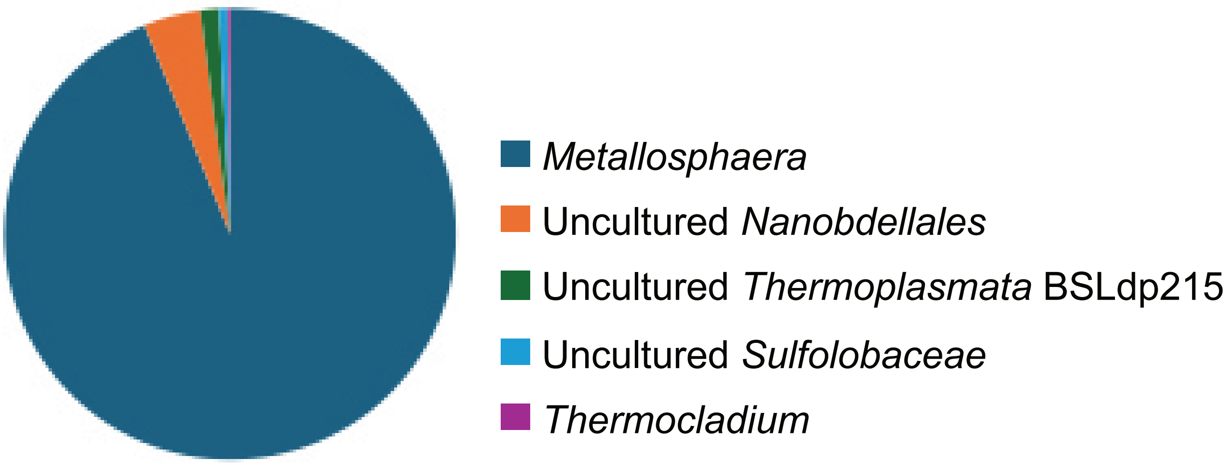


**Supplementary Information Fig. 1.** Microbial community structure of enrichment culture determined by 16S rRNA gene amplicon sequencing.

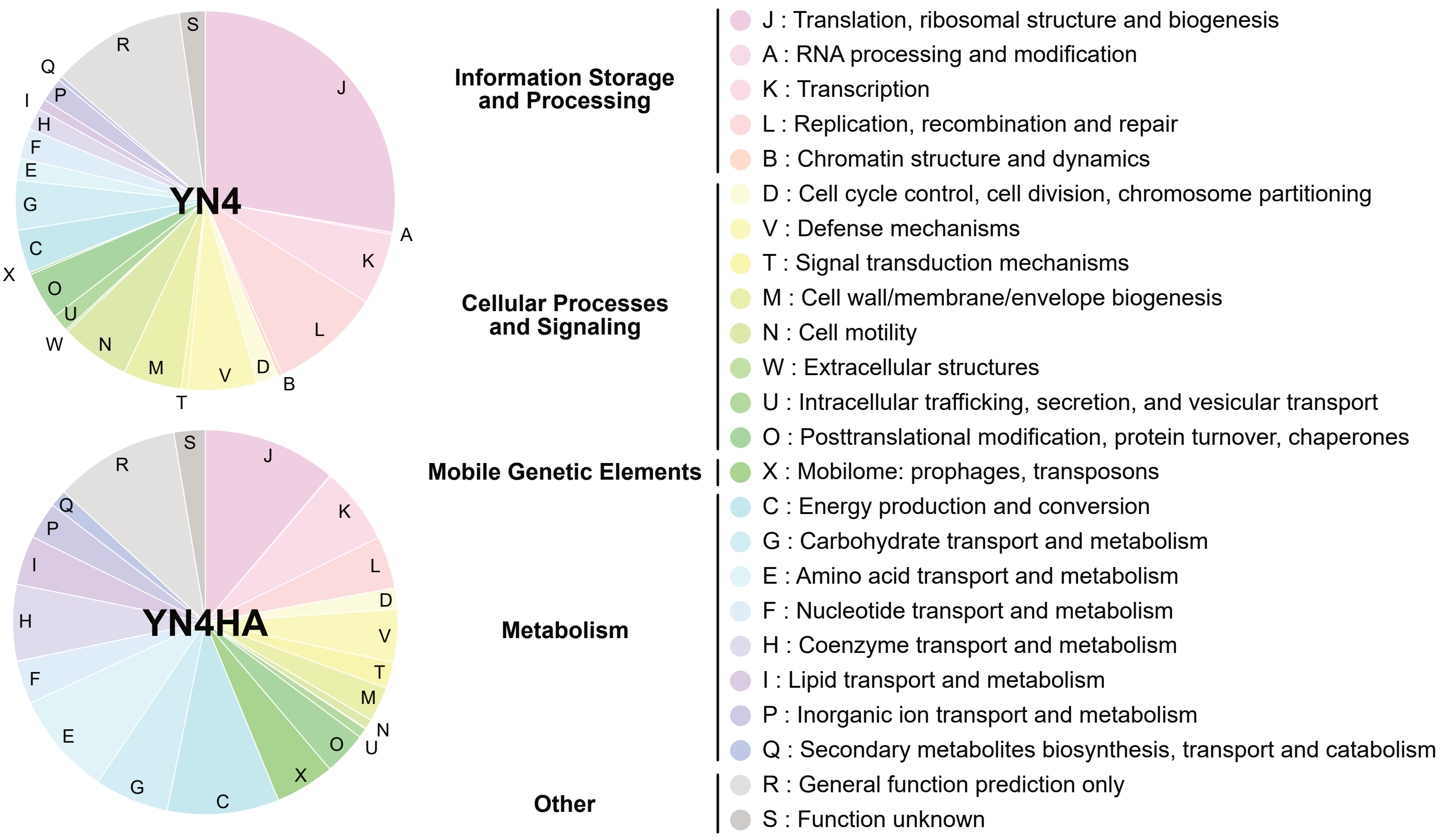


**Supplementary Information Fig. 2.** COG functional categories of genomes of YN4 and YN4HA.

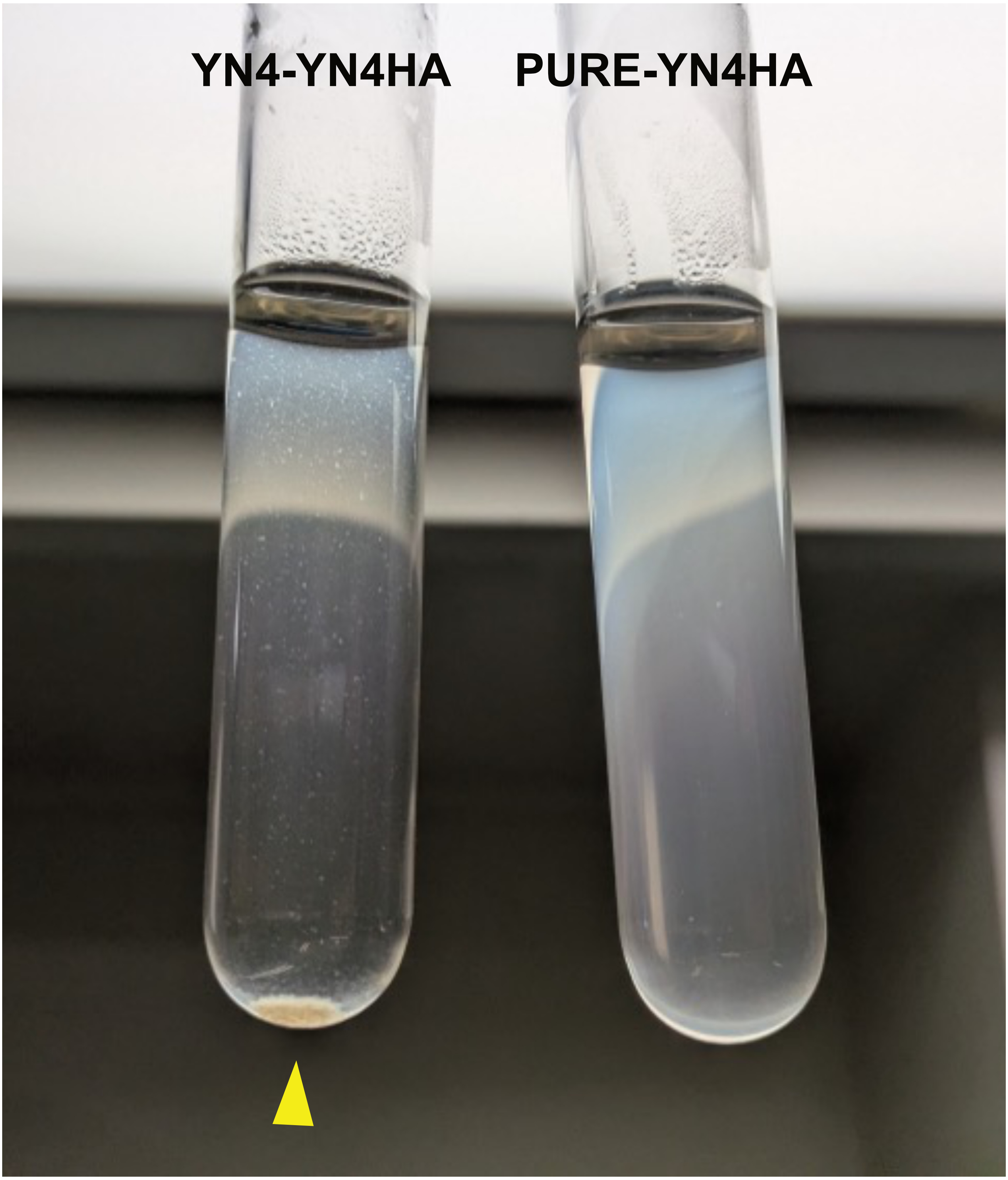


**Supplementary Information Fig. 3.** Culture tubes of the YN4–YN4HA co-culture (left) and the YN4HA pure culture (right). A visible precipitate (yellow arrow) formed in the YN4–YN4HA co-culture, resulting in markedly lower turbidity in the upper portion of the tube. Small flocs are also observed in the YN4–YN4HA co-culture.

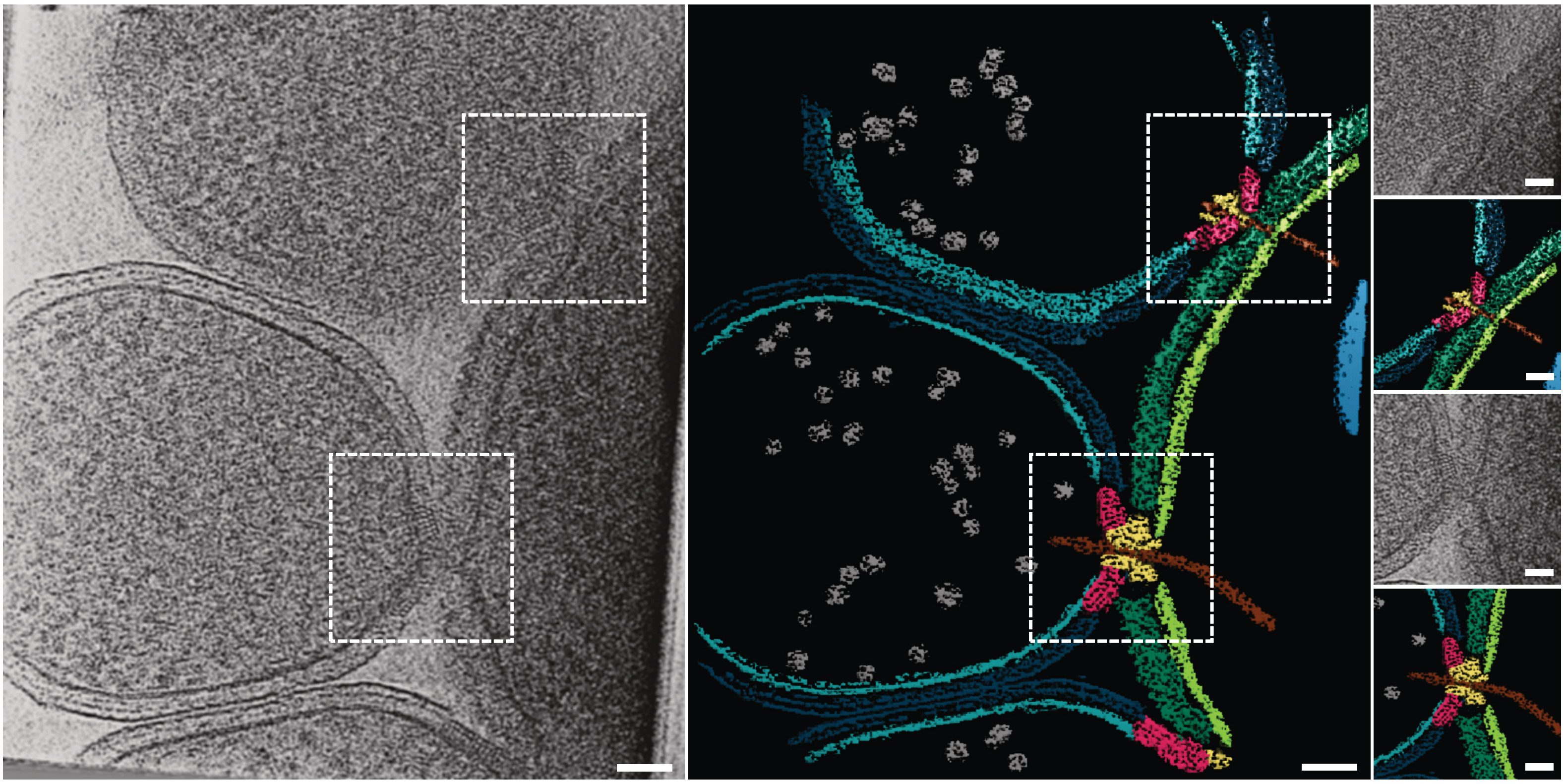

**Supplementary Information Fig. 4.** Multiple YN4-YN4HA interactions. (a) Tomographic slice showing three attachment organelles engaged in interaction with YN4HA, with two of them being in the view and showing the filaments passing through them into the host; (b) segmentation view of the tomographic slice showing the interaction sites;(c) zoomed in view of the top interaction site; (d) Segmentation view of the tomographic slice; (e) zoomed in view of the middle interaction site; (f) segmentation view of the tomographic slice. Segmentation colors are: host membrane green, host S-layer dark green, DPANN membrane teal, DPANN S-layer dark teal, DPANN ribosomes gray, attachment organelle layers red, attachment organelle portal yellow, host granule blue, and filament brown. Scale bars: (first two) 50 nm; (next four) 25 nm.

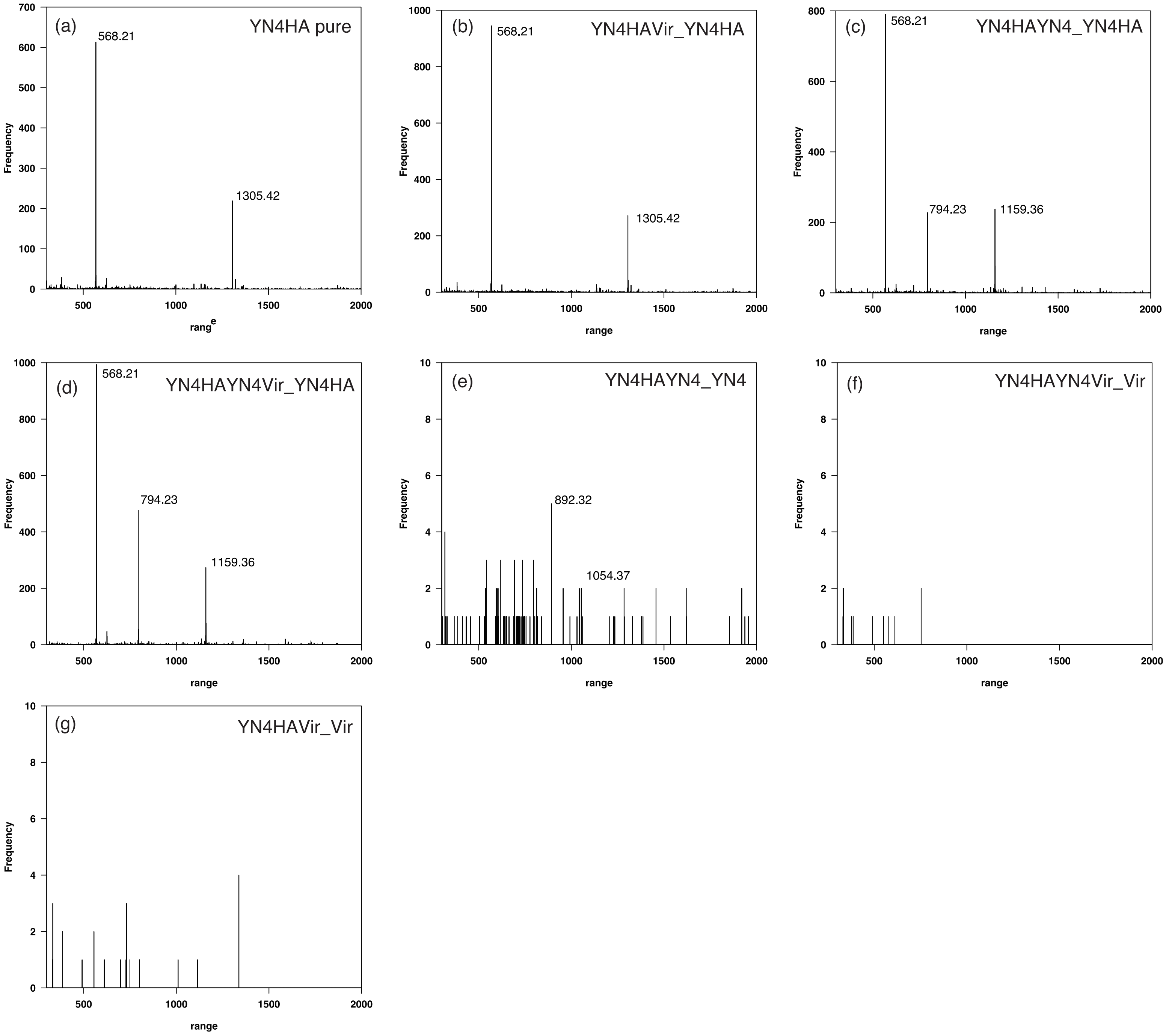


**Supplementary Information Fig. 5.** Delta-mass distribution of glycopeptides in 0.01-Da increments, showing peptide-spectrum matches (PSMs) with modification masses ranging from 300 to 2000 Da.Panels (a–d) represent open-search results for YN4HA, panel (e) for YN4, and panel (f) for MTIV4, derived from the following culture conditions:(a) PURE-YN4HA,(b) MTIV4-YN4HA,(c) YN4-YN4HA,(d) MTIV4-YN4-YN4HA,(e) YN4 in YN4-YN4HA, and(f) MTIV4 in MTIV4-YN4HA.

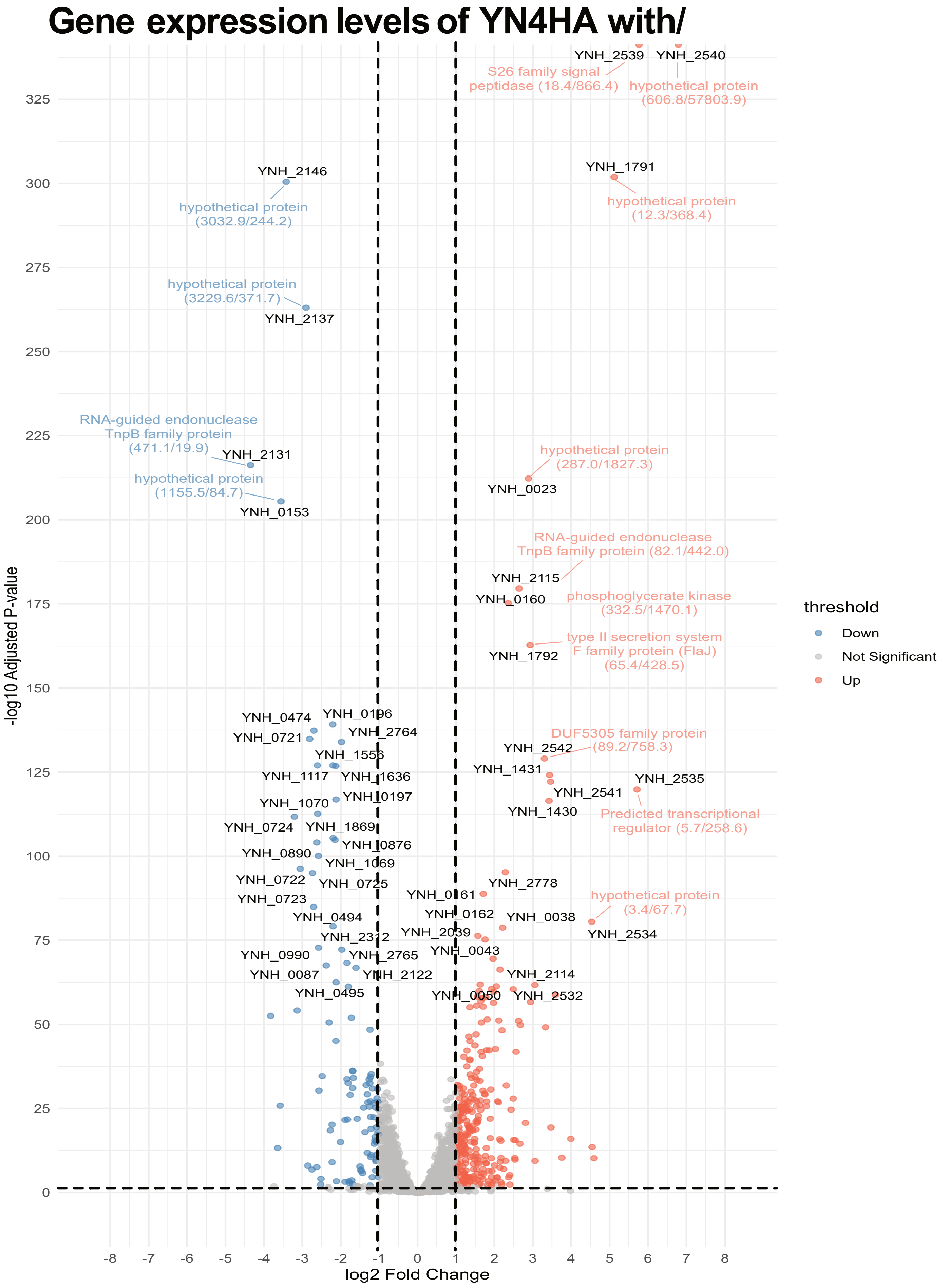


**Supplementary Information Fig. 6.** Volcano plot showing gene expression levels of YN4HA with and without the YN4 symbiont. Thresholds were set at adjusted P-value (padj) < 0.1 and |log2 fold change| > 1. Red and blue dots indicate genes significantly up-regulated and down-regulated, respectively, in the presence of the YN4 symbiont, while gray dots represent genes with no significant change. Some gene names are labeled next to the plots, and TPM values in PURE-YN4HA and YN4-YN4HA are shown in parentheses, respectively.

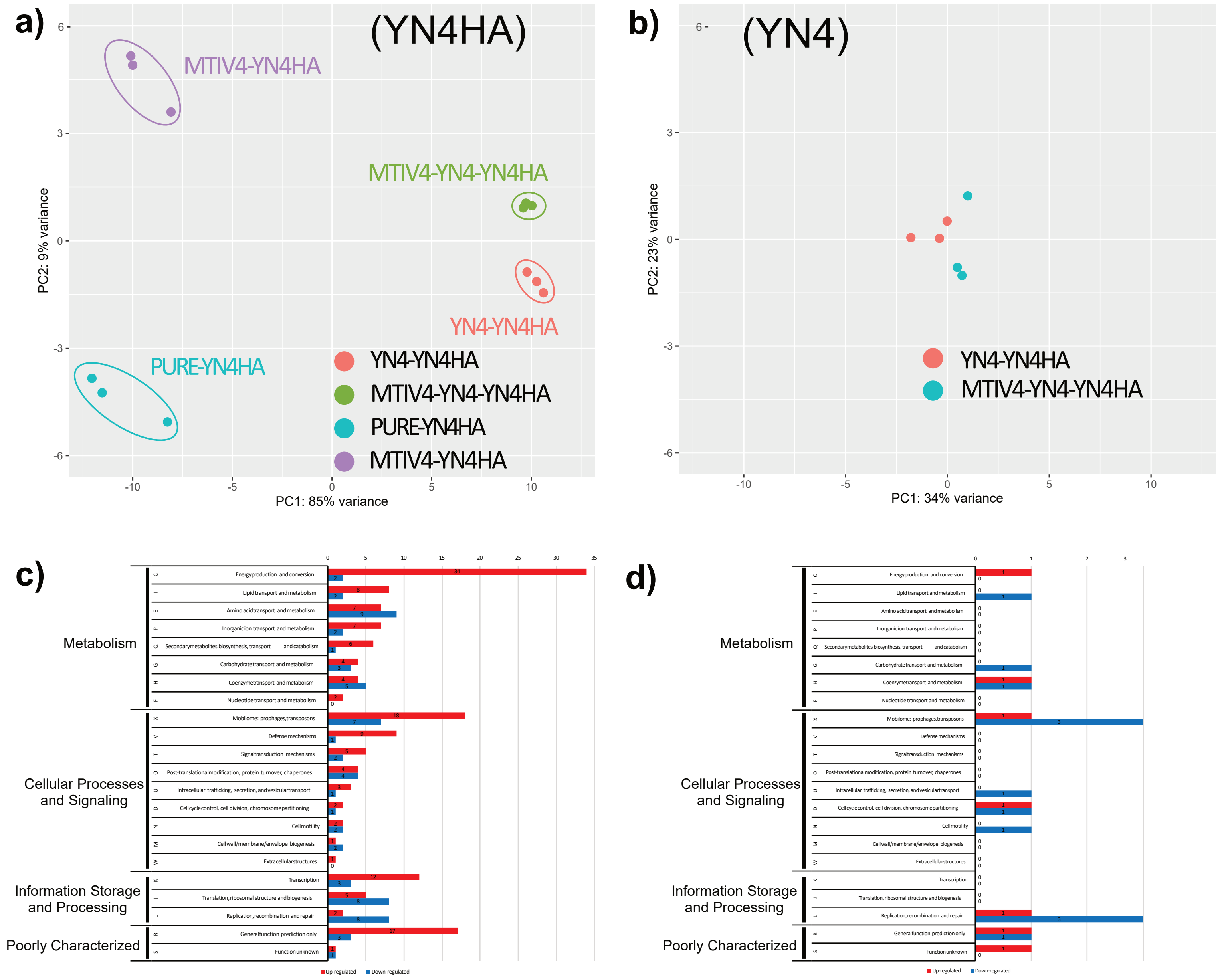


**Supplementary Information Fig. 7.** PCA plots of transcriptomic data for YN4HA (a) and YN4 (b) in each culture condition and the number of upregulated and downregulated genes in YN4HA by the presence of YN4 (c) and MTIV4 (d). The x-axis and y-axis in the PCA represent the first and second principal components, respectively. Red, green, blue, and purple dots indicate the biological triplicates of the following culture groups:YN4-YN4HA, MTIV4-YN4-YN4HA, PURE-YN4HA, and MTIV4-YN4HA, respectively. The upregulated and downregulated genes were extracted from volcano plot in Supplementary Information Figs. 4 and 6, respectively, and classified into the COG categories.

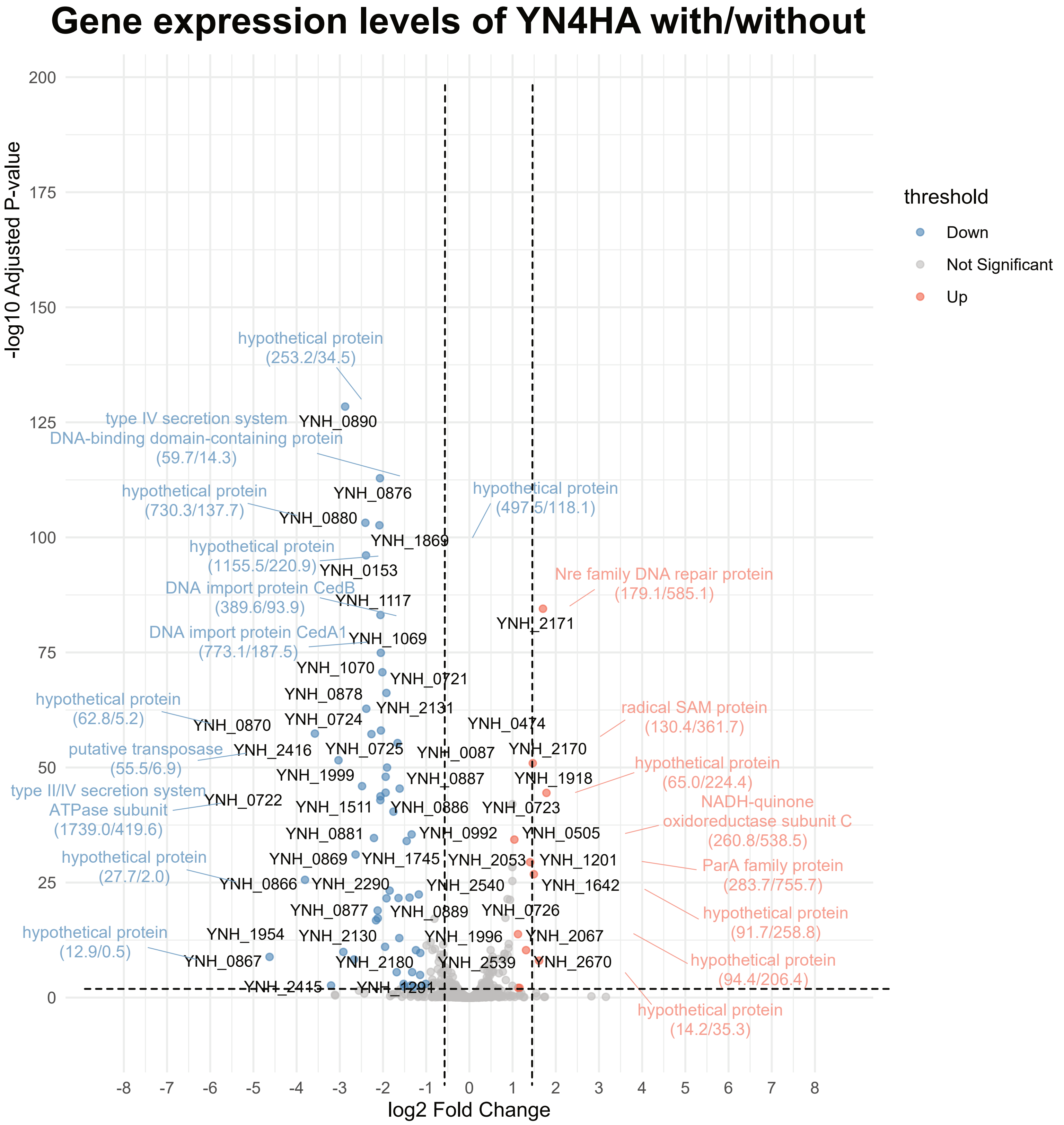


**Supplementary Information Fig. 8.** Volcano plot showing gene expression levels of YN4HA with and without the MTIV4 virus. Thresholds were set at adjusted P-value (padj) < 0.1 and |log2 fold change| > 1. Red and blue dots indicate genes significantly up-regulated and down-regulated, respectively, in the presence of the MTIV4 virus, while gray dots represent genes with no significant change. Some gene names are labeled next to the plots, and TPM values in PURE-YN4HA and MTIV4-YN4HA are shown in parentheses, respectively.

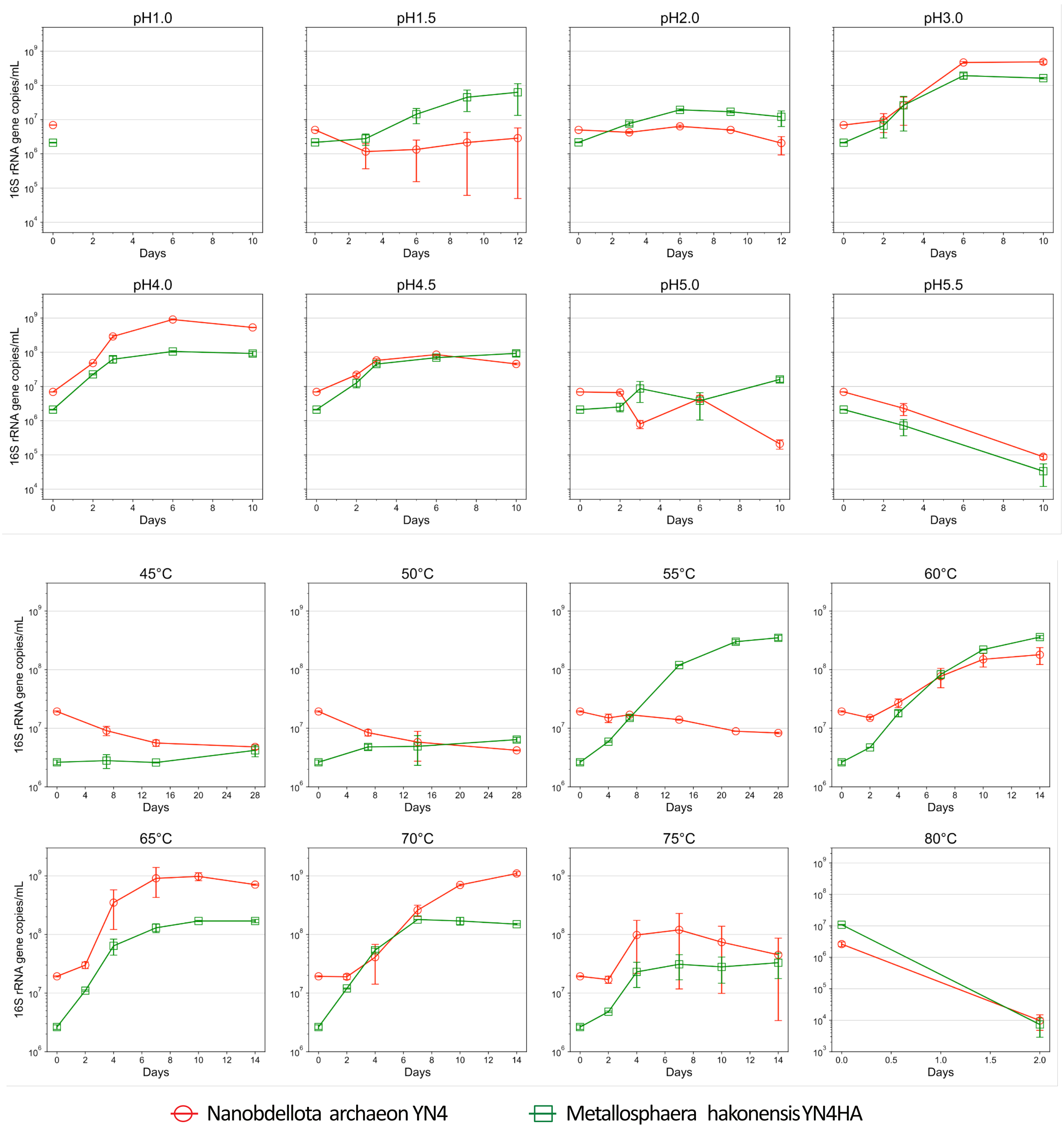


**Supplementary Information Fig. 9.** Growth curves of YN4-YN4HA co-culture at deafferent temperatures and pH values.

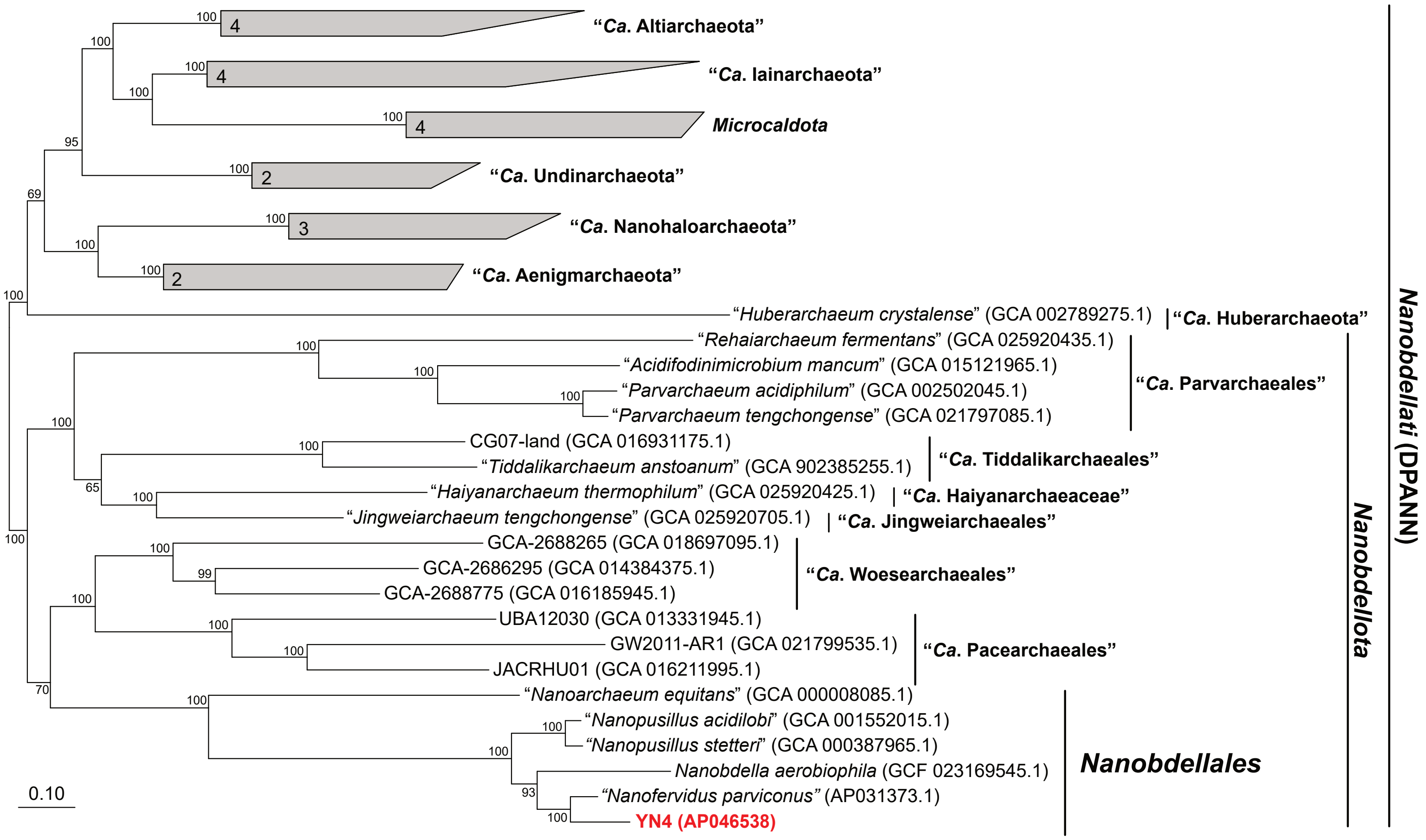


**Supplementary Information Fig. 10.** Phylogenetic position of strain YN4 in the kingdom *Nanobdellati*.

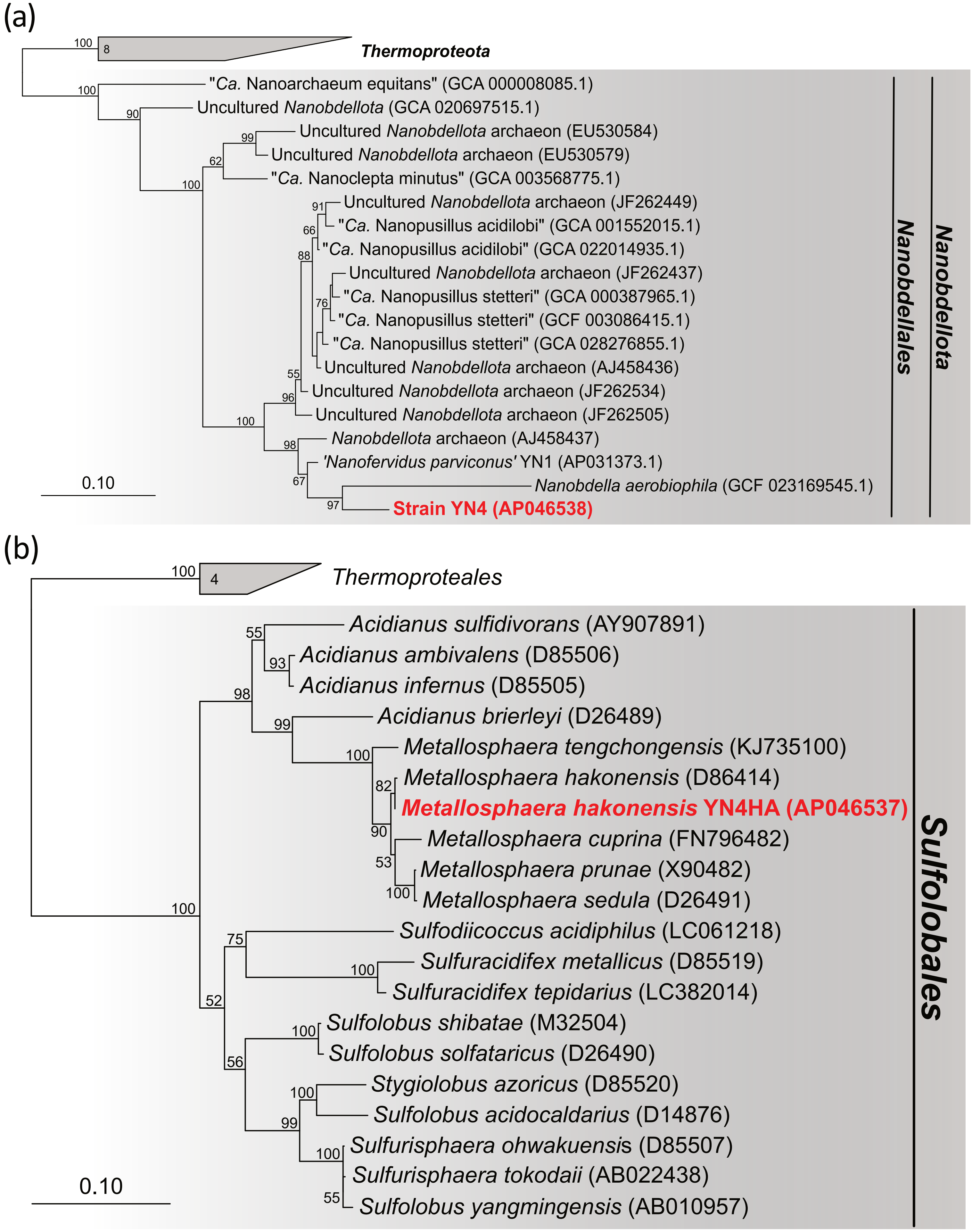


**Supplementary Information Fig. 11.** The 16S rRNA gene phylogenetic trees, showing taxonomic positions of *Nanobdellota* archaeon strain YN4 (a) and its host *Metallosphaera hakonensis* strain YN4HA (b).

### Gene expression levels of YN4 with/without MTIV4 virus

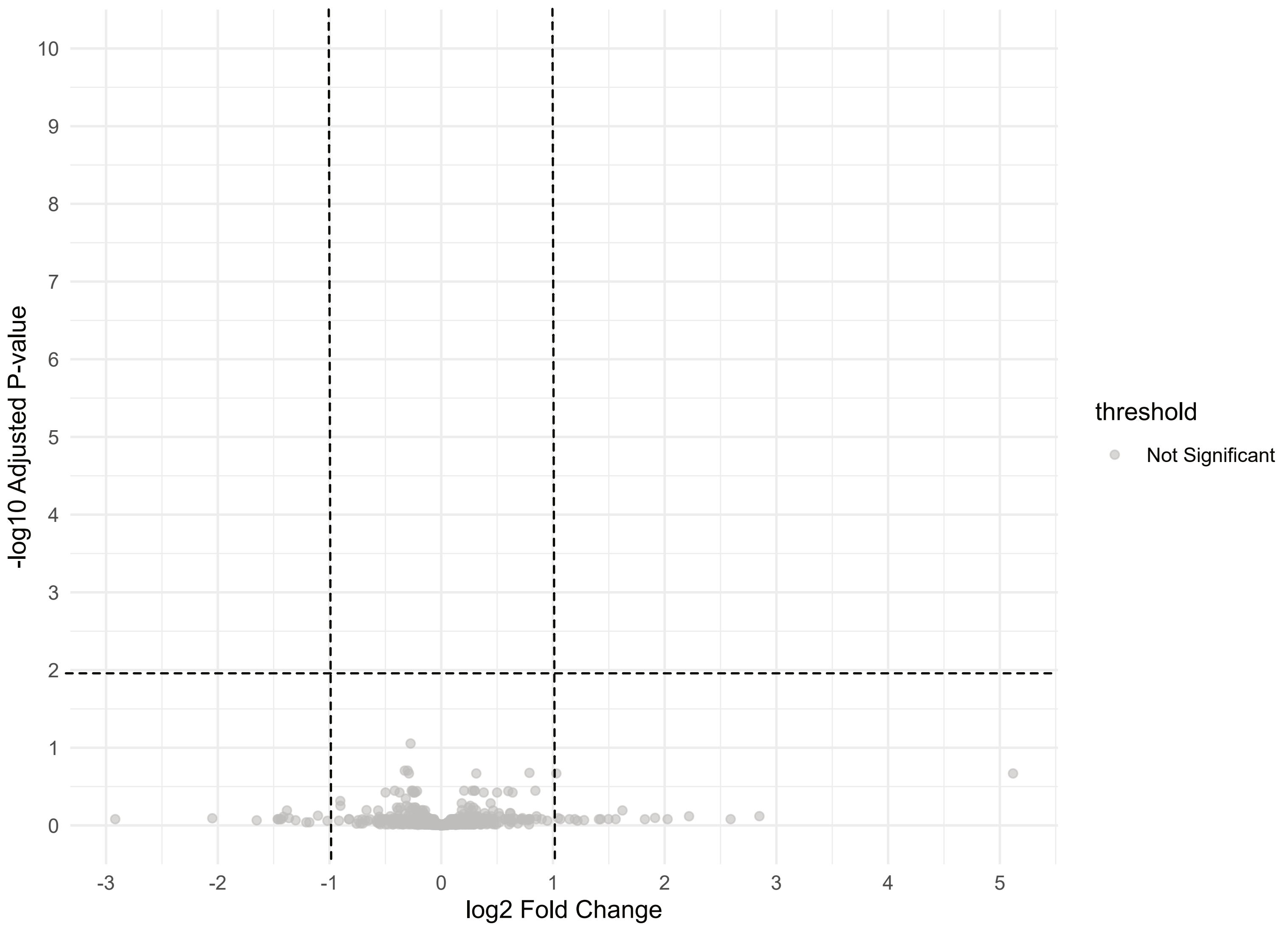


**Supplementary Information Fig. 12.** Volcano plot showing gene expression levels of YN4 with and without the MTIV4 virus (YN4-YN4HA vs MTIV4-YN4-YN4HA). Thresholds were set at adjusted P-value (padj) < 0.1 and |log2 fold change| > 1. Gray dots represent genes with no significant change.

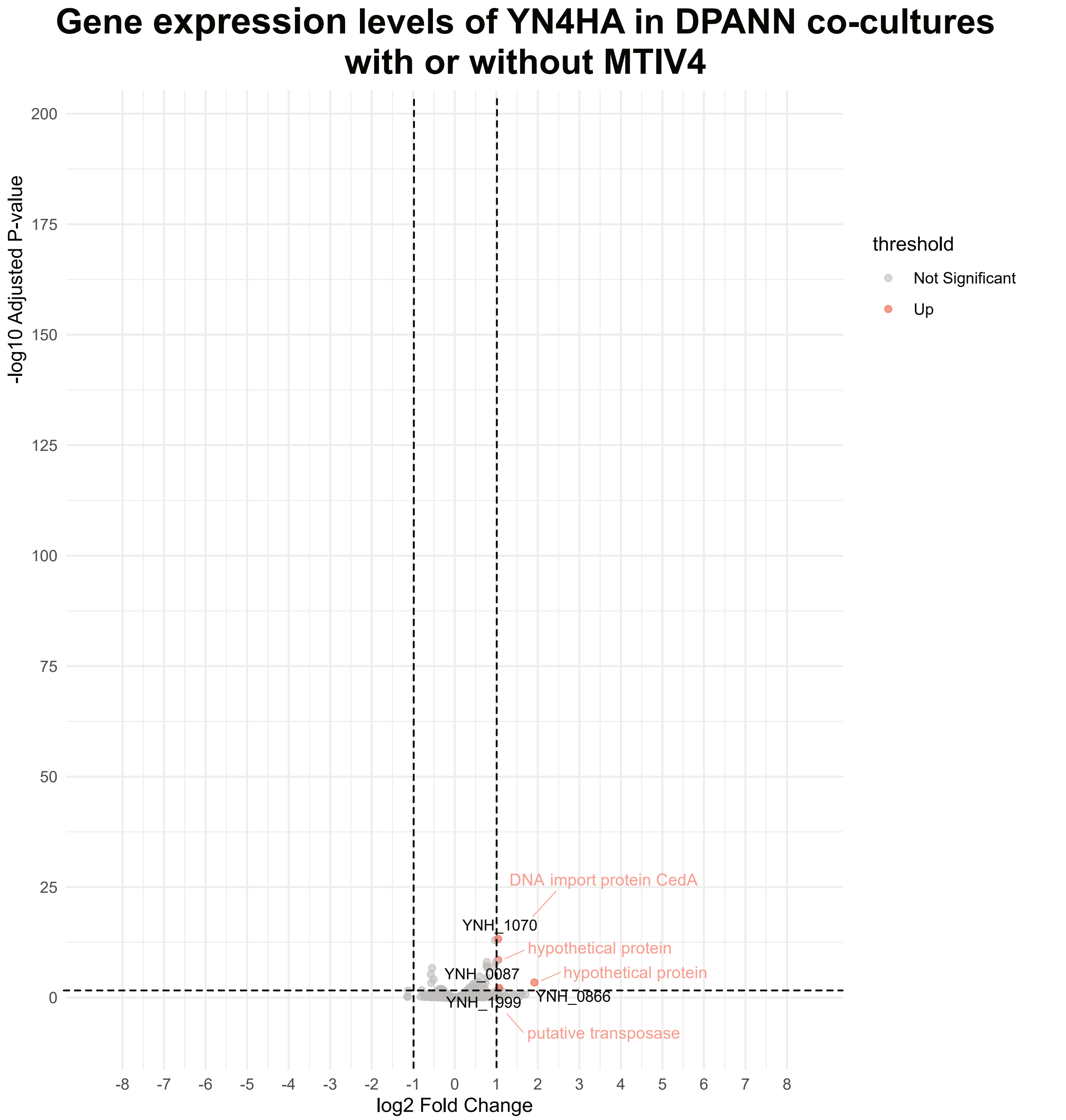


**Supplementary Information Fig. 13.** Volcano plot showing gene expression levels of in DPANN co-cultures with or without MTIV4. Thresholds were set at adjusted P-value (padj) < 0.1 and |log2 fold change| > 1. Red and blue dots indicate genes significantly up-regulated and down-regulated, respectively, in the presence of the YN4 symbiont, while gray dots represent genes with no significant change. Gene names are labeled next to the plots.

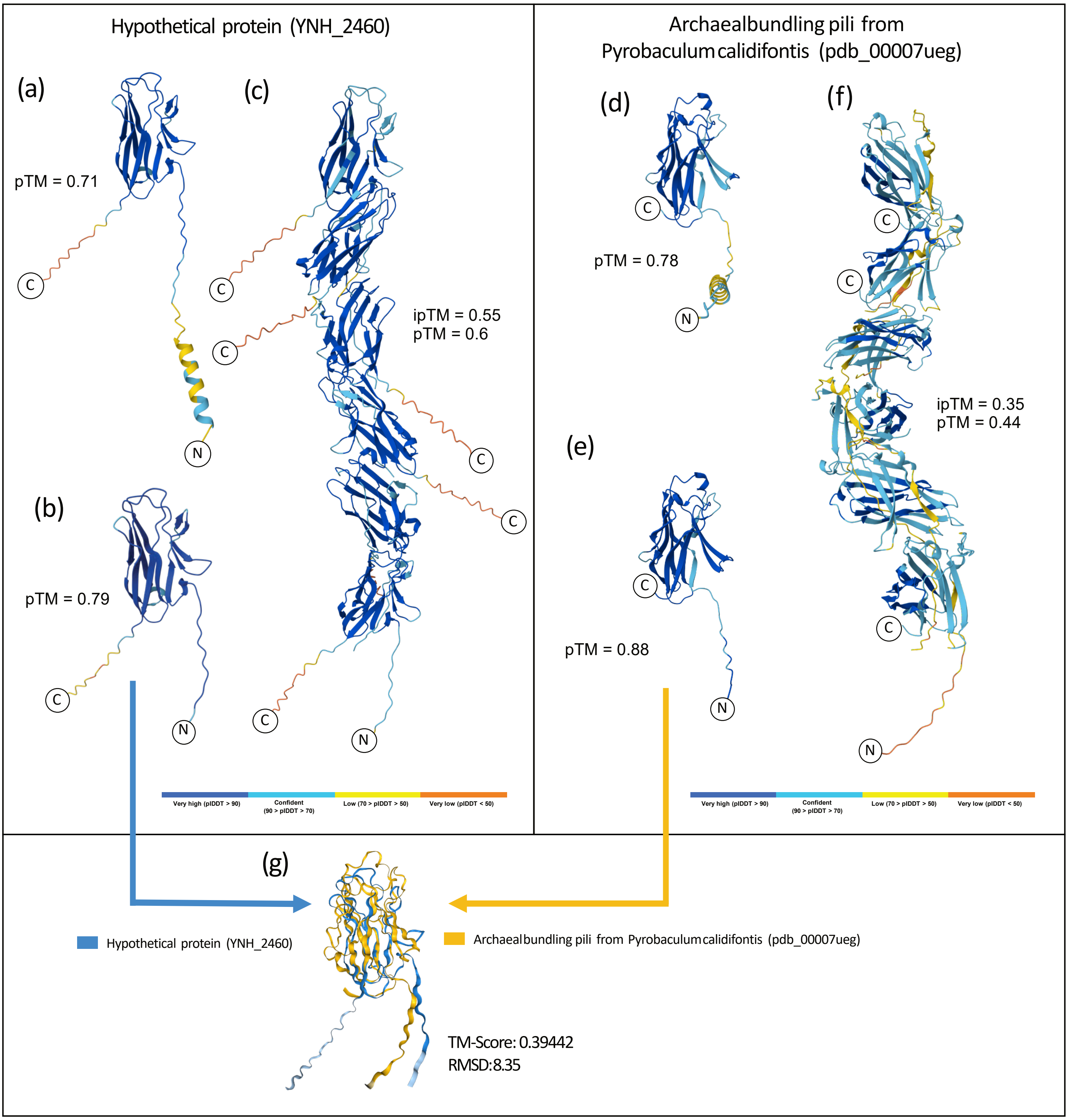


**Supplementary Information Fig. 14. Predicted three-dimensional structures and structural comparison of YNH_2540 and archaeal bundling pilin AbpA from *Pyrobaculum calidifontis*.** (a) Full-length YNH_2540 including the N-terminal α-helical region. (b) Mature YNH_2540. (c) AlphaFold multimer model generated usign six identical mature YNH_2540 proteins. (d) Full-length AbpA. (e) Mature AbpA. (f) AlphaFold multimer model generated using six identical mature AbpA proteins. (g) Structural superposition of YNH_2540 (blue) and AbpA (yellow) (TM-score = 0.394, RMSD = 8.35). Predicted TM-scores (pTM) and inter-chain predicted TM-scores (ipTM) are indicated near each model. Models in (A)-(E) are colored by

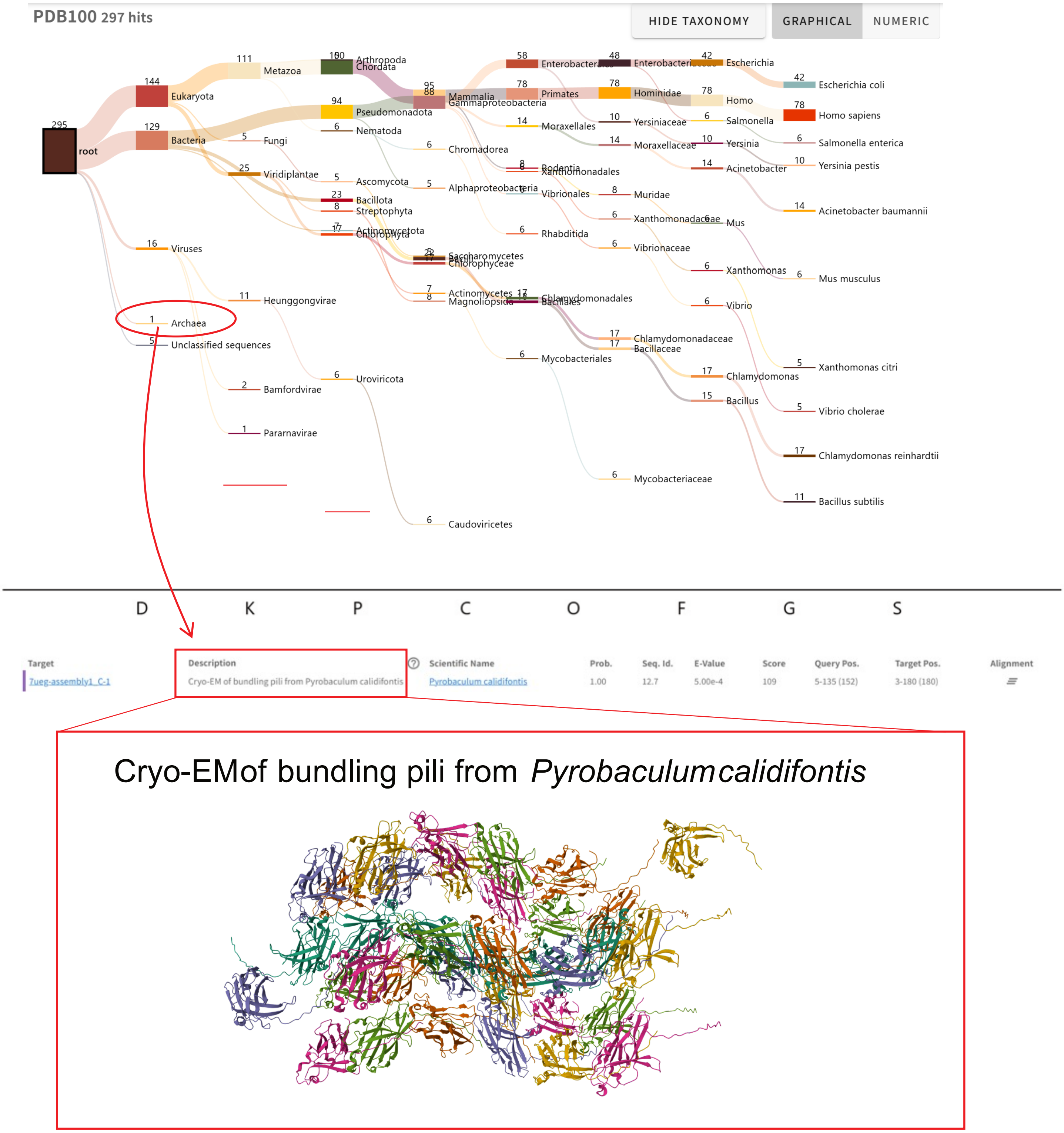


**Supplementary Information Fig. 15. Foldseek search result for YNH_2540 against the PDB100 database.** Taxonomic distribution of Foldseek hits is shown in the Sankey plot (top), indicating that one archaeal hit was detected, corresponding to a Cryo-EM structure of bundling pili from *Pyrobaculum calidifontis*. The corresponding Cryo-EM model, downloaded from the Protein Data Bank (PDB), is shown below.

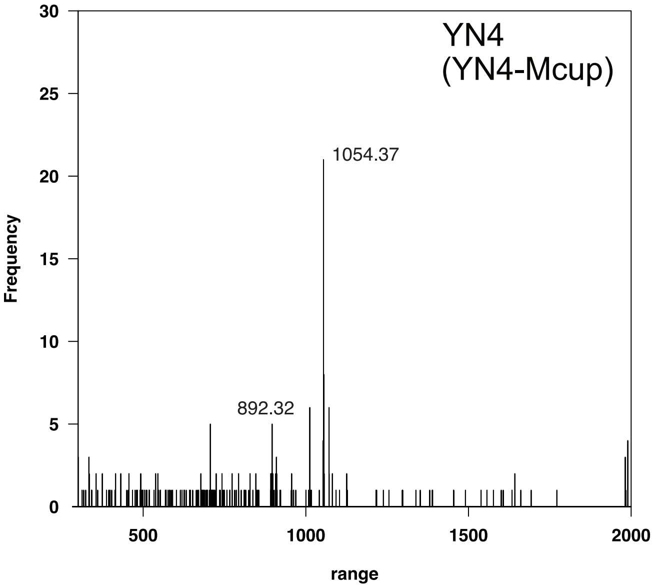


**Supplementary Information Fig. 16.** Delta-mass distribution of glycopeptides in 0.01-Da increments, showing peptide-spectrum matches (PSMs) with modification masses ranging from 300 to 2000 Da for open-search results for YN4 when cultured with *Metallosphaera cuprina*.

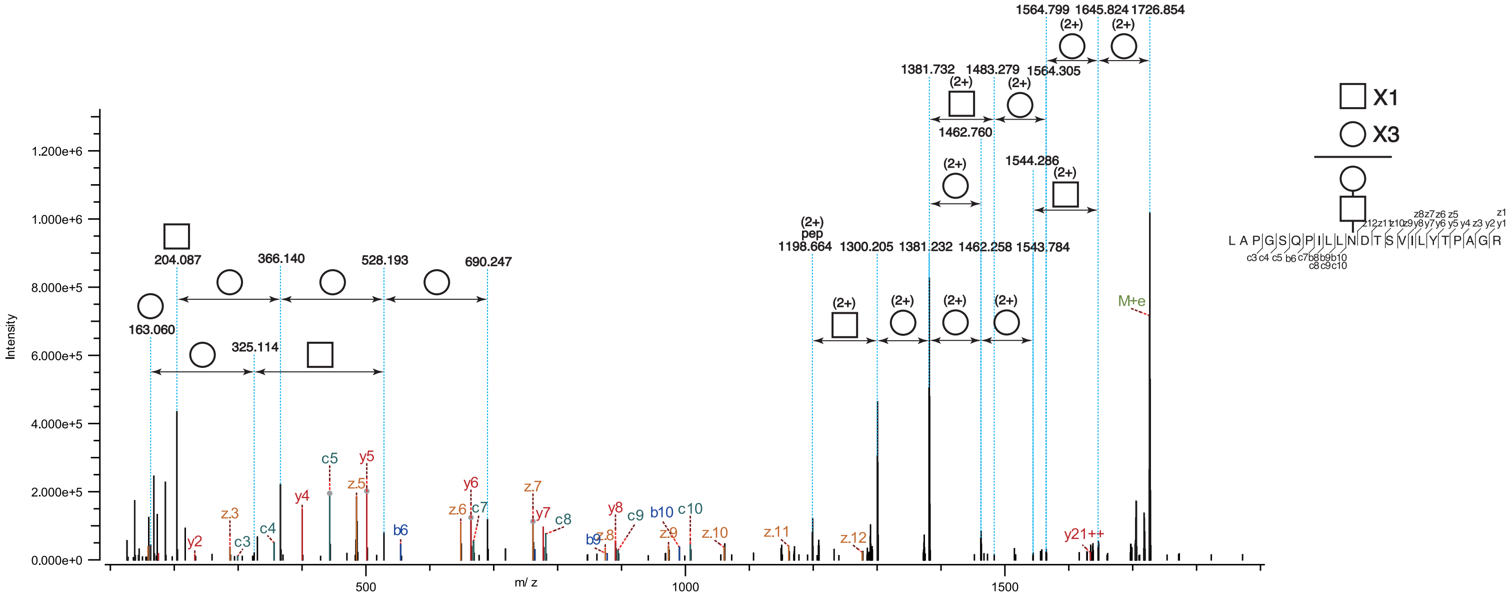



**Supplementary Information Fig. 17. MS/MS spectrum showing glycan assignments on a YN4 protein**. Higher-energy collision dissociation (HCD) spectrum showing the N-glycan structure of 1054.37 Da on archaellin (YN4_015) of strain YN4 when cultured with Metallosphaera cuprina.

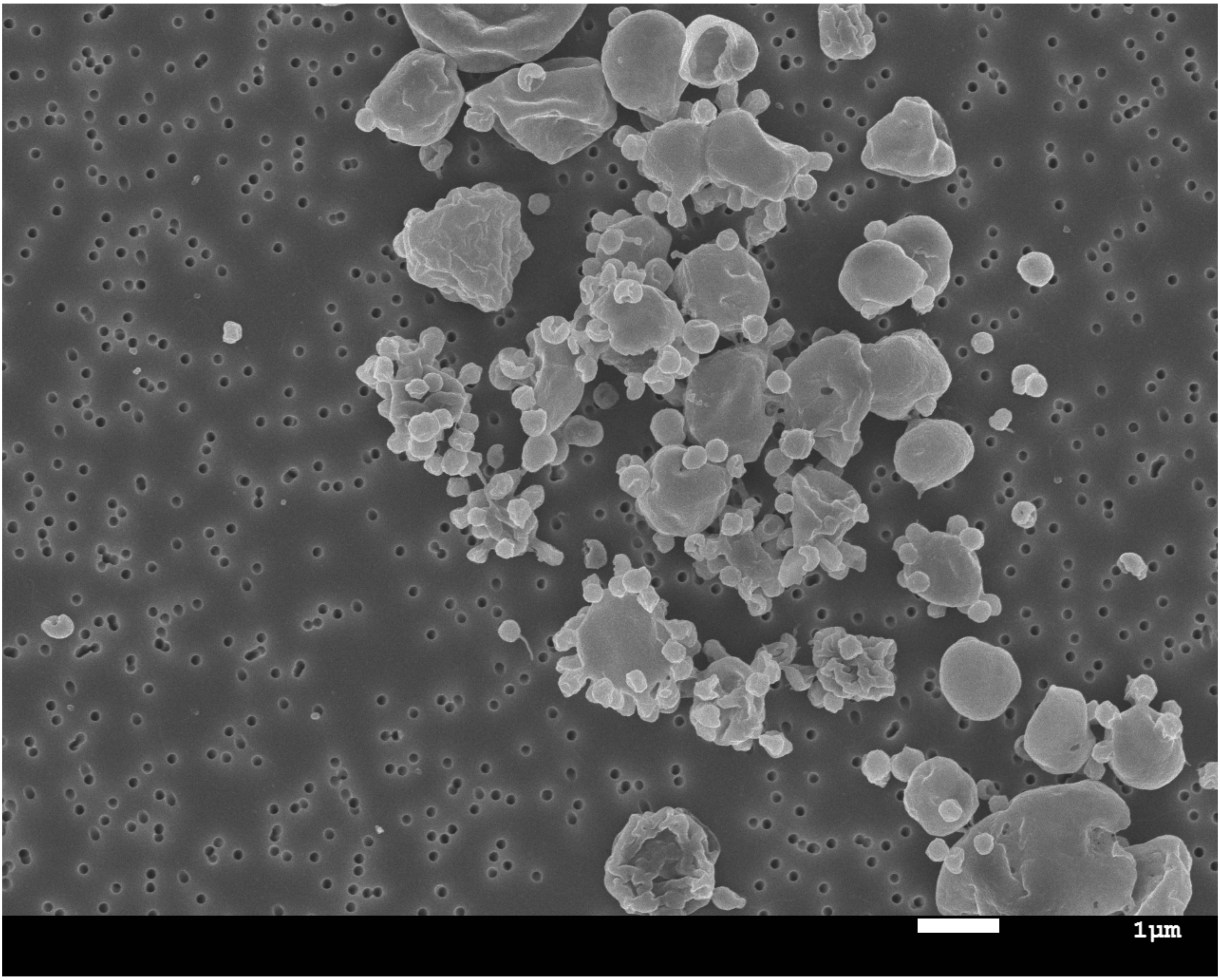


**Supplementary Information Fig. 18.** SEM image showing typical cells of YN4 and YN4HA.
