## Supplementary Text for "A Tripartite Co-culture System Reveals Defensive Mutualism Between a *Nanobdellati* Symbiont and Its Host"

### Elimination of MTIV4 via Host Switching and Reconstruction of a Virus-free Co-culture

By repeatedly conducting a host-switching experiment using *M. cuprina* JCM15769$^{T}$ as an alternative host for the *Nanobdellales* archaeon YN4 (Extended Data Fig. 1c,d,e), we obtained a co-culture of *M. cuprina* and YN4 free of MTIV4. This result indicates that MTIV4 cannot infect *M. cuprina,* whereas YN4 can grow on *M. cuprina* as an alternative host of YN4HA. Using the YN4 and *M. cuprina* co-culture, we again performed a host-switching experiment to re-switch the host from *M. cuprina* to YN4HA in the absence of MTIV4. As a result, a pure co-culture of YN4 and YN4HA, free of MTIV4, was successfully obtained (Extended Data Fig. 1f).

### Genome sequencing of the YN4-YN4HA-MTIV4 co-culture

The total numbers of raw reads, subsampled reads, and quality-filtered reads from Illumina and Nanopore sequencing of the YN4-YN4HA-MTIV4 co-culture are summarized in Supplementary Table S12. Following genome assembly, two circular genomes (758,205 bp and 2,631,175 bp) and one short linear contig (13,941 bp) were obtained. The two circular genomes were identified as a *Nanobdellales* archaeon and *Metallosphaera hakonensis*, respectively, based on their phylogenetic positions as described below. The linear contig was identified as *Metallosphaera* Turreted Icosahedral Virus 4 (MTIV4) (see below in detail). More than 99.9% of quality-filtered short and long reads mapped to the three genomes (Supplementary Table S13), indicating that this system consists solely of the two archaeal strains, YN4 and YN4HA, together with the virus, MTIV4.

### Additional Genomic Features of YN4

Genes related to oxidative stress (i.e., *ahpC*, *cydA*, and *SOD*), as well as the reverse gyrase gene (a marker for hyperthermophiles [1]), were identified, consistent with YN4's growth under oxic, high-temperature conditions. However, as the respiratory system is not encoded in the genome, aerobic respiration is unlikely to occur in YN4. Several transporter-related genes were detected, which may enable the uptake of essential nutrients from the surrounding environment. A CRISPR/Cas system with an array of 18 spacers and restriction-modification systems (types I and III) were identified in the genome of YN4, indicating that this strain possesses immune systems against certain viruses. A BLASTN search of the 18 spacers against the NCBI-nt database yielded no significant hits, suggesting the presence of as-yet-uncharacterized viruses that remain undetected in current metagenomic datasets from this environment.

**Basic growth characteristics of YN4**

Using the YN4-YN4HA co-culture, growth temperature and pH ranges for YN4HA and YN4 were examined (Supplementary Information Fig. 9). *M. hakonensis* YN4HA grew at 45–75°C (optimum: 65°C) and pH 1.5–5.0 (optimum: 3–4), whereas *Nanobdellales* archaeon YN4 grew at 60–75°C (optimum: 65°C) and pH 1.5–4.5 (optimum: 3–4). Compared with YN4HA, YN4 exhibited a narrower range of growth temperatures and pH values. This pattern has also been reported in other *Nanobdellales* species [2,3], suggesting that a narrow range of growth temperature and pH is a common feature within the order *Nanobdellales* in the phylum *Nanobdellota*. In all cases where YN4 grew, its growth consistently followed that of YN4HA (Supplementary Information Fig. 9), suggesting that actively growing host cells, rather than dead cells, are important for the growth of YN4.

**Host specificity of *Nanobdellales* archaeon YN4**

Consistent growth of YN4 on day 7 was observed in two consecutive co-culture experiments with *M. hakonensis* YN4HA (original host), *M. hakonensis* DSM7519$^T$, and *M. cuprina* JCM15769$^T$. In contrast, no growth of YN4 was observed with the following thermoacidophilic species: *Acidianus brierleyi* DSM1651$^T$, *M. javensis* AS-7, *M. sedula* DSM5348$^T$, *Saccharolobus caldissimus* HS-3$^T$, *Scl. shibatae* DSM5389$^T$, *Scl. solfataricus* JCM8930$^T$, *Sulfodiicoccus acidiphilus* HS-1$^T$, *Sulfolobus acidocaldarius* DSM639$^T$, *Sulfuracidifex tepidarius* JCM16833$^T$, *Saf. metallicus* DSM6482$^T$, *Sulfurisphaera javensis* KD-1$^T$, *Sfs. ohwakuensis* TA-1$^T$, and *Thermoplasma acidophilum* JCM9062$^T$. These results indicate that YN4 has a narrow host range, limited to the genus *Metallosphaera*. A narrow host range has been reported in other *Nanobdellales* species (i.e., *Nanoarchaeum equitans*, *Nanobdella aerobiophila*, and "*Ca*. Nanofervidus parviconus"); however, in these cases, host specificity was limited to a single species or, often, to a single strain [2–4]. In contrast, YN4 demonstrated that the original host can be replaced by another strain, *M. hakonensis* DSM7519$^T$, and by a different species, *M. cuprina* JCM15769$^T$. This suggests that the host range of *Nanobdellales* is not necessarily limited to a single species. A relatively broad host range has also been reported in the phylum *Microcaldota* (formerly "*Ca*. Micrarchaeota"), another member of the *Nanobdellati* (formerly DPANN) [2,5], whose host species also belong to the order *Sulfolobales* [6]. Since members of the order *Sulfolobales* are among the most abundant and globally distributed taxa in terrestrial acidic hot springs [7,8], other *Nanobdellati* archaea utilizing *Sulfolobales* species as their hosts might also inhabit the same environments. Enrichment culture-based cultivation targeting *Sulfolobales* species, although conventional, may still be effective for isolating other cultivable *Nanobdellati* species, as exemplified by our recent consecutive studies [2,3,6]. No growth was observed in the triplicate negative controls inoculated with the 0.45 μm filtrate of the YN4-YN4HA co-culture, indicating that YN4 cells do not grow without host cells, underscoring its obligatory symbiotic characteristic.

**Phylogenetic position of *Nanobdellales* archaeon YN4**

The genome tree based on 53 archaeal marker protein sequences is shown in Supplementary Information Fig. 10. Strain YN4 is placed within the order *Nanobdellales* of the phylum *Nanobdellota* [2,5,9]. A similar topology was observed in both the concatenated single-copy protein phylogenetic tree and the 16S rRNA gene phylogenetic tree (Extended Data Fig. 2 and Supplementary Information Fig. 11a), indicating that this phylogenetic position is well supported. The closest relative of YN4 is '*Nanofervidus parviconus*,' recently isolated alongside its host, *Sulfurisphaera ohwakuensis* from the same hot spring [3], followed by *Nanobdella aerobiophila* [2]. Both related species grow under oxic conditions, and their hosts are *Sulfolobales* species [2,3], suggesting the presence of a group within *Nanobdellales* that grows under oxic conditions. The ranges of amino acid identity (AAI) and 16S rRNA gene sequence identity among cultivated *Nanobdellales* archaea are 49.8–80.5% and 80.6–93.0%, respectively (Supplementary Tables 14 and 15). The 16S rRNA gene sequence identity values are below the widely used genus-level threshold of 94.5% [10], suggesting that YN4 represents at least a novel genus within the order of *Nanobdellales*. Based on the 16S rRNA gene similarity values and its position in the phylogenetic trees (Extended Data Figs. 2; Supplementary Information Figs. 10 and 11a), we propose the name '*Nanomutabilis hospitiaugens*' to accommodate strain YN4.

**General transcriptomic profiles of YN4HA, YN4, and MTIV4**

General transcriptomic profiles, normalized as transcripts per million (TPM) for YN4HA (in PURE-YN4HA), YN4 (in YN4-YN4HA), and MTIV4 (in MTIV4-YN4HA) are summarized in Supplementary Tables 16-18.

In the pure culture of *Metallosphaera hakonensis* YN4HA, the most highly expressed genes encoded S-layer proteins (SlaA and SlaB) (Supplementary Table 16). Other abundant transcripts included those related to energy metabolism (e.g., cytochrome oxidases, ferredoxin), protein folding (thermosome subunits), oxidative stress responses (e.g, rubrerythrin, peroxiredoxin, thioredoxin), and translation (e.g., elongation factor EF-1α, Alba). Several membrane- and transporter-associated proteins (e.g., cytochrome oxidase subunits, GlcS, and ABC transporters) were also highly expressed, reflecting the active growth of YN4HA, and corresponding well with the growth curve (Fig. 5A, Day 4).

In the transcriptome of strain YN4 (in YN4–YN4HA), the most highly expressed gene encoded an S-layer–like protein (YN4_391; rank 1), followed by archaellin (ArlB; rank 2) (Supplementary Table 17). Two copies of the genes for cell division protein FtsZ (YN4_420 and YN4_686; ranks 13 and 21) were also highly expressed, suggesting that YN4 retains an FtsZ-based cell division machinery. Core genes involved in translation, transcription, and protein folding and degradation (e.g., elongation factor EF-1α, histones, thermosome, proteasome) were strongly expressed, reflecting active cellular processes. Several glycolysis/gluconeogenesis-related genes, notably the highly expressed fructose-1,6-bisphosphate aldolase/phosphatase (YN4_477; rank 3) and NADP-dependent glyceraldehyde-3-phosphate dehydrogenase (YN4_765; rank 18), were found to be expressed. As the glycolytic/gluconeogenic pathway is incomplete in YN4, these genes may function in carbon recycling rather than in energy production. Many of the other highly expressed genes were hypothetical, and may be related to unique characteristics of YN4, such as host recognition or parasitic/symbiotic interactions.

The transcriptomic profile of MTIV4 was dominated by structural and hypothetical genes, with virion structural proteins (MTIV4_16–18, MTIV4_21) among the most highly expressed (Supplementary Table 18). This expression pattern indicates that MTIV4 was in a late stage of its replication cycle, actively assembling virion components, while the growth of YN4HA appeared slightly suppressed (Fig. 3b, Day 4).

**Principal component analyses of gene expression profiles of YN4HA/YN4**

To investigate the impact of YN4 (symbiont) or MTIV4 (virus) on the transcriptomic profile of the host archaeon YN4HA or YN4, RNA-Seq analysis was performed on the following pure and co-culture systems: YN4HA, YN4-YN4HA, MTIV4-YN4HA, and MTIV4-YN4-YN4HA.

Principal component analysis (PCA) of gene expression profiles from *Metallosphaera hakonensis* YN4HA revealed distinct clustering across all pure and co-culture conditions, with each biological triplicate forming a separate cluster (Supplementary Information Fig. 7a). These results indicate that the transcriptomic profile of YN4HA is strongly influenced by the presence of either the *Nanobdellales* symbiont or the virus. Notably, co-culture with YN4 or MTIV4 resulted in a clear separation from the YN4HA pure culture in PCA space (Supplementary Information Fig. 7a), underscoring the distinct transcriptional impact of each interaction.

In contrast, PCA of the YN4 symbiont gene expression profiles did not show clear separation between co-cultures with and without MTIV4 (Supplementary Information Fig. 7b). Furthermore, volcano plot analysis revealed no significantly upregulated or downregulated genes in YN4 across co-cultures with and without MTIV4 (Supplementary Information Fig. 7b), indicating that the transcriptomic profile of YN4 is not significantly affected by the presence of MTIV4.

The results above reveal an apparent discrepancy: although MTIV4 altered the gene expression profile of YN4HA (Supplementary Information Fig. 7a), no corresponding effect was observed in YN4 (Supplementary Information Figs. 7b and 12). This is unexpected, as transcriptional changes in the host would be expected to influence the gene expression profile of its symbiont as well. One plausible explanation is that the YN4 symbiosis exerts a dominant effect on YN4HA, which may mask the transcriptional influence of MTIV4 and thereby result in no significant changes in the YN4 transcriptome. Indeed, volcano plot analysis comparing YN4HA gene expression between the YN4-YN4HA and MTIV4-YN4-YN4HA co-cultures revealed only four significantly upregulated genes, all with fold changes or adjusted p-values (padj) close to the significance threshold (Supplementary Information Fig. 13). This observation is consistent with the tight clustering of PCA plots for YN4HA in the YN4-YN4HA and MTIV4-YN4-YN4HA conditions (Supplementary Information Fig. 7a). This limited change in gene expression—reflected by only four marginally upregulated genes—further supports the idea that the transcriptional impact of MTIV4 is effectively masked by YN4. If such transcriptional dominance by *Nanobdellati* symbionts also occurs in natural environments, it suggests that their influence on host cellular physiology may far exceed that of viral agents, underscoring their potential ecological importance. However, this relationship may vary depending on the type of virus involved, such as lytic, chronic, or persistent infections. Further studies involving different types of viruses in the context of *Nanobdellati* symbiosis will be necessary to clarify these dynamics.

**A hypothetical protein YNH_2540 resembling AbpA**

Interestingly, a hypothetical protein (YNH_2540) was marked upregulated (106.3-fold) in *Metallosphaera hakonensis* YN4HA during co-cultured with its symbiont YN4 and was the most abundantly expressed transcript (rank 1) in the YN4-YN4HA co-culture (Supplementary Table 19). This protein was also identified as the most abundant in the co-culture, based on the NSAF value (Supplementary Table 20), suggesting an important role in the symbiotic interactions between YN4 and YN4HA. Structural prediction using AlphaFold revealed that YNH_2540 comprises an N-terminal α-helical region followed by a β-sandwich domain. The β-sandwich domain was modelled with very high confidence (pLDDT > 90), whereas the N-terminal helix showed moderate to low confidence scores (pLDDT 50–90) (Supplementary Information Fig. 14). The predicted TM-score (pTM) for the overall model was 0.71, indicating moderate confidence in the global domain arrangement. While TMHMM predicted the N-terminal helix to be a transmembrane segment, both Phobius and LipoP identified it as a Sec-type signal peptide. LipoP further predicted a signal peptidase cleavage site between residues 26 and 27 (VVSLG|FTYTT), suggesting that the mature protein is secreted into the extracellular or periplasmic space. The C-terminal region was predicted to adopt an immunoglobulin-like (Ig-like) β-sandwich fold (Supplementary Information Figs. 14a,b,c). Ig-like domains have been identified in surface-associated proteins of archaea, such as the archaellum-associated protein ArlF (formerly FlaF) in *Sulfolobus acidocaldarius*, and type IV pilins in *Saccharolobus solfataricus* and *Pyrobaculum arsenaticum*, where they are suggested to participate in interactions with the S-layer or viral receptors [11,12]. Coiled-coil prediction using

DeepCoil2 did not detect any coiled-coil motifs, suggesting that the protein is unlikely to undergo coiled-coil-mediated oligomerization, as found in SlaB [13]. If oligomerization occurs, it may instead be mediated by hydrophobic interactions between β-sandwich domains, as reported for ArlF dimers [11].

Foldseek analysis of the mature YNH_2540 protein (excluding the signal peptide) identified 297 structural hits in the PDB100 database (Supplementary Information Fig. 15). Of these, the only structurally similar protein from Archaea was the bundling pili protein (AbpA) from *Pyrobaculum calidifontis* (PDB ID: 7UEG), whose structure was resolved by cryo-electron microscopy (cryo-EM) [14]. Despite the low sequence identity (12.7%), the alignment spanned nearly the entire protein (residues 5–135 of YNH_2540 and 3–180 of AbpA), with a high probability score (1.0) and the lowest E-value ($5.0 \times 10^{-4}$) among all hits. Structural superposition further revealed a close resemblance between the β-sandwich fold of YNH_2540 and that of AbpA from *Pyrobaculum calidifontis* (Supplementary Information Fig. 14g). AbpA polymerizes into a helical filament through donor-strand exchange between adjacent monomers [14]. To examine whether YNH_2540 could adopt a similar assembly mode, AlphaFold multimer predictions were performed using six identical YNH_2540 sequences. The resulting model showed that the monomers were arranged in a continuous linear chain, with the N-terminal β-strand of each subunit extending into the β-sheet of the neighbouring monomer (Supplementary Information Fig. 14c), suggesting that YNH_2540 may also assemble through a donor-strand exchange mechanism to form a filament-like structure. These observations collectively suggest that YNH_2540 shares a structural and assembly principle with AbpA, potentially functioning as a filament-forming surface protein involved in host–symbiont interactions.

**Protein diversity in strains YN4, YN4HA, and MTIV4**

In the glycoproteome analysis of cultures containing strains YN4, YN4HA, and MTIV4, we analysed two culture samples using an N-glycosylation open search and identified 317 and 165 proteins from YN4, 1,122 and 1,046 proteins from YN4HA, and 6 and 10 proteins from MTIV4, respectively (Supplementary Tables 21 and 22). In the co-culture of strains YN4 and YN4HA, 177 proteins were identified from YN4 and 990 from YN4HA (Supplementary Table 20). These numbers represent approximately 19.7-37.9% of the total coding sequences (CDSs) of YN4, 35.1-39.8% of YN4HA, and 20.0-33.3% of MTIV4. Comparatively, our previous study on *Nanobdella aerobiophila* MJ1$^{T}$ across various growth phases identified between 289 and 309 proteins [15], suggesting that members of the order *Nanobdellales* typically express at least 150 proteins, with many cases near 300. In strain YN4, the following proteins were consistently ranked among the top 10 based on NSAF values across three independent samples: archaeal histones (YN4_409 and YN4_239), putative snRNP Sm-like protein (YN4_253), 50S ribosomal protein P1 (YN4_476), hypothetical protein (YN4_252), and elongation factor EF-1 subunit alpha (YN4_516). These proteins were among the most highly expressed in the transcriptome analysis of YN4 co-cultured with YN4HA (rank 4-76, Supplementary Table 17). Similarly, histones, 50S ribosomal proteins, and elongation factors were abundantly detected in the proteome of *N. aerobiophila* MJ1$^{T}$ [15]. Apparent S-layer-like proteins (YN4_391 and YN4_612), which are typically abundant in free-living archaea, were not detected in strain YN4, consistent with our previous analysis of *N. aerobiophila* MJ1$^{T}$ [15], although they were highly expressed in the transcriptome of YN4 (rank 1 and rank 17, Supplementary Table 17). The absence of such proteinaceous crystalline cell surface layers among the highly expressed proteins may be a common characteristic of the order *Nanobdellales*. Possibly, the lack of S-layer-like proteins in the glycoproteome was influenced by technical or biochemical factors, such as limited solubility or strong membrane association, which make them more difficult to extract and detect under the conditions used. In contrast, a highly abundant S-layer protein (SlaA, YNH_0338) was consistently detected in strain YN4HA across all samples (Supplementary Tables 20-22).

**Predominant N-glycan modification masses under different culture conditions**

While wide-tolerance open searching facilitates the detection of heterogeneous peptide modifications by *N*-linked glycans without predefined structures [15–17], this method often yields pseudo-positive scans with random modification masses, even after excluding low-confidence peptide MS/MS scans. Consequently, we created histograms of peptide modification values, binning the observed values in 0.01 Da increments (Supplementary Information Fig. 16). Examination of the modifications in the pure culture of strain YN4HA and in the co-culture of strains YN4HA and MTIV4 revealed two predominant modification values of 568.21 and 1305.42 Da (Supplementary Information Fig. 5a,b). In contrast, when strain YN4HA was co-cultured with strain YN4, the modification at 568.21 Da remained predominant, whereas the 1305.42 Da modification was absent. Additionally, new predominant modifications at 1159.36 and 794.23 Da emerged, suggesting a change in the glycan profile of strain YN4HA when co-cultured with strain YN4 (Supplementary Information Fig. 5c,d). In strain YN4, four predominant modification values of 1200.43, 1054.37, 1038.38, and 892.32 Da were observed, all exhibiting apparent sugar oxonium ions, despite low PSM counts and noisier histograms (Supplementary Information Fig. 5e). No predominant modifications were observed in strain MTIV4, suggesting the absence of glycan modification in the proteins of the virus (Supplementary Information Fig. 5f,g).

**Host-dependent glycan remodeling in strain YN4**

Similarly, when cultured with *Metallosphaera cuprina*, the major delta mass value of 1200.43 was no longer detected in strain YN4 (Supplementary Information Fig. 16). Major glycoproteins of strain YN4 detected in the co-culture with *M. cuprina* were the same as those detected in the co-culture with strain YN4HA. The MS/MS analysis of the predominant glycopeptide indicated a loss of dHex compared with the co-culture with strain YN4HA, suggesting the same glycan structure as reported in *N. aerobiophila* MJ1$^{T}$ [15] (Supplementary Information Fig. 17). This change is potentially attributable to the absence of dHex in *M. cuprina* [15], supporting our previous findings that *Nanobdellati* archaea acquire and alter host glycans to modify their own proteins. This probably represents a significant difference between *Nanobdellati* archaea and viruses [15].

**Proposal of *Nanomutabilis hospitiaugens* for strain YN4**

Na.no.mu.ta'bi.lis. Gr. masc. n. *nânos*, a dwarf; mutabilis. L. masc./fem. adj. *mutabilis*, changeable, referring to the narrow host specificity while being capable of utilizing at least two species as hosts.

hos.pi.ti.au.gens. L. neut. n. *hospitium*, a dwelling or host; L. neut. n. *augens*, enlarging or increasing; N.L. neut. n. *hospitiaugens*, a host-enlarging organism, referring to its ability to induce swelling in its host.
